# High Content Screening and Proteomic Analysis Identify a Kinase Inhibitor that rescues pathological phenotypes in a Patient-Derived Model of Parkinson’s Disease

**DOI:** 10.1101/2020.06.12.148031

**Authors:** Nasia Antoniou, Kanella Prodromidou, Georgia Kouroupi, Martina Samiotaki, George Panayotou, Maria Xilouri, Leonidas Stefanis, Regis Grailhe, Era Taoufik, Rebecca Matsas

## Abstract

Combining high throughput screening approaches with induced pluripotent stem cell (iPSC)-based disease modeling represents a promising unbiased strategy to identify therapies for neurodegenerative disorders. Here we applied high content imaging on iPSC-derived neurons from patients with familial Parkinson’s disease bearing the G209A (p.A53T) α-synuclein (αSyn) mutation and launched a screening campaign on a small kinase inhibitor library. We thus identified the multi-kinase inhibitor BX795 that at a single dose effectively restores disease-associated neurodegenerative phenotypes. Proteomics profiling mapped the molecular pathways underlying the protective effects of BX795, comprising a cohort of 118 protein-mediators of the core biological processes of RNA metabolism, protein synthesis, modification and clearance, and stress response, all linked to the mTORC1 signaling hub. In agreement, expression of human p.A53T-αSyn in neuronal cells affected key components of the mTORC1 pathway resulting in aberrant protein synthesis that was restored in the presence of BX795 with concurrent facilitation of autophagy. Taken together, we have identified a promising small molecule with neuroprotective actions as candidate therapeutic for PD and other protein conformational disorders.

## Introduction

Parkinson’s disease (PD) is a complex neurodegenerative disorder affecting 2% of the world population over 65 years of age (1). PD is characterized by motor dysfunction related to the progressive loss of midbrain dopamine neurons (2) while a wide range of non-motor symptoms are also present such as psychiatric manifestations and cognitive impairment (3). The neuropathological hallmark of PD is the presence of intracytoplasmic inclusions in neuronal cell bodies and neurites, respectively termed Lewy bodies and Lewy neurites (4, 5). These are protein aggregates composed mainly of α-synuclein (αSyn), the major protein linked to sporadic PD (6). αSyn belongs to a class of intrinsically disordered amyloid proteins that form specific forms of oligomeric and fibrillar aggregates and exert neurotoxicity through various molecular pathways (7). Several point mutations (A30P, E46K, A53T, G51D) and multiplications of the *SNCA* locus encoding for αSyn cause autosomal dominant forms of PD (8–10). Among the different variants, the p.A53TαSyn mutation is generally considered to accelerate aggregation (11) resulting in widespread accumulation of insoluble α-syn deposits that have been identified in the post-mortem p.A53T human brain (12, 13). Despite extensive efforts in understanding PD pathogenesis, no disease modifying drugs exist. Currently only symptomatic or palliative treatments are available with none capable to prevent or slow-down disease progression. Dopamine-replacement drugs, such as levodopa, which was identified 53 years ago (14), are used to ameliorate motor symptoms and remain the primary and most effective treatment despite the undesired side-effects and deterioration of efficacy with disease progression. Therefore, the development of disease-modifying drugs is an urgent unmet need. Most present-day efforts in identifying novel PD therapeutics target the aggregation of misfolded αSyn as the major pathogenic factor that causes cellular toxicity (6, 15–17). Alternative strategies to tackle early steps in neurodegeneration, particularly in an unbiased approach, have lagged behind. Recent advances in patient-derived induced pluripotent stem cell (iPSC)-based models for neurodegenerative diseases permit the detection of early, potentially triggering, pathologic phenotypes and provide amenable systems for drug discovery. In combination with high throughput high content screening technologies, these approaches open new perspectives for identification of disease-modifying compounds (18–21).

We have previously established a model of iPSC-derived neurons from patients with familial PD harboring the p.A53T αSyn mutation (G209A in the *SNCA* gene) that displays disease-relevant phenotypes at basal conditions (22).In this study, we successfully adapted this cellular system to perform the first small molecule screen on human p.A53T-neurons and discovered that the multi-kinase inhibitor BX795 significantly reverts disease-associated phenotypes. A single treatment of patient neurons with BX795 has sustainable effects in supporting neuritic growth, restoring axonal pathology and limiting αSyn protein aggregate formation. Protection from p.A53T-associated pathology was also confirmed in human iPSC-derived neurons in which the mutation was introduced by genome editing, against isogenic wild-type controls. Strikingly, proteomics profiling by quantitative mass spectrometry revealed that BX795 treatment results in significant downregulation of a cohort of 118 proteins that are abnormally upregulated in p.A53T-neurons. Enrichment analysis demonstrated that these proteins are associated with mRNA metabolism, mRNA transport and translation, protein metabolism and degradation processes. Using neuronal cells expressing the human p.A53T-αSyn, we demonstrate that BX795 affects the mTORC1 pathway to restrict excessive protein synthesis and facilitate autophagy. Taken together, our data highlight the BX795 kinase inhibitor as a compelling compound and candidate therapeutic that ameliorates p.A53T-related pathology.

## Results

### Assay development for high-content screening of p.A53T-iPSC derived neurons

iPSCs used in this study were previously generated from a PD patient bearing the p.A53T αSyn mutation and thoroughly characterized (22). For directed differentiation a dual SMAD inhibition protocol was used in the presence of Noggin and TGFβ inhibitor (22–24), which favors the generation and expansion of Pax6+/Nestin+ neural progenitor cells (NPCs; Fig. 1a). NPCs were further differentiated into βIII-tubulin (TUJ1)+ neurons (Fig. 1a) with 15-20% also expressing the dopaminergic marker TH at 21 DIV (Fig. 1a). The expression of dopaminergic lineage markers, such as Nurr1, TH, and aromatic amino acid decarboxylase (AADC) was confirmed by qRT-PCR (Fig. S1a). As readout for compound screening, we assessed TH immunofluorescence in iPSC-derived neurons adapted in miniature 384-well plates, seeking to identify putative neuroprotective compounds enhancing dopaminergic neuron output. To this end, the fluorescent signal for TH within a well was normalized to the fluorescent signal for the pan-neuronal marker βIII-tubulin (TUJ1) (Fig. 1b).

**Fig 1.**
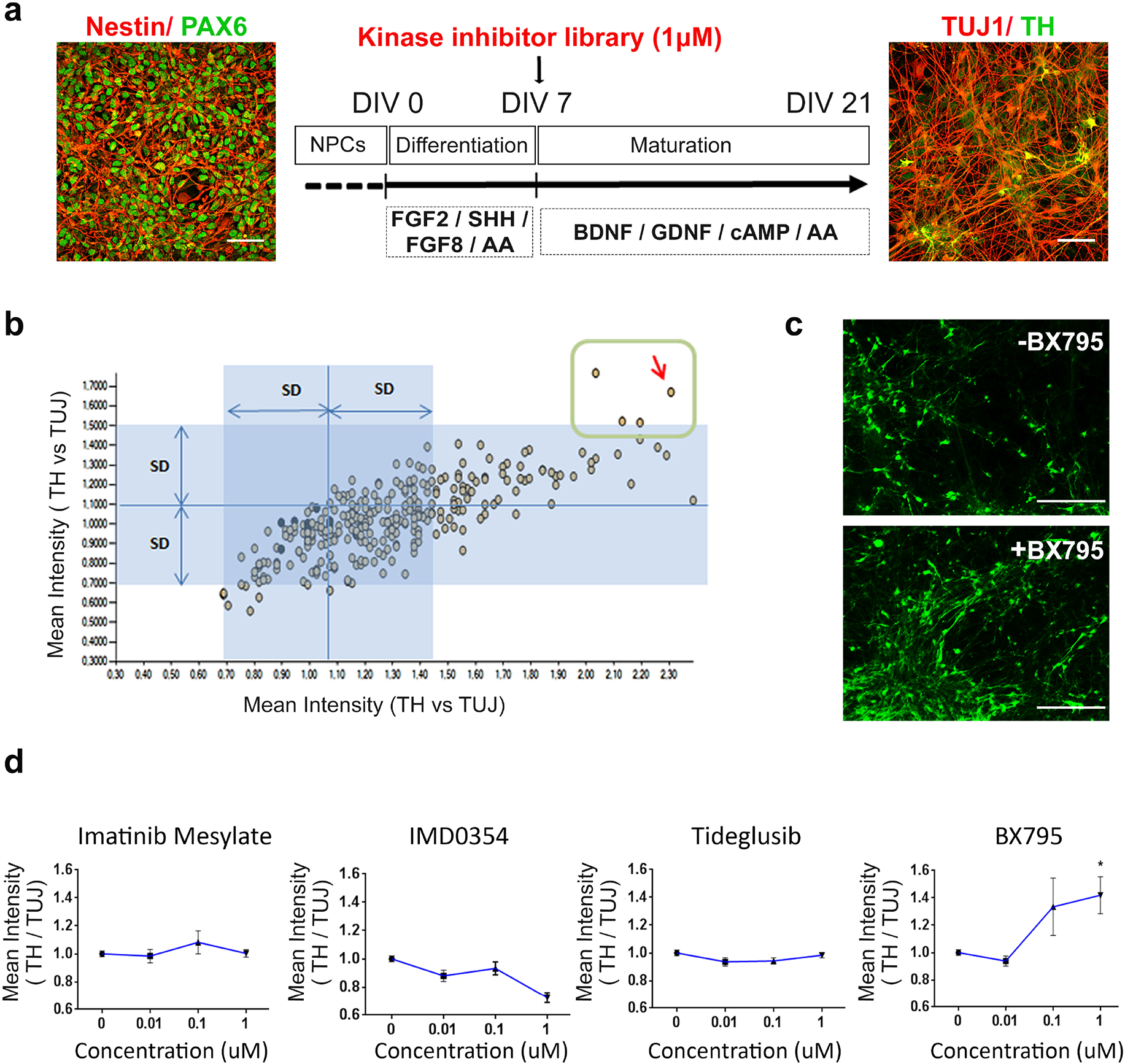
Identification of BX795 by high content screening of a kinase inhibitor library. a. Directed differentiation of Pax6+ (green)/ Nestin+ (red) neural precursor cells (NPCs; DIV 0, left) into TUJ1+ (red)/ TH+ (green) neurons (DIV 21, right). The differentiation protocol and timeline of analysis are shown in the drawing in the middle.. FG2 and FGF8, fibroblast growth factors 2 and 8; SHH, Sonic Hedgehog; AA, ascorbic acid; Scale bar represents BDNF, brain-derived neurotrophic factor; GDNF, glial cell-derived neurotrophic factor (GDNF); cAMP, cyclic AMP. Scale bars, 50 μm. b. Scatter plot showing the ratio of TH versus TUJ1 fluorescence intensity in duplicate upon treatment with 273 small molecule kinase inhibitors. The dots inside the green square correspond to the 4 hit compounds showing significant increase of TH versus TUJ1 fluorescence ratio as compared to the DMSO controls (blue dots). The red arrow indicates BX795. c. Representative images of patient-derived p.A53T-neurons immunolabelled for TH in 384-well plates. Upper micrograph shows control DMSO-treated cells while lower micrograph represents BX795-treated cells. Scale bar represents 150 μm. d. Tests of the four hit compounds in a dose-response format. Data are presented as mean ± SEM.(one-way ANOVA, *P<0.05, n=3 independent experiments).

### High content screening of a kinase inhibitor library identifies BX795 as a compound that increases TH immunofluorescence in p.A53T-neurons

Protein kinases represent central molecular hubs that regulate numerous cell processes, thus constituting potentially attractive clinical targets. Indeed, the success of kinase inhibitors in treating cancer has spurred the evaluation of such compounds in phase II/III clinical trials as candidates for treatment of various neurodegenerative diseases (25, 26). Since several kinases have been implicated in PD pathology (27), we screened a collection of 273 small molecule kinase inhibitors (Table S1) to identify compounds with prospective neuroprotective properties. p.A53T cells were exposed once (7 DIV) to the library of kinase inhibitors at 1μM concentration and quantitative image analysis was performed at 21 DIV (Fig. 1a). Hits were defined as compounds that robustly conferred an increase in TH immunofluorescence compared to DMSO-treated p.A53T neurons within a well, normalized to the immunofluorescence of the pan-neuronal marker βIII-tubulin (TUJ1) (Fig. 1b). Toxic compounds were excluded by assessing cellular viability (total nuclei count) of compound-treated as compared to DMSO-treated cells (Fig. S2). Four hits were identified in the primary screen (Fig. 1b), which were re-tested for validation in a dose-response assay (Fig. 1d). Of these BX795, an aminopyrimidine compound that acts as a multi-kinase inhibitor with pro-survival and/or anti-inflammatory effects (28), significantly increased TH immunofluorescence at 1 μM (Figs. 1c, d). BX-795 was initially developed as an ATP-competitive inhibitor of 3-phosphoinositide-dependent kinase 1 (PDK1), but was later shown to inhibit the IKK-related kinase, TANK-binding kinase 1 (TBK1) and IKKε, as well as to have numerous additional targets (29–31). Based on the sustained effect of a single dose of BX795 on p.A53T dopaminergic neurons (Fig. 1d), we focused further on this compound to explore its function.

### BX795 rescues neuropathological features of p.A53T neurons

The effects of BX795 on p.A53T-neurons were tested in cells that received a single treatment of the kinase inhibitor (1 μM) at 7 DIV and were analyzed at 21 DIV, in accordance with the protocol applied during the screening procedure. Prior to this, an initial set of experiments was performed using drug concentrations from 0.1-2 μM and repeated drug additions every 3 days, with the selected scheme ensuring optimal efficacy and minimal toxicity. Initially, we asked if the enhancement in TH immunofluorescence could be attributed to an increase in cell survival/proliferation or dopaminergic differentiation in p.A53T-cultures. We could not detect BX795-driven changes in either proliferation, as assessed by the percentage of Ki67+ cells (Fig. S1b; % Ki67+ cells, DMSO: 43.3 ± 4.4; BX795: 50.3 ± 1.5, n=3), or in differentiation as estimated by the percentage of TH+ cells in the culture (% TH+ / TUJ1+ neurons, DMSO: 13.9 ± 3.1; BX795: 18.1.0 ± 3.9, n=3). These observations indicate that the effect of BX795 on dopaminergic neurons is not related to an increase in either survival/proliferation or differentiation.

Next, we investigated if treatment with BX795 could rescue neuropathological features previously identified in p.A53T-neurons, such as compromised neuritic growth, dystrophic or fragmented neurites and the presence of intracellular protein aggregates (22, 32). Overall, disease-associated phenotypes were assessed in iPSC-derived neurons from two p.A53T patients [22] and an iPSC gene-edited line in which the p.A53T mutation was inserted in one allele, against healthy or isogenic controls. Evaluation of total neurite length in TH+ dopaminergic neurons from the first p.A53T patient revealed a significant increase in response to BX795 compatible with the observed increase in TH immunofluorescence (length in μm, ctl: 221.7 ± 16.8, p.A53T:127.2 ± 13.5, p.A53T+ BX795: 196.8 ± 21.1, n=5, Fig.2a). Moreover, examination of the distinct pathological morphology of TUJ1+ p.A53T neurons revealed an almost 50% reduction in axonal degeneration (axon degeneration index: ctl: 2.945 ± 1.325, p.A53T:13.03 ± 1.491, p.A53T+ BX795: 7.276 ± 1.017 n=3; Fig. 2b). Finally, exposure to BX795 resulted in a notable 60% decrease in protein aggregate formation in p.A53T cells (number of aggregates per cell, p.A53T: 8.431 ± 0.77, n= 51, p.A53T+ BX795: 3.242 ± 0.40, n=62; Fig. 2c). This was accompanied by a consistent decline in the levels of (Ser129)-phosphorylated αSyn (Fig. 2d), a modification that renders αSyn prone to self-assembly and is commonly associated with synucleinopathy (33, 34). The neuroprotective effects of BX795 were confirmed in p.A53T-neurons from a second patient (22, 32) (Fig. S3).

**Fig 2.**
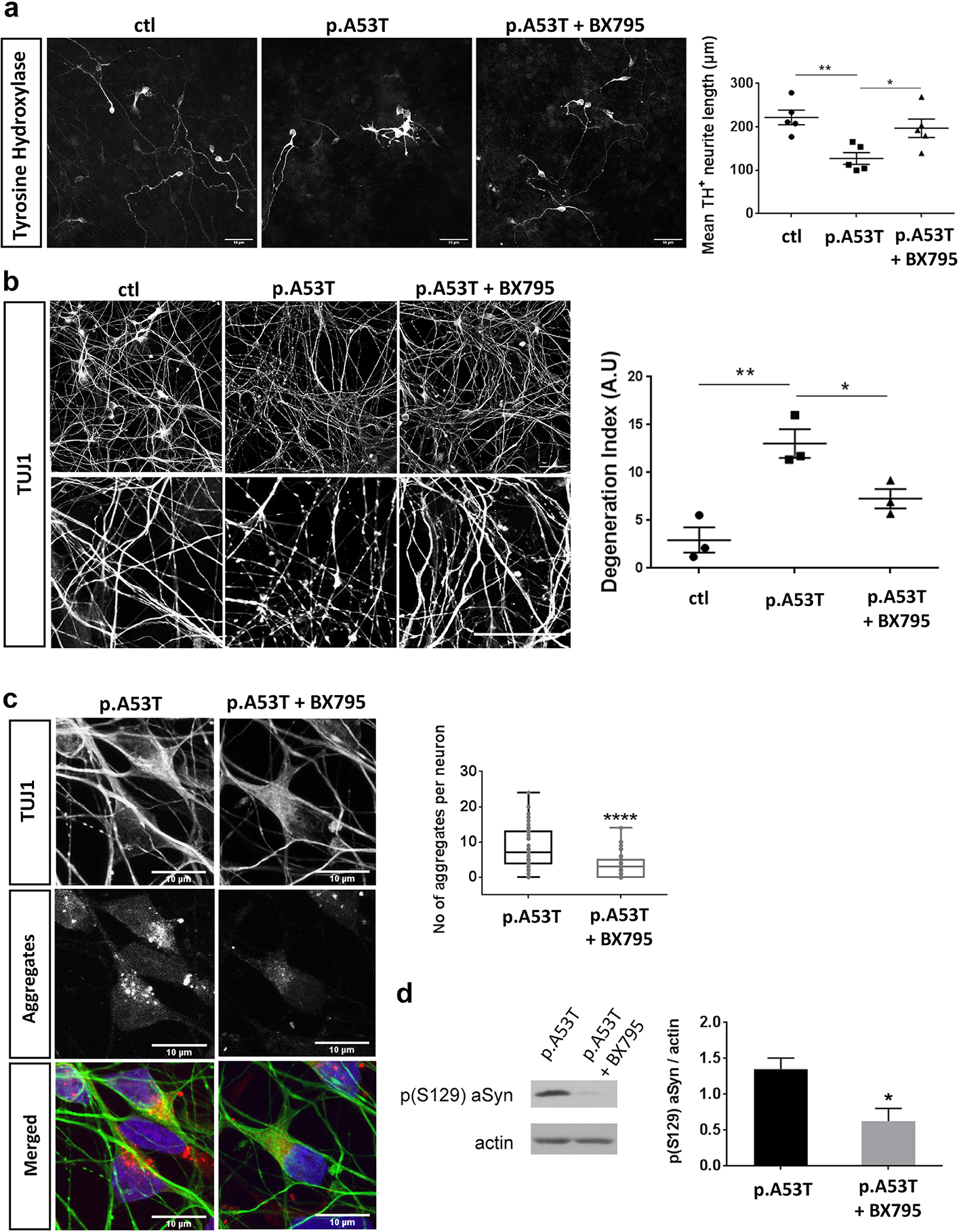
Rescue of neuropathological features in patient-derived p.A53T neurons by BX795. a. BX795 has a positive effect on neurite length of p.A53T-neurons. Representative confocal images of healthy control (ctl) and p.A53T-neurons immunostained for TH and quantification of total neurite length of TH+ cells. Data represent mean ± SEM. (Comparisons by ANOVA with Tukey correction *P<0.05, **P<0.01, n=4 independent experiments with at least 50 cells analyzed in each experiment). Scale bar, 50μm. b. BX795 alleviates axonal neuropathology in p.A53T-neurons. Higher magnification at the right (upper, DMSO-treated cells; lower, BX795-treated cells) shows neurites with swollen varicosities or fragmented processes (arrows). Scale bar, 30μm. Quantification of axonal degeneration is estimated in the accompanying graph by measuring the ratio of TUJ1+ spots over the total TUJ1+ area in untreated (p.A53T) or BX795-treated p.A53T-neurons. Data represent mean ± SEM.(Comparisons by ANOVA with Tukey correction, *P< 0.05, **P<0.01, n = 20 randomly selected fields for each condition). c. BX795 reduces protein aggregates in p.A53T-neurons. Representative confocal images showing protein aggregates in p.A53T TUJ1+ neurons (Scale bar, 10μm) and quantification in untreated or BX795-treated TUJ1+ cells (Mann–Whitney test; n=at least 50 cells per group; ****P< 0.0001). d. Detection and quantification of p(Ser129)αSyn by Western blot; Actin shows equal protein loading. Data represent mean ± SEM (*t*-test, *P<0.05, n=4 independent experiments).

We also assessed the neuroprotective effects of BX795 in a highly enriched culture of mature human midbrain dopaminergic neurons (Fujifilm Cellular Dynamics Inc). These comprised an isogenic pair of wild-type (iCell DOPA) and gene-edited (iCell A53T DOPA) iPSC-derived neurons in which a heterozygous p.A53T mutation was inserted into one allele of the *SNCA* gene. After 14 days, more than 90% of cells were TUJ1+ and more than 80% were TH+ dopaminergic neurons (Fig3a). At this time and similarly to patient-derived cells, abundant protein aggregates were detected in the p.A53T iCell neurons compared to their isogenic control, and treatment with BX795 resulted in a significant reduction (number of aggregates per cell, ctl:2.7±0.49,n=57, p.A53T: 9.9 ± 1.1, n=76, p.A53T+ BX795:4.9 ±0.7, n=76; Fig3b,c). Taken together our results indicate that BX795 exerts prominent and sustainable neuroprotection in p.A53T neurons by improving neuritic growth, limiting the levels of pathological αSyn and restricting aggregate formation whilst maintaining axonal integrity. The beneficial effects of BX795 were noted whether it was added early during neuronal differentiation or at later stages of neuronal maturation when disease-associated phenotypes were was already established.

**Fig 3.**
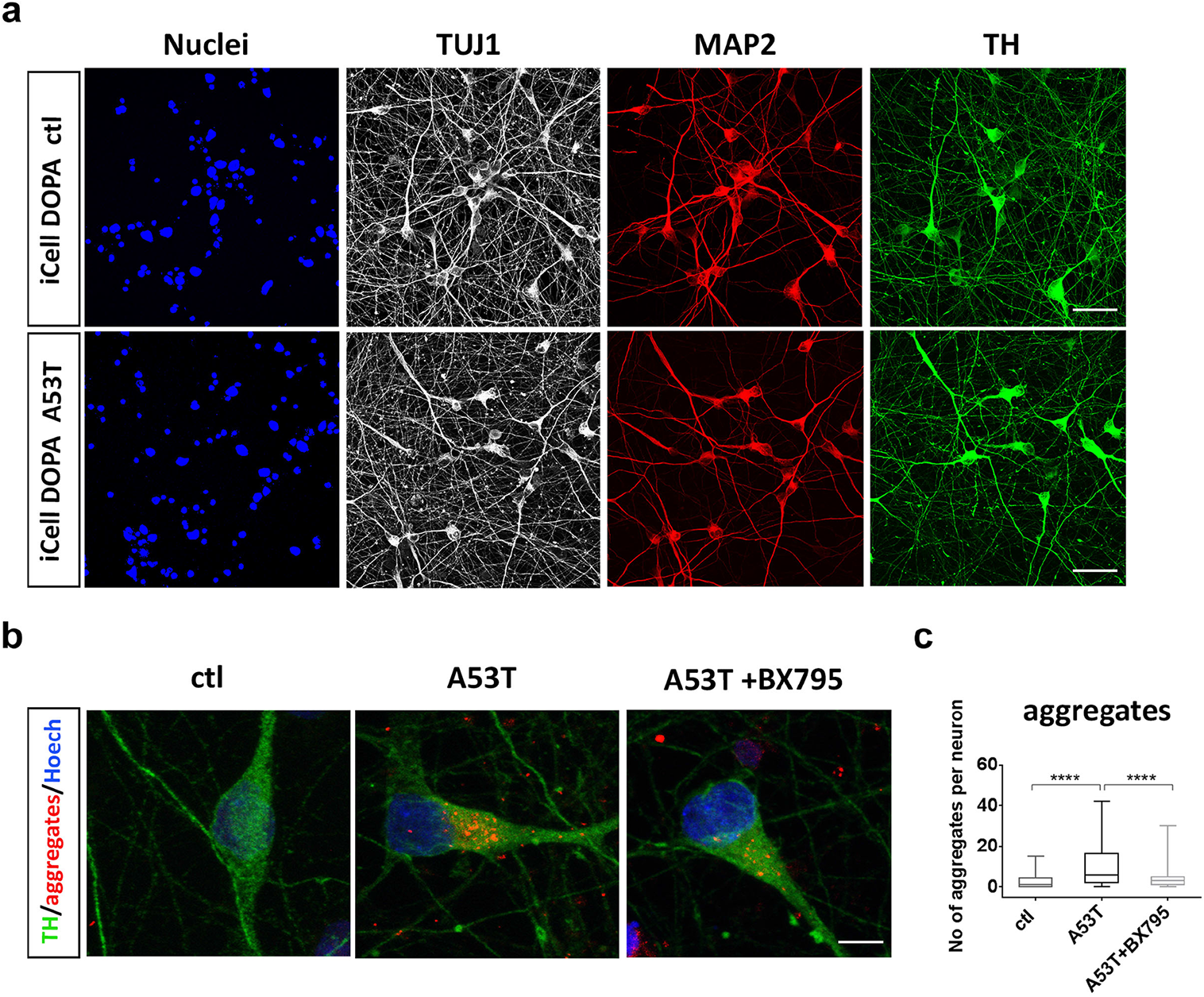
BX795 reduces protein aggregates in a gene-edited p.A53T line of mature human iPSC-derived TH neurons. a. Representative confocal images of wild-type (ctl) and isogenic p.A53T iCell Dopa neurons immunolabelled for Nuclei, TUJ1, MAP2 and TH. Scale bar, 30 μm b. Representative confocal images of wild-type (ctl) and isogenic p.A53T iCellDopa neurons showing immunostaining for tyrosine hydroxylase (TH green) and protein aggregates (red). p.A53T cells were treated or not with BX795, as indicated. Scale bar, 5μm c. Quantification of aggregates in TH+ neurons. Data represent mean ± SEM. (Comparisons by ANOVA with Tukey correction, ****P<0.0001, n = at least 50 randomly selected TH+ cells for each condition).

### Proteomics analysis identifies cellular pathways targeted by BX795 in p.A53T neurons

Identification of the BX795 affected cellular pathways which vary according to the system investigated (30, 31, 35), is a challenging task. Therefore, we used an unbiased approach based on comparative proteomics. Similarly to the screening procedure, BX795 was added once at DIV 7 and proteomics analysis was performed at DIV 21 when rescue of neuropathological phenotypes was noted (Fig. 2). A total of 1652 proteins were identified and quantified using the MaxQuant software (36, 37), followed by filtering of low quality protein hits with the Perseus software. Initial comparison between p.A53T versus control neurons in the absence of BX795, revealed differential expression of 640 proteins (Fig. S4a) from which only 67 were down-regulated whilst the rest 573 were up-regulated (Fig. S4b, Table S2). This large increase in protein expression was linked by GO enrichment analysis mainly to the biological processes of transcription, translation, protein synthesis and modification (Fig. S4b). Remarkably, the levels of a cohort of 118 proteins lying mostly within these biological processes and representing approximately 20% of the total dysregulated proteins in p.A53T neurons, were restored upon treatment with BX795 (p<0.05) (Fig. 4a, Table S3). Most important, this outcome was specific to p.A53T-neurons as BX795 had no significant effect on the proteome of control neurons.

**Fig 4.**
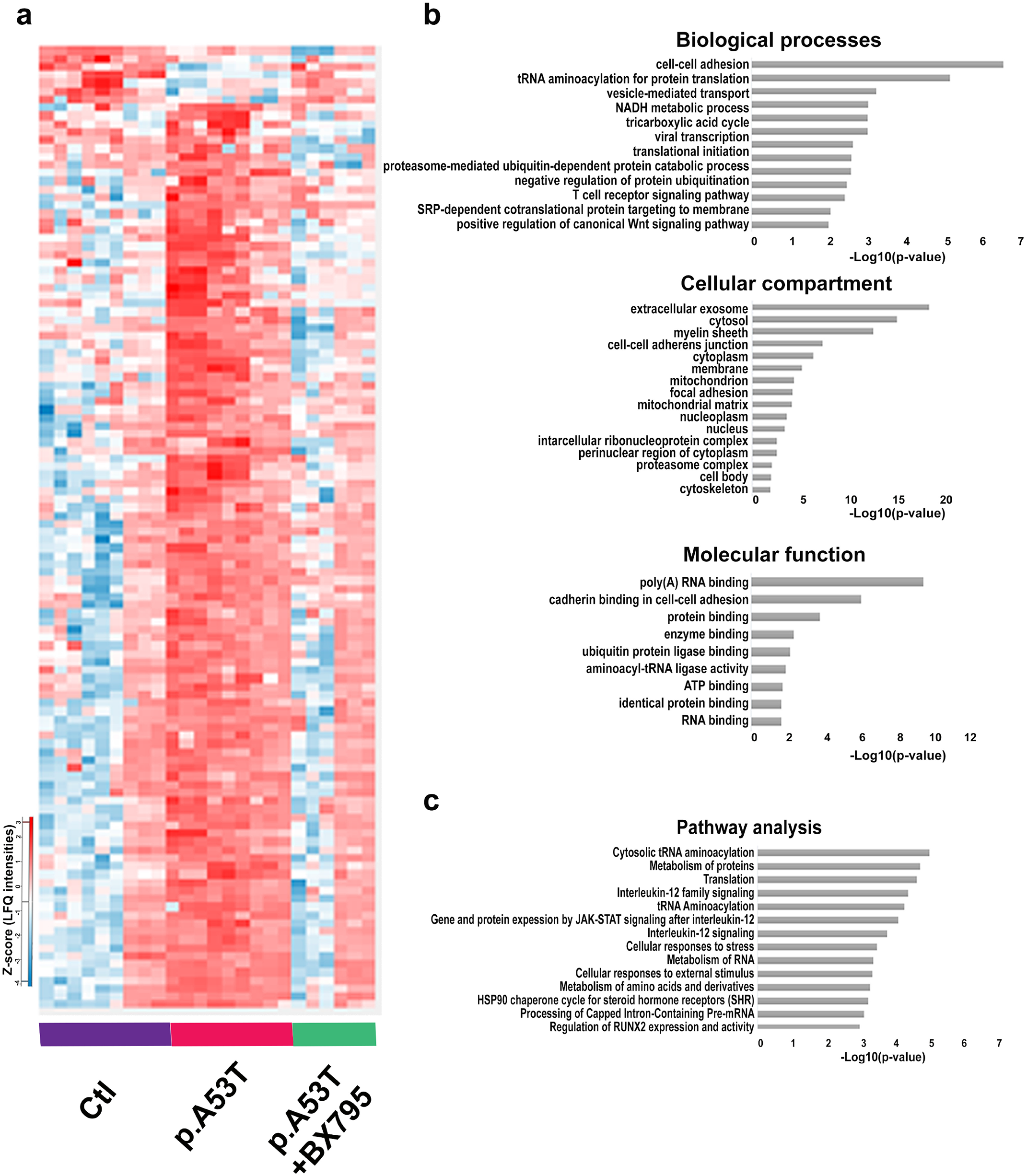
Bioinformatics analysis of dysregulated proteins in p.A53T-neurons that are restored by BX795. a. Hierarchical clustering of 118 upregulated proteins in patient-derived p.A53T-neurons that are restored upon treatment with BX795 (one-way ANOVA analysis). Columns in the different groups (control, p.A53T-neurons and p.A53T-neurons treated with BX795) correspond to individual samples tested and rows represent single proteins (blue, low expression; red, high expression; n=3 for control and p.A53T; n=2 for p.A53T+BX795). b. GO enrichment analysis for biological processes, molecular function and cellular compartments was performed using DAVID software (p<0.01). c. Pathway analysis using Reactome software (p<0.01)

Extensive data mining by GO enrichment analysis for biological processes, molecular function and cellular compartments (p<0.01), complemented by reactome pathway analysis (p<0.01), highlighted the dysregulated core pathways in p.A53T-neurons and, amongst them, those targeted by BX795 to restore neuronal physiology (Fig. 4b). These include proteins associated with RNA metabolism, protein synthesis, protein modification and transport, stress response, and neurodegeneration, as outlined below.

#### RNA metabolism

The p.A53T proteome showed enrichment for proteins in subcellular compartments known to be associated with αSyn (38), including membrane bound organelles (204 proteins), mitochondria (118), ribosomal complexes (29), nucleus (292), and neuron projection/axon cytoplasm (10) (Table S4). Processes such as cellular metabolism, translational initiation and regulation, tRNA aminoacetylation and export from nucleus, mRNA stability and export from nucleus, rRNA processing, formation of pre-initiation complex and protein folding were among the top pathways enriched in the p.A53T proteome (Fig. S4). A previous study has identified mRNA binding proteins (RBPs) and those involved in protein biosynthesis within the protein network residing in immediate vicinity of αSyn, suggesting that perturbation of these pathways may be directly related to pathology (38). Herein, we provide evidence that these same pathways are altered when p.A53T is expressed in human neurons (Fig. S4). Specifically, a significant number of RBPs (60 proteins) were differentially expressed, including members with known neuronal localization and involvement in neuronal functions, such as ELAV-1, ELAV-3, RBBP7, RNPS1, RNMT, TARDBP, XPO1, XPO5, HNRNPA1, HNRNPA1L2, HNRNPF, HNRNPL, HNRPNPM, HRNNPUL1, PABPC1, PABPC4, PTBP2 and CELF1 (Table S2). Since even small changes in RBP expression or activity are amplified due to their broad impact on expression, splicing and translation of numerous RNA substrates, changes in such a large number of these RNA regulators suggest a severe perturbation in RNA homeostasis in p.A53T-neurons. A cluster of RBPs implicated in splicing and adenylation events in the nuclear compartment (DEK, MYEF2, UBTF, SNRPB, PCBP1, ZNF207, HINT1, RAE1, HNRNPUL1) was restored after BX795 treatment (Fig. 5a).

**Fig 5.**
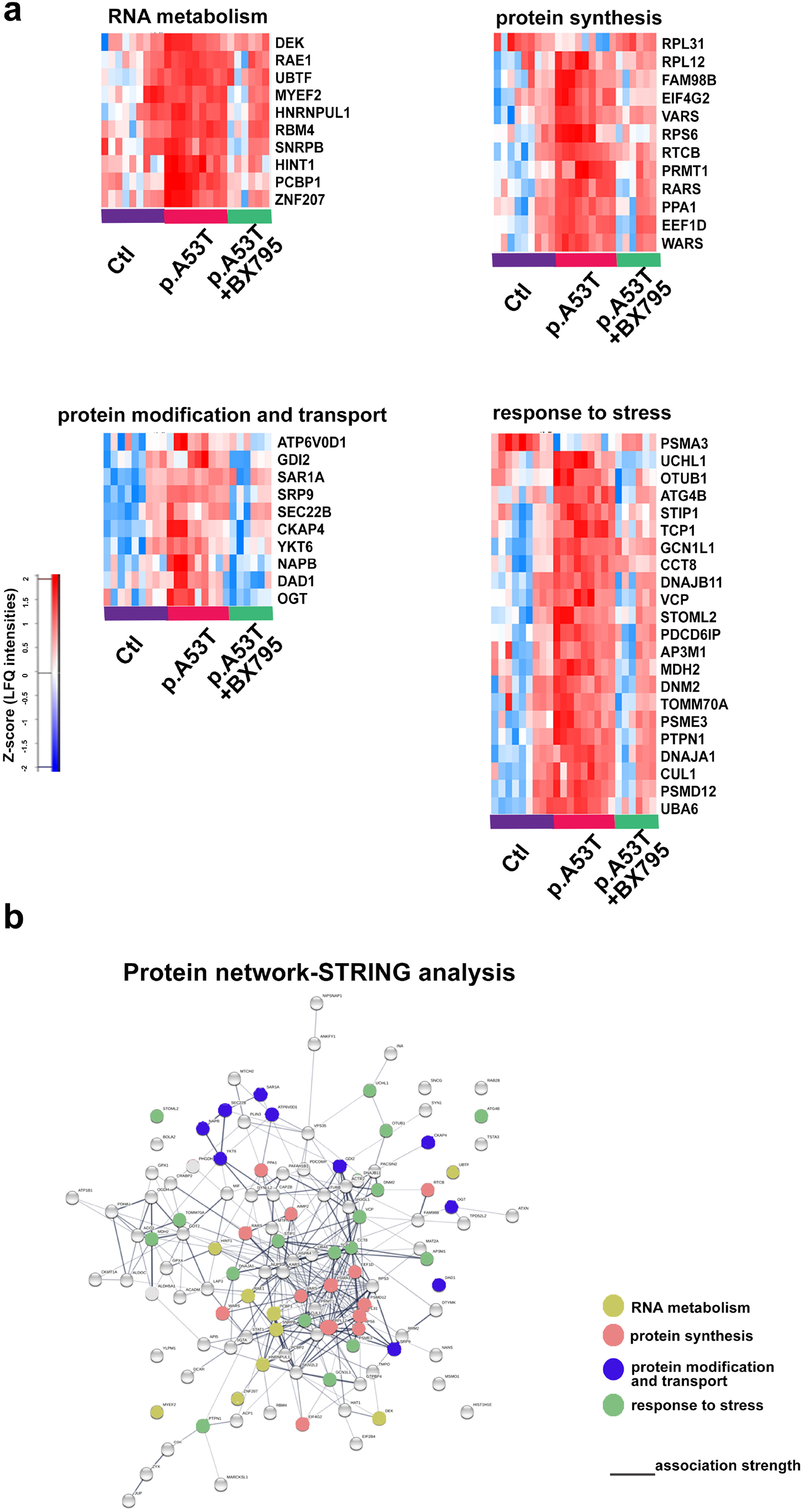
Protein network of pathways and processes restored by BX795 treatment. a. Heatmaps illustrating specific proteins upregulated in p.A53T-neurons that are involved in RNA metabolism, protein synthesis, protein modification and transport and response to stress, which are restored after BX795 treatment. High expression is in red and low expression is in blue. b. STRING-analysis representation of the protein-protein interaction network of the 118 upregulated proteins in p.A53T-neurons that are restored by BX795. Each circular node depicts one protein and the different colors represent the different pathways/processes as indicated. Connecting lines represent protein-protein associations and line intensity represents the confidence score of a functional association.

#### Protein Synthesis

Disturbances in RBP dosage have detrimental consequences also outside the nucleus, as they control the targeted localization of mRNAs, either proximally in the cell soma or distally in the projecting axon, affecting whether an mRNA will be translated or remain translationally silent and whether it will be stored for local mRNA translation or degraded (39). Aberrant expression of the translational machinery emerged in the p.A53T proteome with translational initiation and regulation processes being the most affected in mutant neurons (Fig. S4b, Table S2). A total of 18 proteins involved in the formation of the pre-initiation complex were identified and included EIF2, 3, 4 and 5, of which EIF4G2 that functions as a general suppressor of translation by forming translationally inactive stress granules, was targeted by BX795 (Fig. 5a). Ribosomal proteins (29 proteins), structural components of ribosome subunits, were upregulated in p.A53T-neurons (Table S2) and a significant fraction returned to near-control levels after BX795 treatment (Fig. 5a). These included RPL31 and RPL12, which are involved in 60S biogenesis, and RPS6, a component of the 40S subunit and downstream effector of the mTORC1 signaling pathway. tRNA processing represents another important part of the translational cascade that was altered in p.A53T-neurons (Table S2), while a significant fraction was restored by BX795, including the aminoacyl-tRNA synthetases RARS (arginyl-tRNA synthase), VARS (valyl-tRNA synthase), and WARS (tryptophanyl-tRNA synthase) together with regulatory or accessory proteins such as PPA1, EEF1D PRMPT1, FAM98B and RTCB. Growing evidence associates changes in tRNA gene biogenesis and processing with neurodegenerative diseases (40). Our data reveal for the first time a link between p.A53T-αSyn expression and this molecular process (Fig. 5a, Table S2).

#### Protein modification and transport

p.A53T-αSyn toxicity has been attributed to problematic modifications at the ER membrane and disturbances in ER-Golgi and early endosomal/vesicle trafficking (33, 38, 41, 42). In accordance, p.A53T-neurons exhibit altered protein levels in components of these pathways (Table S2). Among these, five members of the adaptor protein complexes that function in transport-vesicle mediated transfer between membranous structures are increased by p.A53T-expression (AP1B1, AP2A2, AP3B1, AP3D1 and AP3M1). Another prominent category included members of the largest branch of Ras-like small GTPases involved in membrane trafficking (RAB2A, RAB2B, RAB6B). In addition, proteins participating in ER to Golgi transport and macroautophagy (SEC22B, SEC31A, RAB18, ARF1, ARF3) (43, 44), vesicle budding/uncoating in the Golgi apparatus (ARF1,ARF3) (45), SNARE-mediated autophagosome-lysosome fusion (RAB21) (46), retrograde Golgi to ER transport (COPA, COPB, COPG) (Table S2) were also differentially expressed in p.A53T neurons.

BX795 had a selective effect on p.A53T-altered membrane transport proteins (SRP9, GDI2, ATP6VOD1, DAD1 subunit of oligosaccahryl transferase complex and OGT, and NAPB) and components of the SNARE complex (SAR1A, SEC22B and YKT6) (Fig. 5a) whilst alterations on molecules of the RAB, adaptor protein complex and coatomer remained largely unaffected. ***Stress Response.*** p.A53T-αSyn protein expression acts as a primary neurotoxin triggering a battery of stress responses in human neurons (47). The proteomics analysis indicated that p.A53T neurons activate most of these mechanisms. Both the unfolded protein response (UPR), as evidenced by mis-expression of chaperones CCT2, 3, 4, 5, 7 and 8, as well as the heat shock protein response (HSP), with proteins such as DNAJA1, DNAJB11, DNAJC7, HSPA4L, HSP9 and HSPE1, were apparent in the p.A53T-proteome (Table S2). These stress response pathways were significantly downregulated in p.A53T neurons treated with BX795, which seems to target many stress response mediators (Fig. 5a). These included TCP-1, a member of the chaperonin TCP1 complex (CCT), PTPN1, a UPR regulator, STIP1, a coordinator of HSP70 and HSP90 function and the chaperone/ co-chaperone proteins DNAJB11, GCN1L1, CCT8, and DNAJA1.

Such a dysregulation of the UPR/HSP response systems in p.A53T neurons should result in the production of dangerous protein cargo and the formation of protein aggregates, as indeed identified by immunofluorescence (Fig. 2c). The p.A53T proteome also revealed alterations in protein clearance pathways with mediators of both proteasomal and autophagic systems being affected (Table S2). BX795 improved the expression of multiple ubiquitin-associated proteins suggesting partial restoration of proteasome targeting of aberrant protein products, in accordance with the decrease of protein aggregates in BX795-treated p.A53T neurons (Fig. 2c). BX795 restored the expression of components of the proteasome complex and activators of the E2 and E3 ligase binding process (PSMA3, UCHL1, OTUB1, PSME3, CUL1, PSMD12 and UBA6), and VCP, an AAA ATPase that extracts ubiquitinated proteins from large protein complexes for degradation, previously shown to co-localize with protein aggregates in various neurodegenerative diseases (Fig. 5a).

Components of the lysosomal pathway of autophagy targeted by BX795 included vacuole transport components such as ATG4B and proteins required for multivesicular body (MVB) biogenesis and sorting (PDCD6IP, AP3M1 and DNM2) (Fig. 5a). Finally, BX795 also modulated oxidative stress response mechanisms, as the mitochondrial biosynthesis regulators TOMM70A and MDH2 were brought to near control levels. In addition, STOML2, a stimulator of cardiolipin biosynthesis recently shown to be associated with p.A53T neurotoxicity in human dopamine neurons was also positively targeted by BX795 (33). When STRING analysis was used to assess the relatedness level of all 118 proteins affected by BX795, a network with strong functional linkage among the majority of these proteins was revealed (Fig. 5b).

#### Proteins associated with neurodegeneration

An important measure of the biological significance of the proteomic profile of p.A53T neurons comes from comparisons with human genetic studies. Enrichment analysis for PD and other neurodegenerative diseases identified several proteins comprising both known and novel converging targets that were modified by BX795 (Fig. 6a). Among those, UCHL1/PARK5 is linked to lower susceptibility for PD, while a point mutation co-segregating with the disease has been identified in one family (48) and VPS35/PARK17-D620N mutated protein causes late-onset autosomal dominant PD (49). FAM98B has been linked to SMA and ALS (50), VCP mutations can cause FTD, ALS and Charcot-Marie-Tooth diseases (51, 52), HINT1 autosomal recessive mutations lead to neuromytotonia and axonal neuropathy (53), PAFAHB1 mutations and gene deletions lead to lissencephaly syndrome (54) and RBM4 is linked to Down’s syndrome (55) (Fig. 6a, b). STRING analysis of the BX795-modified protein network to which αSyn was also incorporated, demonstrated a strong association between αSyn and other neurodegeneration-linked proteins (Fig. 6c).

**Fig 6.**
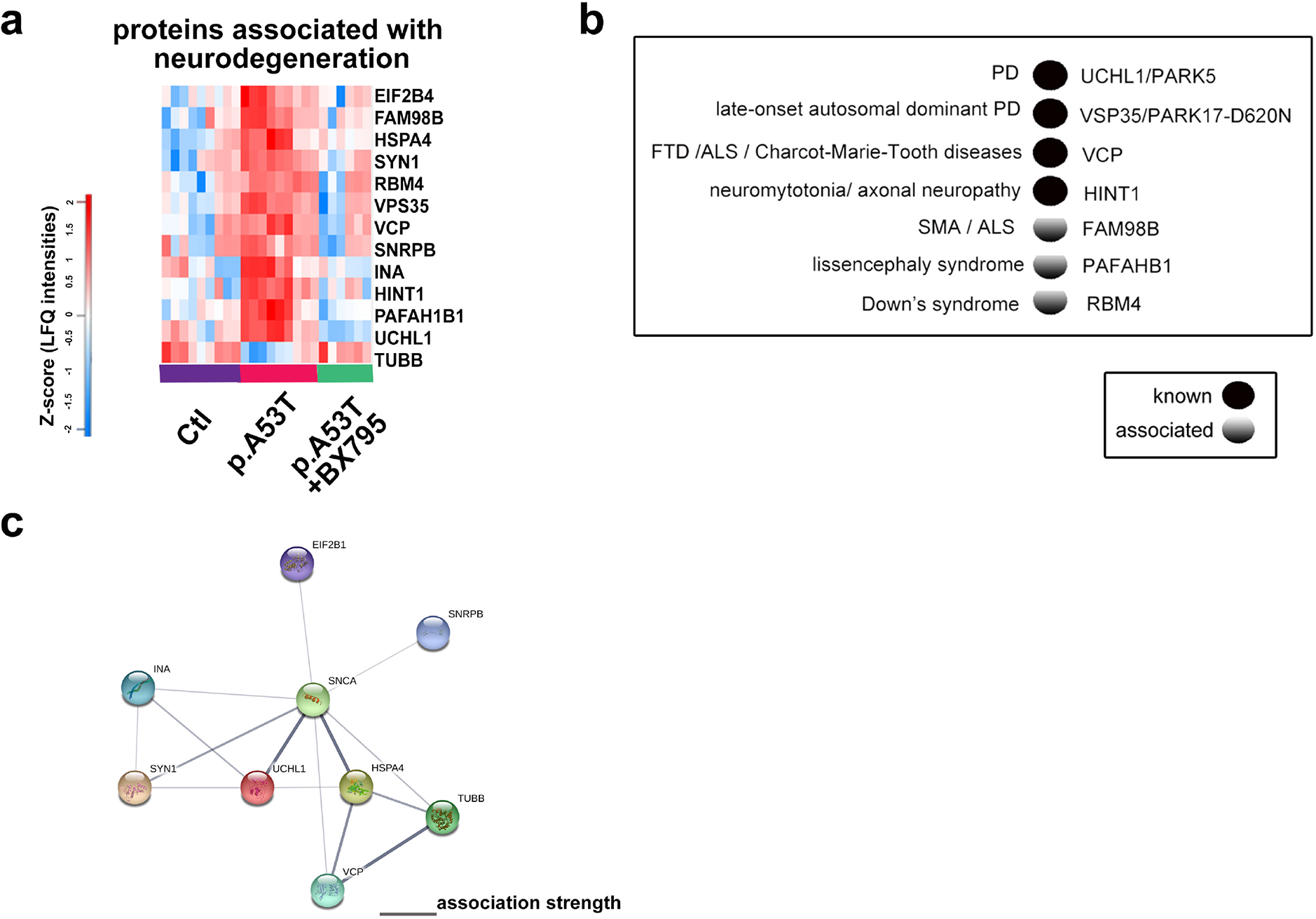
Restoration of disease-associated proteins by BX795 in p.A53T-neurons. a. Heatmap of proteins associated with neurodegeneration that are restored after BX795 treatment. High expression is in red and low expression is in blue. b. Disease-associated proteins that are modified by BX795 are either known or associated genetic risk factors for neurodegenerative diseases as revealed by human genetic studies. c. STRING network analysis of the neurodegeneration-associated proteins restored by BX795 in p.A53T-neurons and their interaction with αSyn. Each αSyn interactor is shown as a colored circle and connecting lines between proteins represent protein-protein associations. The intensity of lines represents the confidence score of a functional association.

These findings deepen our understanding of p.A53T-mediated neurotoxicity and reveal key biological processes that are targeted by BX795 to alleviate p.A53T-αSyn-related phenotypes in human neurons.

### BX795 affects the mTORC1 signaling pathway to attenuate protein synthesis and facilitate autophagic flux in p.A53T neurons

The p.A53T proteome clearly indicates aberrant mRNA translation and protein clearance mechanisms, both linked to mammalian aging and neurodegenerative diseases that can be effectively restored by BX795. The mammalian target of rapamycin (mTOR) signaling pathway is a central regulator of proteostasis and the p.A53T proteome clearly indicates hyperfunctional overactive biosynthetic processes that could be associated with alterations in mTORC1 activation. Components of this signaling cascade have emerged in the proteomics analysis of p.A53T-neurons, including RPS6, a major downstream effector of mTORC1, together with several RAG GTPases like IQGAP1, required for efficient activation of mTORC1, which were largely restored after BX795 treatment (Table S3).

To confirm that the p.A53T mutation is causally related to dysregulation of protein metabolism and verify that BX795 can restore this effect in mature human neurons, we exploited the isogenic system of iCell DopaNeurons where we measured the levels of the activated form of RPS6, (phospho-RPS6; pRPS6), and the total protein synthesis rate. The presence of the p.A53T mutation led to a significant increase in the levels of pRPS6 (Fig. 7a,b) that correlated with a significant increase of global protein synthesis in iCell Dopa p.A53T neurons (Fig. 7c, d). BX795 could lower significantly the levels of pRPS6 and reverse the aberrantly increased protein synthesis rate (Fig. 7a, d). This data suggests that BX795 targets and restores dysregulated mRNA translation and protein synthesis pathways instigated by the p.A53T mutation in neuronal cells.

**Fig 7.**
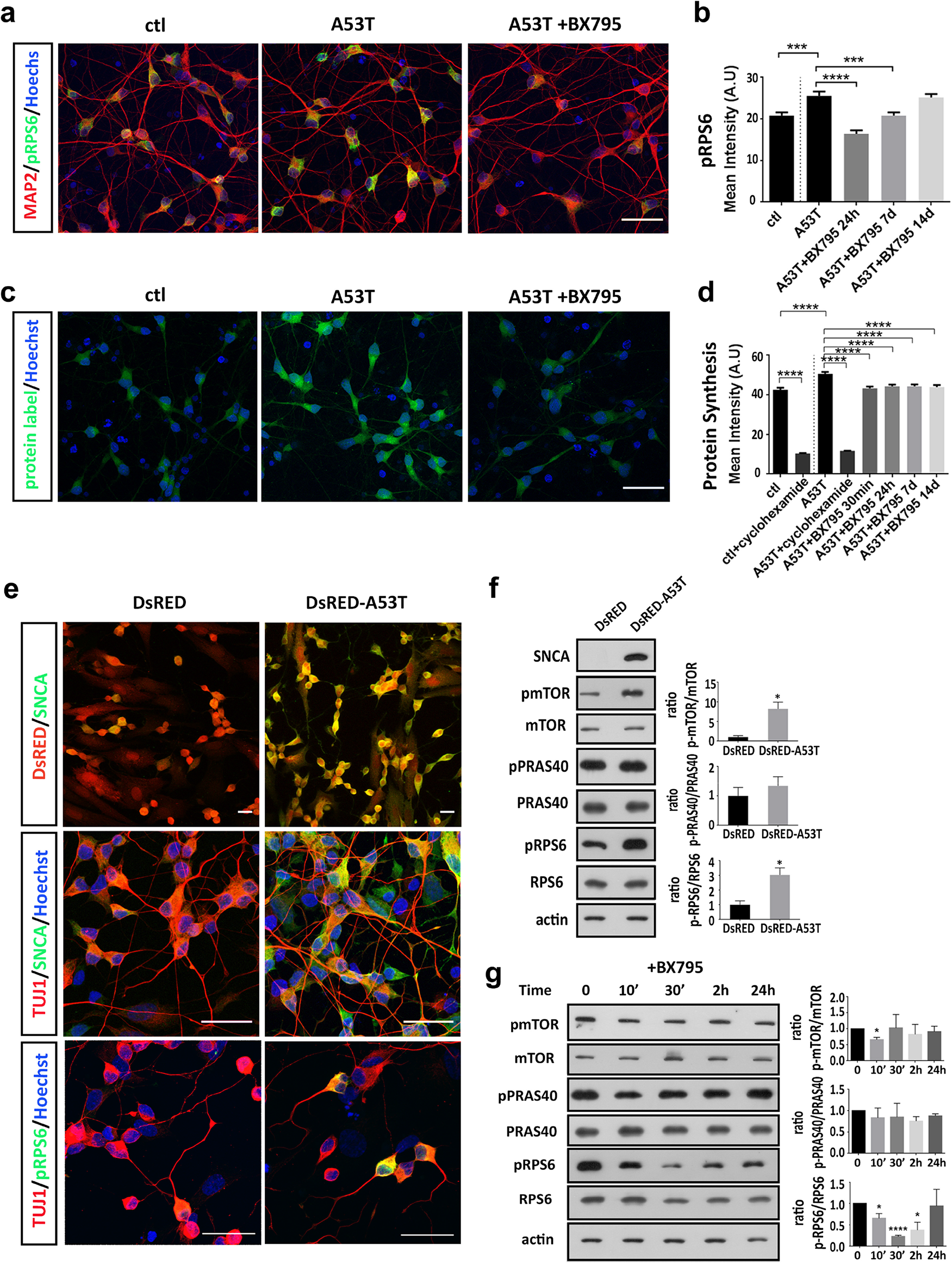
BX795 affects the mTORC1 signaling pathway to attenuate protein synthesis. a. Representative confocal images of control (ctl) and isogenic gene-edited p.A53T iCellDopa neurons, either non-treated or treated with BX795. Cells were immunolabeled for phosphorylated RPS6 (green) and microtubule associated protein 2 (MAP2; red). Nuclei are seen with Hoechst dye (blue). Scale bar, 30μm b. BX795 reduces phosphorylated RPS6 levels in p.A53T-neurons. Quantification of fluorescence intensity in control, untreated p.A53T or BX795-treated p.A53T neurons. Data represent mean ± SEM (Comparisons by ANOVA with Tukey correction, ***P< 0.001 ****P<0.0001, n = 100 randomly selected cells for each condition). c. Representative confocal images of control and isogenic gene-edited p.A53T iCellDopa neurons, non-treated or treated with BX795, labeled for total protein synthesis (protein label, green). Nuclei are visualized by Hoechst counterstaining (blue). Scale bar, 30μm d. BX795 reduces total protein synthesis in p.A53T-neurons. Quantification of fluorescence intensity in non-treated control and p.A53T neurons non-treated or treated with BX795. Cyclohexamide blocks protein synthesis in both genotypes and is used as a negative control. Data represent mean ± SEM (Comparisons by ANOVA with Tukey correction, ****P<0.0001, n = 100 randomly selected cells for each condition). e. Representative images of SH-SY5Y cells stably transduced to express DsRed only or DsRed and human pA53T-αSyn. After neuronal differentiation, cells were immunolabled for αSyn (SNCA), TUJ1 and pRPS6. f. Western blot showing that the presence of mutant SNCA in differentiated p.A53T-transduced SH-SY5Y cells, results in an increase in the levels p-mTOR and p-RPS6. Actin shows equal protein loading. Data represent mean ± SEM (*t*-test, *P<0.05, n=3 independent experiments). g. Western blot showing an acute reduction in the levels of p-mTOR and p-RPS6 in the above stably transduced and differentiated SH-SY5Y cells, in the presence of BX795. Actin shows equal protein loading. Data represent mean ± SEM (Comparisons by ANOVA with Tukey correction, *P<0.05, ****P<0.0001, n=3 independent experiments).

To examine further the effect of the p.A53T mutation on mTORC1 activity and protein synthesis, we created stably transduced SH-SY5Y neuroblastoma cells co-expressing the human p.A53T-αSyn and the fluorescent protein DsRed or DsRed only as a control (Fig7e). Upon neuronal differentiation, SH-SY5Y cells expressing the human p.A53T-αSyn displayed a prominent upregulation in the levels of phosphorylated mTOR and pRPS6 as compared to control cells (Fig 7e, f), whilst BX795 had an acute effect in downregulating their levels (Fig. 7g), as determined by Western blot analysis.

mTORC1 also controls autophagy, the major degradation pathway essential for removing aggregation-prone αSyn (56, 57). To test if BX795 could also affect this clearance pathway, we utilized a previously established inducible SH-SY5Y cell line that expresses the human p.A53T-αSyn upon withdrawal of Doxycycline (-Dox). In this model, expression of mutant p.A53T has been shown to cause perturbation of the autophagy lysosomal pathway resulting in increased steady-state levels of LC3II and p62 ((58); Fig. 8 a, b). p62 is a receptor for ubiquitinated cargo destined to be degraded by autophagy and is associated with LC3-II, the processed form of LC3, within autophagosomes and autolysosomes (59, 60). To visualize LC3-II and quantify GFP-LC3-II+ puncta comprising brightly fluorescent autophagosomes and more weakly labeled autolysosomes (Fig. 8a), we transfected the inducible SH-SY5Y line with a fusion construct containing the green fluorescent protein tagged to LC3 (GFP-LC3) (61). In agreement with the Western blot data, GFP-LC3-II+ puncta were scarce in p.A53T cells treated with DMSO while in the presence of BX795 there was a small, yet not significant increase (Fig. 8c, d). As expected, when DMSO-treated cells were exposed to bafilomycin, a blocker of autophagosome-lysosome fusion that prevents lysosome-mediated protein degradation, GFP-LC3-II+ puncta increased significantly (Fig. 8c, d)., Addition of both bafilomycin and BX795 further increased the number and brightness of GFP-LC3-II+ puncta, suggesting that BX795 acts as an autophagy inducer (Fig 8c, d).

**Fig 8.**
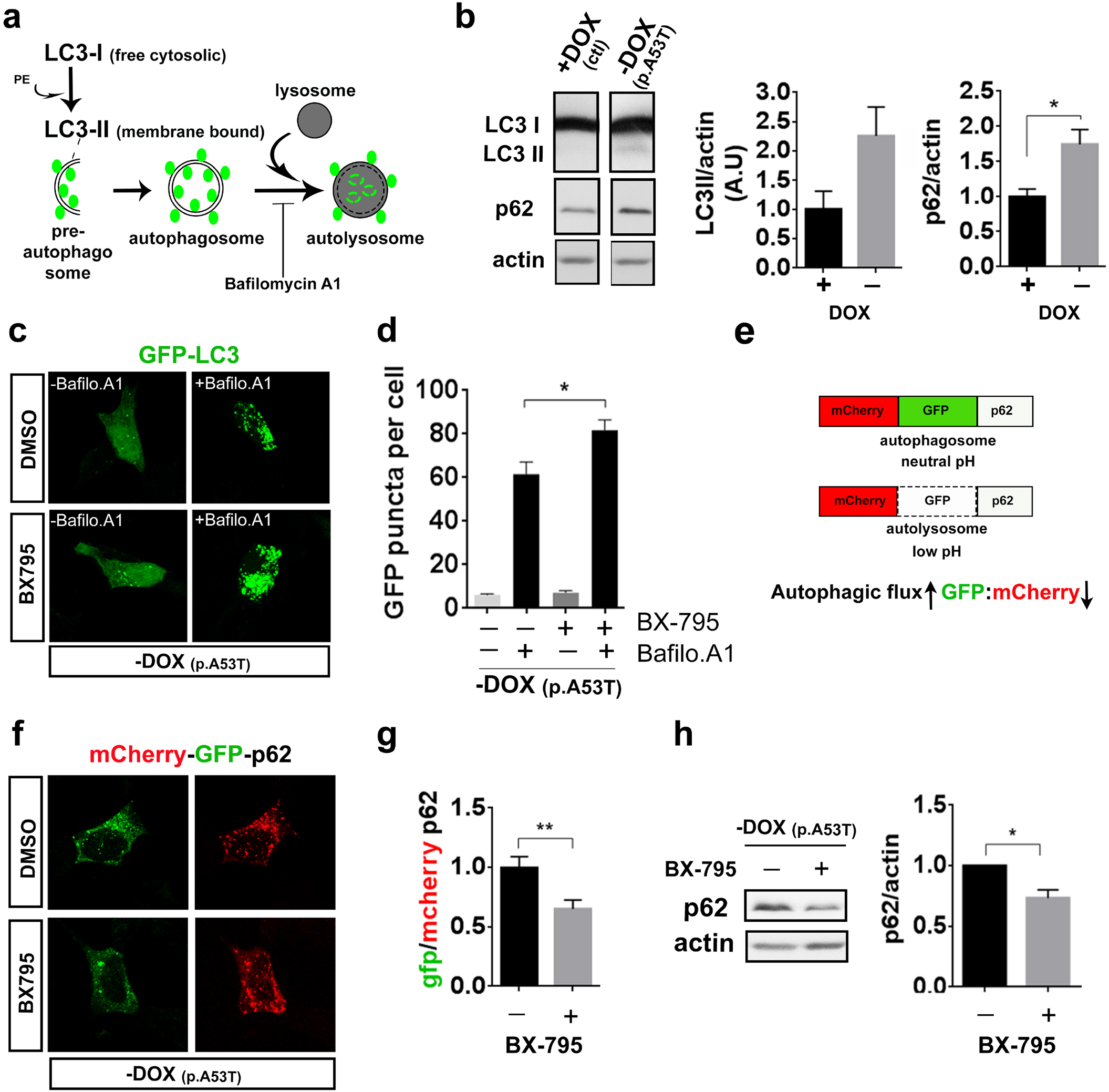
BX795 facilitates autophagy in an inducible SH-SY5Y cell line expressing human p.A53T-αSyn. a. Schema illustrating that cytosolic LC3 is cleaved to yield LC3-I, which is subsequently conjugated to phosphatidylethanolamine (PE) to form membrane-bound LC3-II (green circles). Pre-autophagosomal structures engulfing protein cargo and organelles destined for degradation close to form double membrane spherical autophagosomes. These fuse with lysosomes to yield autolysosomes and their contents are degraded. Bafilomycin blocks autophagic flux by inhibiting autophagosome-lysosome fusion, which results in accumulation of LC3-II+ autophagosomes. b. Representative immunoblot showing steady-state levels of LC3-II and p62 in lysates of inducible SH-SY5Y cells expressing the human p.A53T-αSyn (-Dox) and quantification relative to actin. Data represent mean ± SEM, *t*-test, *P<0.05, n=3 independent experiments. c. Representative confocal images of individual p.A53T SH-SY5Y cells (-Dox) transfected with GFP-LC3 that were treated or not with bafilomycin A1 in the absence or presence of BX795. d. Quantification of GFP-LC3 puncta per cell. Comparisons by ANOVA with Tukey correction.*P < 0.05, n=72 cells (control DMSO), n=79 cells (BX795), n=67 cells (Bafilomycin A1), n=68 cells BX795+Bafilomycin A1. Data are representative of three independent experiments). e. Assessment of autophagic flux using mCherry-GFP-LC3 color change between autophagosomes and autolysosomes. Autophagic flux is induced when the GFP:mCherry ratio is reduced. F, g. Representative confocal images of individual cells [inducible SH-SY5Y cell line expressing p.A53T-αSyn (-Dox)] transfected with GFP-mCherry-p62 that were treated with DMSO (control) or BX795 and quantification of the ratio of GFP+/mCherry+ puncta (*t*-test,n= 60 (control DMSO), n=53 (BX795) **P < 0.01 Data are representative of three independent experiments). h. Representative immunoblot showing steady-state levels of p62 in cells [inducible SH-SY5Y cell line expressing p.A53T-αSyn (-Dox)] treated or not with BX795, and quantification relative to actin. Data represent mean ± SEM, *t*-test, n=3 independent experiments.

To distinguish labeled autophagosomes from autolysosomes and monitor the autophagic flux, we used a dual fluorophore probe consisting of a tandem fluorescent mCherry-GFP-p62 construct (62). GFP fluorescence is sensitive to low-pH and labels only neutral-pH autophagosomes, while mCherry retains fluorescence in both autophagosomes and low-pH autolysosomes (60) (Fig. 8e). Calculation of the ratio of GFP+/mCherry+ puncta presents a measure of the autophagic flux, and a reduction in this ratio mirrors an increase in the progress of autophagy. Indeed, quantification of green and red puncta revealed a significantly lower GFP/mCherry ratio in the presence of BX795 as compared to DMSO-treated cells, indicating that BX795 facilitates the the autophagic flux (Fig. 8f, g). In agreement, a decrease in the total levels of p62 was noted upon treatment with BX95 (Fig. 8h).

Overall, our results indicate that BX795 can restore proteostasis in p.A53T cells by modulating aberrant protein synthesis and facilitating protein clearance mechanisms.

## Discussion

The generation of novel human models based on patient-derived iPSCs has opened up new perspectives for investigation of disease mechanisms and discovery of new therapeutics. In this work, we used a well-characterized human model of p.A53T pathology (22) to screen for small molecules with protective function. We identified the multi-kinase inhibitor BX795 as a compound that exerts a consistent and sustainable beneficial effect on patient-derived p.A53T-neurons. Remarkably, we found that a single treatment with BX795 has long-lasting consequences in supporting neuritic growth, limiting αSyn protein aggregate formation and restoring axonal neuropathology, recorded two weeks after its addition in human p.A53T neurons.

To our knowledge, this study represents the first high-content drug discovery screen performed in human p.A53T iPSC-derived neurons to identify candidate therapeutics for PD. Using an unbiased screening approach in combination with quantitative proteomics profiling, we were able to show that treatment with BX795 restored proteins associated with key cellular processes, most notably RNA metabolism, protein synthesis and degradation processes, as well as stress response, suggesting that restoration of proteostasis is key for rescuing the neuropathological features in p.A53T-neurons. Dissecting further the pathways affected by BX795, we demonstrated that BX795 modulates the mTORC1 pathway to restrict excessive protein synthesis and facilitate autophagy. Taken together, our data highlight the BX795 kinase inhibitor as a promising compound and candidate therapeutic that ameliorates p.A53T-associated pathology.

Considerable progress in understanding the neurotoxic properties of α-Syn has been achieved by exploiting causal mutations resulting in rare familial forms of PD, most notably the p.A53T-αSyn mutation (G209A in the *SNCA* gene) (63, 64). We and others have shown that disease-associated characteristics can be recapitulated in patient-derived p.A53T-neurons, including axonal degeneration and accumulation of protein inclusions resembling Lewy bodies and neurites (22). These have been linked to multiple molecular defects in mRNA processing and translation, endocytic and retrograde trafficking (38, 42), protein misfolding, redox homeostasis (20, 33) and the synaptic protein machinery (22). The p.A53T proteome examined here revealed a profound increase in proteins related to the biological processes of RNA metabolism, protein synthesis, modification and transport, protein clearance and stress response. Notably, the cohort of 118 proteins that was specifically restored in p.A53T-neurons upon treatment with BX795, was associated with these key cellular processes.

The pathways affected by mutant αSyn in our study have a high similarity with the αSyn connectome reported by Chung et al (38) for mouse neurons, and the predictions of the *in silico* “humanized” map of αSyn proteotoxicity reported in the accompanying study of Khurana et al (42). Our proteomics analysis, the first accomplished in p.A53T-human neurons, identified perturbations in RNA metabolic processes that started from the nucleus and reached the ribosome. Alternative mRNA processing greatly increases the dimensions of gene expression through splicing, polyadenylation, targeted localization and post-transcriptional silencing. Neurons take advantage of all these strategies as the brain has the highest levels of alternative splicing compared to any other human tissue (65). This process has recently been shown to be defective in the PS19 Tau model of Alzheimer’s disease, where alternative splicing events affected genes particularly involved in synaptic transmission (66). Similarly, the p.A53T-proteome suggests that this process could be excessively induced in p.A53T-neurons as a number of RBPs known to be linked to αSyn aggregation have emerged, including ELAV1, ELAV3 and CELF, suggesting a possible association with the abnormal expression of synaptic genes and the defective synaptic connectivity we have previously reported in p.A53T neurons-(22).

An excess of mRNAs coming out of the nucleus in p.A53T-neurons could explain the abnormal expression of proteins involved in translation, the next step of mRNA processing. The significant increase of components of the tRNA splicing ligase complex, various aminoacyl-tRNA synthetases, ribosomal subunits and eukaryotic translation initiation factors indicate an enhanced translation of spliced mRNAs. Moreover, in post-mortem PD brains, region and stage-dependent alterations in the machinery of protein synthesis have been reported and have been associated with α-synuclein oligomers in remaining neurons (67).

The mTOR kinase is a master regulator of cellular metabolism that functions in two distinct complexes: mTORC1 and mTORC2 (68) with the first implicated in protein and lipid biosynthesis through a signaling cascade that includes SK6 and 4E-BP1 proteins (69). Unlike proliferating cells where this pathway is utilized for growth and division, in neurons it acts as a regulator of healthy metabolism and aging (70) with its restriction being associated with prolonged life span and delay of age-related pathologies. p.A53T neurons have increased RPS6, IQGAP1 and RAG-GTPases, components of mTORC1 pathway and this seems to be associated with an increased translation of a subset of mRNAs that are linked to RNA metabolism and the stress response. Similarly, a quantitative proteomics study of a pre-symptomatic p.A53T-αSyn Drosophila model shows significant upregulation of ribosomal proteins in the p.A53T flies (71). Although the mechanistic link between p.A53T-αSyn and mTORC1 remains to be established, recent evidence shows that genetic variability in the mTOR pathway contributes to SNCA effects in disease pathogenesis (72).

Concomitantly with promoting protein synthesis mTORC1 acts to repress autophagy through ULK1 phosphorylation. Autophagy has a central role in promoting health and longevity while this process is impaired in neurodegenerative diseases and αSyn pathology (73, 74). The p.A53T-proteome shows that neurons are under stress as proteins involved in the UPR or the heat-shock stress response, proteasome assembly and regulation, known to be orchestrated by mTORC1 in neurons, are significantly upregulated (70). Restoration of numerous components of RNA metabolism and protein translation cascades by BX795 is directly related to the diminished stress response that emerges by the lower levels of UPR and heat-shock-associated proteins also conferred by this molecule. In parallel, a significant number of ubiquitin/proteasome-associated proteins were brought back to near control levels suggesting that BX795 helps misfolded protein clearance by limiting protein synthesis. This is in agreement with its demonstrated ability to decrease protein aggregates in p.A53T-neurons, as shown in this study, along with facilitation of autophagy in SYSH-5Y cells expressing p.A53T.

BX795 is a multi-kinase inhibitor that targets numerous pathways, including the kinases TBK1 and PDK1 (29–31, 35). Although in our system differences in the total or phosphorylated levels of these two kinases were not observed in the presence of BX795 (data not shown), we cannot exclude that its effects are mediated through these two kinases as both are involved in neurodegeneration, mTOR signaling and autophagy (75, 76). Yet four other PDK1 inhibitors that were included in the Selleck library did not emerge as hits during the screening campaign. Interestingly, even though we demonstrated an acute effect of BX795 in mTOR and RPS6 in p.A53T-expressing cells, multiple other inhibitors of mTOR phosphorylation present in the kinase inhibitor library tested (26 in total, including rapamycin), failed to show any protective effects. Considering that BX795 has been proposed to act through distinct mechanisms in different pathologies, future mechanistic studies should reveal its direct targets in p.A53T neurons. Nevertheless, the work presented here uniquely identifies BX795 as a promising compound that may have therapeutic potential for patients with PD and other protein conformational disorders. Further, our collective data along with previous proteomics and systems approaches shed light into the molecular and cellular pathways of αSyn proteotoxicity unveiling new disease targets for the development of combined therapeutics.

## Materials and Methods

### iPSC lines

iPSCs used in this study were previously generated and characterized from two Parkinson’s disease patients harboring the p.A53T-α-synuclein mutation and a healthy subject (control, wild-type SNCA) (22). All procedures for generation of human iPSCs were approved by the Scientific Council and Ethics Committee of Attikon University Hospital (Athens, Greece), which is one of the Mendelian forms of Parkinson’s disease clinical centers, and by the Hellenic Pasteur Institute Ethics Committee overlooking stem cell research. Written informed consent was obtained from all donors before skin biopsy.

### Directed neuronal differentiation

For directed differentiation, iPSCs were allowed to form embryoid bodies and neural induction was initiated by applying a dual SMAD inhibition protocol in the presence of Noggin and TGFβ inhibitor for generation of neural precursor cells (NPCs) (22). NPCs were expanded in DMEM/F12/B27/N2-medium supplemented with HEPES, Glutamax, non-essential amino acids [NEAA] and 20ug/ml FGF2. For neuronal differentiation, NPCs were dissociated with accutase and seeded onto poly-L-ornithine (20 μg/ml; Sigma-Aldrich)/laminin (5 μg/ml; Sigma-Aldrich)-coated dishes in DMEM/F12/ B27/N2-medium supplemented with 200 ng/ml human recombinant sonic hedgehog (SHH, R&D Systems) and 100 ng/ml murine recombinant fibroblast growth factor 8b (FGF-8b, R&D Systems) for 7 days in vitro (DIV). Cells were then replated in medium supplemented with 20 ng/ml brain-derived neurotrophic factor (BDNF, R&D Systems), 20 ng/ml glial cell-derived neurotrophic factor (GDNF, R&D Systems), 200 μM ascorbic acid (AA, Sigma-Aldrich) and 0.5 mM cyclic AMP (cAMP, Sigma-Aldrich). The medium was changed every 2 to 3 days for 2 weeks.

### iCell Dopa neurons and isogenic iCell DopaNeurons PD SNCA A53T HZ

Commercially available iCell DopaNeurons 01279, Catalog No C1028, and a heterozygous (HZ) A53T allelic variant isogenic to iCell DopaNeurons, PD SNCA A53T HZ 01279, Catalog No C1113, in which the site-specific p.A53T mutation was introduced into the *SNCA* gene by nuclease-mediated SNP alteration, were purchased from Fujifilm Cellular Dynamics International and were maintained according to the User’s Guide protocol for two weeks.

### Compound screening and High Content image analysis

iPSC-derived NPCs at 7 DIV were dissociated with accutase, seeded (9,000 cells/well) onto poly-L-ornithine/ laminin-coated 384-well optical bottom plates containing the kinase inhibitors (Greiner Bio-One, Kremsmünster, Austria) and cultured in neuronal differentiation medium for two weeks (Fig. 1a). A collection of 273 small molecule kinase inhibitors from Selleck Chemicals was used. The list of inhibitors and their known targets according to the provider, is shown in Table S1. The compounds were dispensed in duplicate in 384-well optical bottom plates at a final concentration of 1μM, followed by NPC seeding. After 2 weeks of neuronal differentiation, cells were fixed in 4% paraformaldehyde for 20 min followed by immunofluorescence for βΙΙΙ-tubulin (TUJ1) and Tyrosine hydroxylase (TH) at 4°C overnight and incubation with appropriate secondary antibodies (Molecular Probes, Thermo Fisher Scientific) conjugated to AlexaFluor 488 (green) or 546 (red), for at least 1 h at room temperature. Nuclei were stained with Hoechst dye. Images were captured by automated confocal microscopy (Opera High-Content Screening System, Perkin Elmer, Hamburg, Germany). A total of 15 images per well were acquired using a 10X magnifying objective. Cell nuclei and fluorescence staining were quantified by segmentation on 15 images per well in a duplicate experimental setup. Parameters were set as follows: primary object detection (cell nuclei) was based on Hoechst staining, captured in channel 1. Detection of neurons was based on TUJ1 immunofluorescence signal, captured in channel 2 and on TH immunofluorescence signal, captured in channel 3. For quantification of TUJ1 and TH intensity Image Mining was used, a custom-made image processing and analysis application with an extendable “plug-in” infrastructure (77).

### RNA isolation, cDNA Synthesis and qRT-PCR

Total RNA was extracted from cell pellets using the TRIzol® Reagent (Life Technologies). Following digestion with DNaseI, 1 μg of total RNA was used for first strand cDNA synthesis with the ImProm-II Reverse Transcription System (Promega) according to the manufacturer’s instructions. Quantitative RT-PCR analyses were carried out in a Light Cycler 96 (Roche) real time PCR detection system using KAPA SYBR FAST qPCR Master Mix (KapaBiosystems). All primers used are listed in Table S2.

### Immunofluorescence staining

Cells were paraformaldehyde-fixed, blocked with 5% donkey serum in PBS/ 0.1% Triton X-100 (Sigma-Aldrich) for 30 min and immunofluorescence labelled as above. Coverslips were mounted with ProLong Gold antifade reagent with DAPI (Cell Signaling) and images were acquired using a Leica TCS SP8 confocal microscope (LEICA Microsystems) and analyzed using ImageJ software (NIH).

### Neurite analysis

Neurite length was estimated manually by tracing the length of all neurites on TH-labeled neurons at 21 DIV using the NeuronJ plugin of ImageJ (NIH). At least 50 single TH+ neurons per sample were analyzed.

### Axon degeneration index

The number of TUJ1+ spots in blebbed or fragmented axons was counted manually (ImageJ) on twenty randomly selected fields and the ratio between the number of spots and the total TUJ1+ staining area (ImageJ) was defined as axon degeneration index [22].

### Protein aggregate quantification

Protein aggregates were detected with the PROTEOSTAT Aggresome Detection Kit (Enzo) followed by immunolabeling for TUJ1 or TH (22, 32). Manual analysis was performed by isolating individual cells from images (ROIs), applying a threshold, and utilizing the ‘analyze particles’ ImageJ function.

### Proteomic Analysis

iPSC-derived neurons at 21 DIV were suspended, lysed and the proteins reduced in 4% SDS, 100 mM DTT, 100 mM Tris pH 7.8 through heating for 5 min. Next, the proteins were alkylated by 100 mM iodoacetamide treatment for 30 min in the dark. Samples were further processed according to the Single-Pot Solid-Phase enhanced Sample Preparation (SP3) method of Hughes et al (78). Digestion was carried out overnight at 37°C using Trypsin/LysC mix (Promega) at a protein/enzyme ratio of 50:1 in a ThermoMixer under continuous mixing at 1000 rpm. After digestion, the tubes were placed on a magnetic rack, and the supernatant containing the peptides was collected and dried down in a centrifugal evaporator (Savant SPD 1010, Thermo scientific). The peptide mixtures were reconstituted in a solution of 2% (v/v) ACN/ 0.1% (v/v) formic acid and incubated for 3 min in a sonication water bath. Peptide concentration was determined by nanodrop absorbance measurement at 280 nm.

### Ultra-high pressure nanoLC

2.5 μg peptides were pre-concentrated with a flow of 3 μL/min for 10 min using a C18 trap column (Acclaim PepMap100, 100 μm x 2 cm, Thermo Scientific) and then loaded onto a 50 cm long C18 column (75 μm ID, particle size 2 μm, 100Å, Acclaim PepMap100 RSLC, Thermo Scientific). The binary pumps of the HPLC (RSLCnano, Thermo Scientific) consisted of Solution A (2% (v/v) ACN in 0.1% (v/v) formic acid) and Solution B (80% (v/v) ACN in 0.1% (v/v) formic acid). The peptides were separated using a linear gradient of 4% B up to 40% B in 340 min with a flow rate of 300 nL/min. The column was placed in an oven at 35°C.

### LC-MS/MS

Eluted peptides were ionized by a nanospray source and detected by an LTQ Orbitrap XL mass spectrometer (Thermo Fisher Scientific, Waltham, MA, USA) operating in a data dependent mode (DDA). Full scan MS spectra were acquired in the orbitrap (m/z 300– 1600) in profile mode with resolution set to 60,000 at m/z 400 and automatic gain control target at 106 ions. The six most intense ions were sequentially isolated for collision-induced (CID) MS/MS fragmentation and detection in the linear ion trap. Dynamic exclusion was set to 1 min and activated for 90 sec. Ions with single charge states were excluded. Lockmass of m/z 445,120025 was used for continuous internal calibration. XCalibur (Thermo Scientific) was used to control the system and acquire the raw files.

### Protein identification and quantification

The raw mass spectral files were processed using MaxQuant software (version 1.6.9.0) with default parameters for protein identification and quantification. Trypsin specificity was set to allow two missed cleavages and minimum peptide length was set to 7 amino acids. Cysteine carbamidomethylation was set as fixed, and methionine oxidation, deamidation of asparagine and glutamine and N-terminal acetylation were set as variable modifications. A maximum of 5 modifications per peptide was set. The false discovery rate both for peptide and protein was set to 1%. For calculation of protein abundances, label-free quantification (LFQ) was performed with both “second peptides” and “match between run” options enabled. The human FASTA files were from UniProt downloaded on 15 October 2019.

### Proteomic data analysis

Statistical analysis was performed using Perseus (1.6.6.0). Proteins identified as contaminants, “reverse” and “only identified by site” were filtered out. The LFQ intensities were transformed to logarithmic values [log2(x)]. The protein groups were filtered to obtain at least 2 valid values in at least one group. The label-free quantified proteins were subjected to statistical analysis with ANOVA test (permutation-based p-value with 0.05 cutoff). LC-MS/MS data after statistical analysis were plotted in a volcano graph based on the difference between the two samples expressed as log2(x) versus their statistical significance expressed as –Log10(p-value). Hierarchical clustering was carried out on Z-score transformed LFQ values using average linkage of Euclidian distance. For statistical and bioinformatics analysis, as well as for visualization, Perseus, which is part of Maxquant, was used (79). GO Enrichment analysis for biological processes, molecular function and cellular compartment was performed using DAVID functional annotation tools with official gene symbol as identifiers, the Homo sapiens background and the GOTERM_DIRECT annotation categories. A P value of 0.05 was selected as the cutoff criterion. The enrichment of proteins involved in signaling pathways was performed using the Reactome pathway database. A P value of 0.01 was selected as the cutoff criterion.

### Western blot

Cells were lysed at 4°C for 15 min in ice cold lysis buffer [150mMNaCl, 50 mM Tris pH 7.5, 1% Triton X-100, 1mM EDTA, 1mM EGTA, 0.1% SDS, 0.5% sodium deoxycholate containing PhosSTOP phosphatase inhibitors and a complete protease inhibitor mixture (Roche Life Science)], and centrifuged at 20,000 g. Protein concentration was estimated in the supernatant by Bradford assay (Applichem). Proteins were separated by SDS-polyacrylamide gel electrophoresis and transferred onto nitrocellulose membranes (Maine Manufacturing). For phospho-(Ser129)-αSyn detection, the membrane was heated at 65 ^0^C overnight in PBS. Nonspecific binding sites were blocked in TBS/ 0.1% Tween 20/ 5% skimmed milk for 1 hour at 20 ^0^C followed by overnight incubation with primary antibodies diluted in the same buffer. Incubation with appropriate HRP-conjugated secondary antibodies (Thermo) was for 2 hours at room temperature and protein bands were visualized using the Clarity Western ECL Substrate (BIO-RAD). Densitometric analysis was performed using ImageJ software (NIH).

### Production of CMV.DsRed and CMV.DsRed.A53T lentiviral vectors

Four plasmids were used for lentivirus generation: the lentiviral transfer vector and three lentiviral packaging vectors (pMDL, pRev and pVSVG; provided by Dr. Fred Gage, the Salk Institute for Biological Studies). The lentiviral transfer vectors for expression of either the red fluorescent protein DsRed under the control of CMV promoter (LV.CMV.DsRed) or for co-expression of the red fluorescent protein DsRed, a T2A bicistronic configuration and human p.A53T-αSyn under the control of CMV promoter (LV.CMV.DsRed.T2A.A53T) were constructed by VectorBuilder. The preparation and purification of the lentiviral vectors were performed as previously described (80).

### Generation of stably transduced SH-SY5Y cells

SH-SY5Y cells were transduced with the control vector LV.CMV.DsRed or LV.CMV.DsRed.T2A.A53T for expression of DsRed or co-expression of DsRed and human p.A53T-αSyn. Transduced cells were maintained in regular RPMI 1640 medium/ 10% FBS (Gibco)/ 1% penicillin/streptomycin (Life Technologies) for 48h with one change of medium, and were then transferred in selection medium containing 300 μg/ml gentamycin-disulfate G418. After 3 weeks of selection, when 100% of cells expressed the DsRED protein, they were frozen as a polyclonal pool.

### Differentiation of SH-SY5Y cells

Cells were plated on PLL/Laminin coated plates (2×10^4^ cells/cm^2^) in regular RPMI 1640 medium/ 5% FBS/ 1% penicillin/streptomycin (DIV 0). The following day, 10 μM Retinoic Acid (RA) was added (DIV1). On DIV3, the medium was changed to Neurobasal supplemented with B27, N2, Glutamax and BDNF (50ng/ml) with fresh medium added every 2-3 days until DIV9.

### Cell culture and transfection of an inducible SH-SY5Y line expressing human p.A53T-αSyn

The inducible SH-SY5Y cell line, in which expression of p.A53T-αSyn was switched off in the presence of doxycycline (Dox, 2 µg/mL), was previously reported (58). Transfection with GFP-LC3 or mCherry-GFP-p62 plasmids (provided by Dr Tamotsu Yoshimori, Osaka University, Japan and Dr Terje Johansen, University of Tromso, Norway, respectively) was performed in the absence of Dox using Lipofectamine 2000, according to the manufacturer’s protocol (Invitrogen; Thermo Fisher Scientific, Inc.).

### Protein synthesis assay

For detection of total protein synthesis, an assay Kit (ab239725; Abcam) was used that utilizes a cell permeable analog of puromycin, which once inside the cell, stops translation by forming covalent conjugates with nascent polypeptide chains. Truncated polypeptides can be detected based on a click reaction with fluorescent azide. Cells were pre-treated with DMSO vehicle or BX795 for different time points and were incubated for 2h with fresh aliquots of media containing either Protein Label or Protein Label and BX795. Cyclohexamide that blocks protein synthesis was used as a negative control. Fluorescence images were acquired using a Leica TCS SP8 confocal microscope (LEICA Microsystems) and analyzed using ImageJ software (NIH).

### Statistics

All experiments were replicated at least three times and data from parallel cultures were acquired. Statistical analysis was performed using GraphPad Prism 6 software. Before performing parametric tests, data were assessed for normality with a D’Agostino– Pearson omnibus. Statistical significance was calculated for two groups using Student’s t-tests or the Mann-Whitney test for non-parametric distribution. Group comparisons of data were performed by one-way ANOVA test followed by Tukey post hoc test using PRISM (Graph Pad). P-values < 0.05 were considered significant; *p < 0.05, **p < 0.01, ***p < 0.001, ****p<0.0001.

### Study approval

All studies on human pluripotent stem cells were approved by the Hellenic Pasteur Institute Ethics Committee overlooking stem cell research.

## Data availability

The mass spectrometry proteomics data have been deposited to the ProteomeXchange Consortium via the PRIDE partner repository with the dataset identifier PXD019574.

## Acknowledgements

We thank Drs. Tamotsu Yoshimori and Terje Johansen for providing GFP-LC3 and mCherry-GFP-p62 plasmids, respectively. This work was supported by: a Stavros Niarchos Foundation grant to the Hellenic Pasteur Institute as part of the Foundation’s initiative to support the Greek Research Center ecosystem; the Greek General Secretariat for Research and Technology (GSRT) grant BIOIMAGING-GR MIS 5002755 implemented under the Action “Reinforcement of Research and Innovation Infrastructure”, funded by the Operational Programme “Competitiveness, Entrepreneurship and Innovation” (NSRF 2014-2020) and co-financed by Greece and the European Union (European Regional Development Fund); the GSRT Flagship Action for Neurodegenerative Diseases on the basis of Personalized Medicine; the Hellenic Foundation for Research and Innovation 899-PARKINSynapse grant to G.K; the South Korea Ministry of Science and ICT (MSIT) grant (NRF-2017M3A9G6068257) to RG. N.A. was recipient of a Calmette & Yersin Fellowship for a technology exchange visit to the Institut Pasteur Korea.

## Author contributions

NA carried out the experiments, analyzed and interpreted the data, generated the figures, participated in the study design and in writing the manuscript. KP and ET analyzed proteomics data. GK generated the patient-derived p.A53T and control iPSCs used in this study and provided training on iPSC culture and differentiation. MS and GP performed the proteomic analysis. MX and LS provided reagents, analytic tools and guidance for autophagy experiments. NA and RG performed high-content imaging and drug screening on p.A53T neurons. ET and RM conceived, designed and supervised the study, analyzed the data and wrote the paper with contribution from all authors.

---

**Fig S1.**
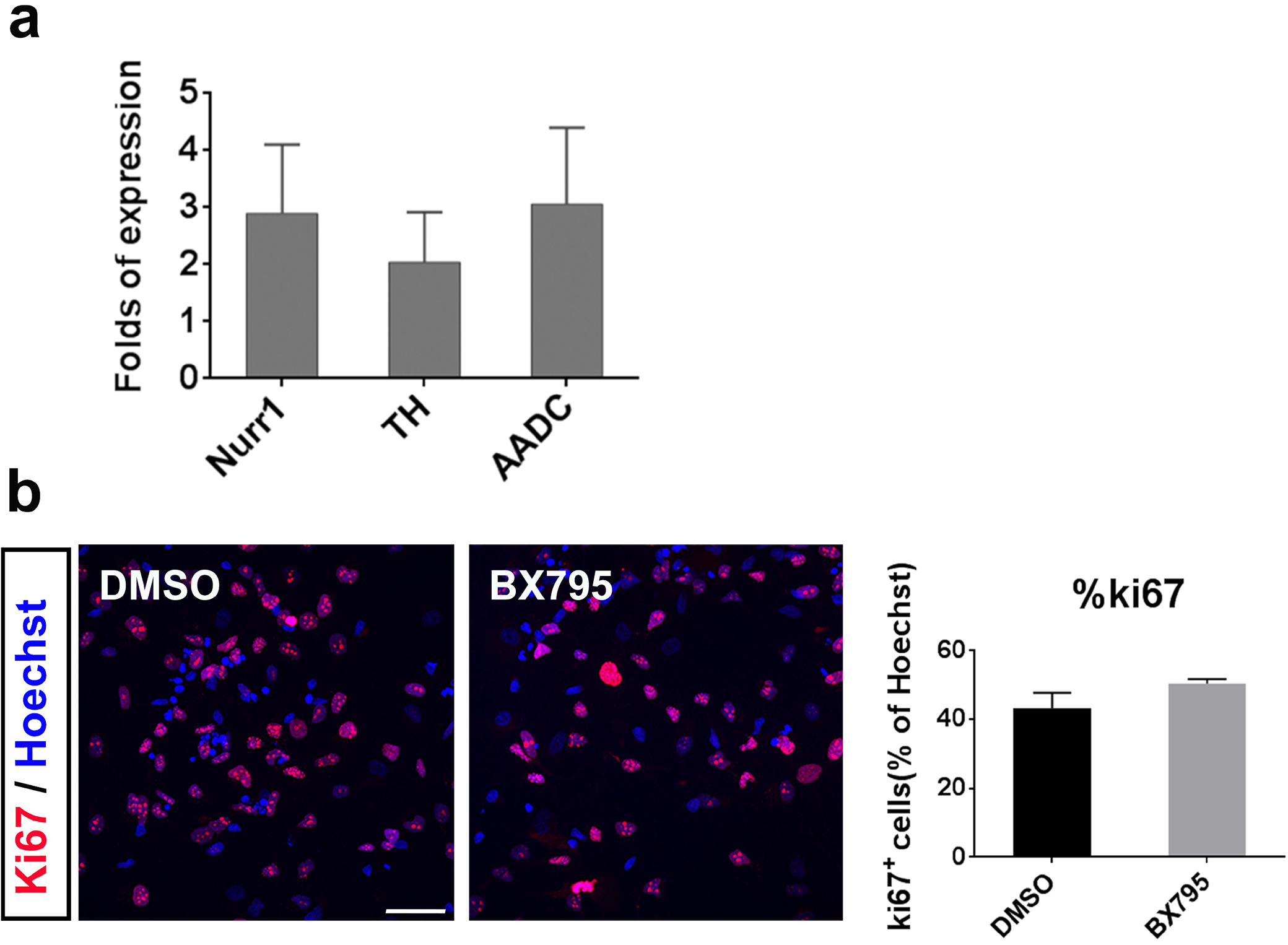
Expression of dopaminergic markers in patient p.A53T-iPSC-derived neurons. a. RT-qPCR analysis of selected domaminergic markers in p.A53T iPSC-derived neurons at 21 DIV: Tyrosine Hydrosylase (TH), Nuclear receptor related 1 protein (Nurr1) and Aromatic L-amino acid decarboxylase (AADC) normalized to GAPDH levels. Data represent mean ± SEM (n = 3). Student’s t-test was used. b. Representative images of p.A53T iPSC-derived neurons at 21 DIV immunostained for Ki67 (red) to label cycling cells. Hoechst+ nuclei are in blue (Scale bar, 50 μm). Quantification of the percentage of Ki67+ cells in the presence or absence of BX795. Data represent mean ± SEM (n = 3). Student’s t-test was used.

**Fig S2.**
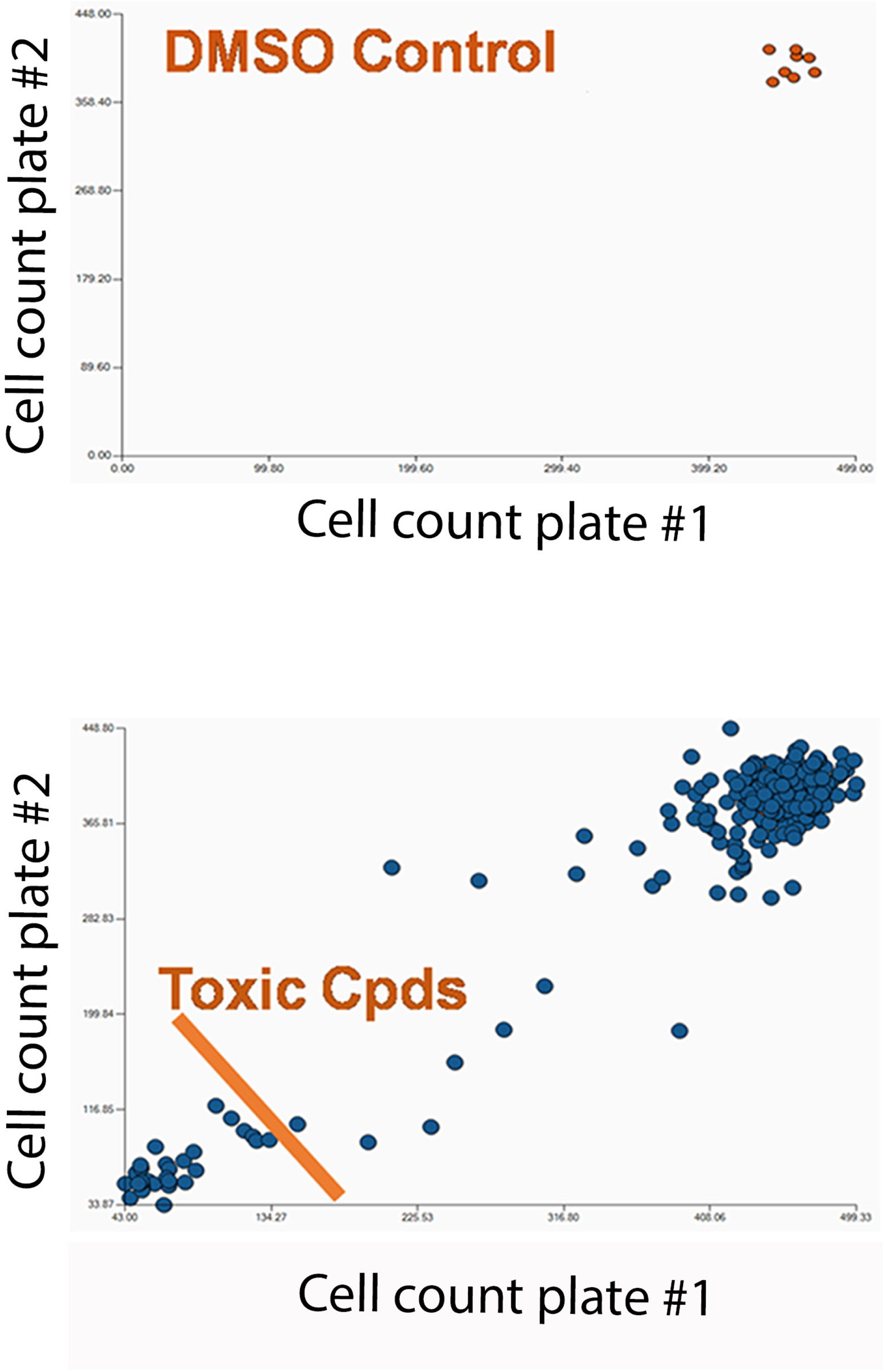
Identification of toxic compounds in the small molecule library of kinase inhibitors. Summary of total nuclei counts from two screening plates. Compounds in cells with low nuclei counts were considered toxic and where excluded from the analysis. Each assay plate was normalized to DMSO.

**Fig S3.**
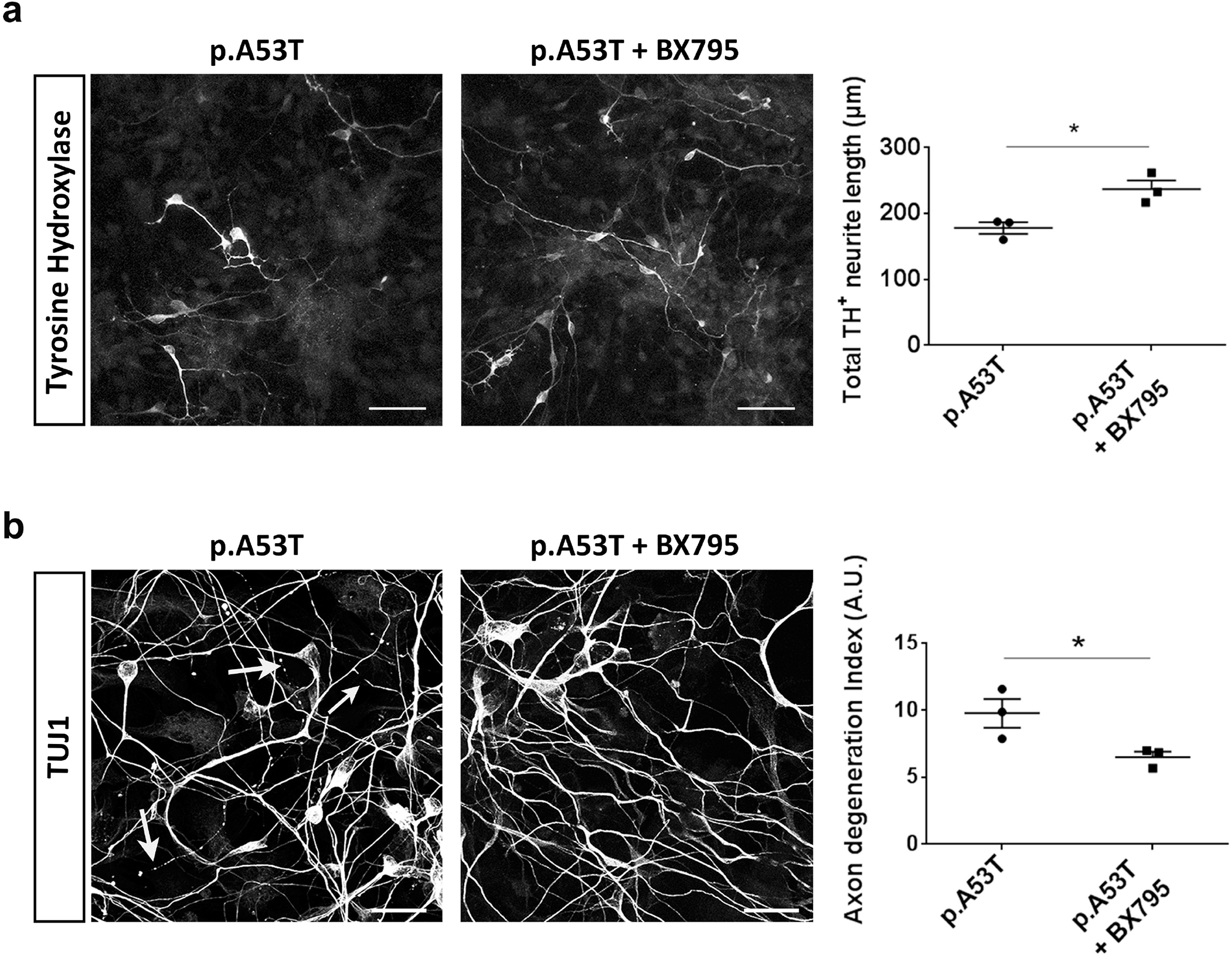
Rescue of neuropathological features by BX795 in p.A53T neurons from a second patient. a. BX795 has a positive effect on neurite length of p.A53T-neurons. Representative confocal images of p.A53T-neurons immunostained for TH and quantification of total neurite length of TH+ cells. Data represent mean ± SEM. Student’s t-test was used. Scale bar, 50μm. b. BX795 alleviates axonal neuropathology in p.A53T-neurons as demonstrated by immunostaining for βIII-tubulin (TUJ1; confocal images). Neurites with swollen varicosities or fragmented processes are indicated with arrows. Scale bar, 30μm. Axonal degeneration is estimated in the accompanying graph by measuring the ratio of TUJ1+ spots over the total TUJ1+ area in untreated (DMSO) or BX795-treated p.A53T-neurons. Data represent mean ± SEM. Student’s t-test was used.

**Fig S4.**
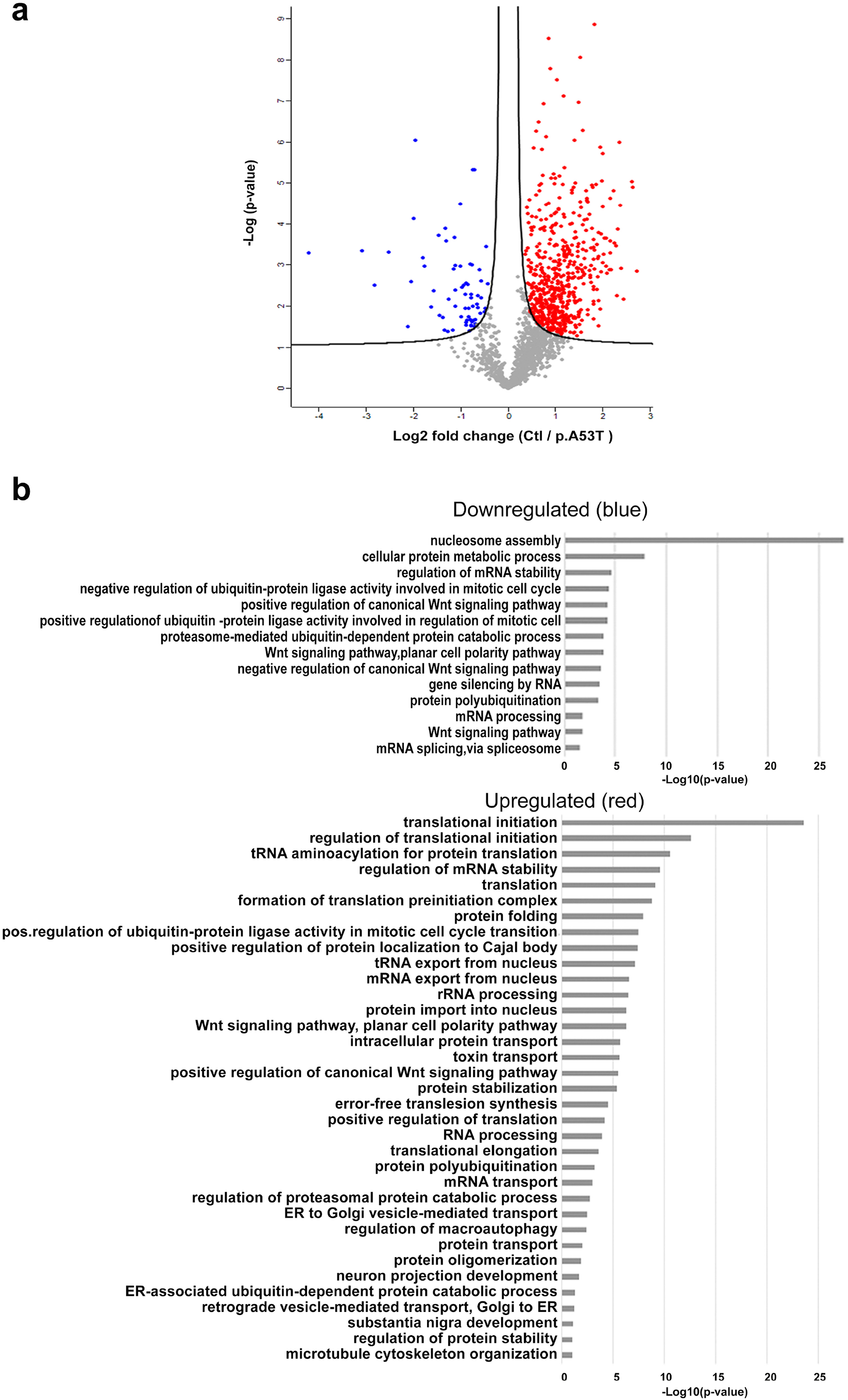
Identification of the biological processes that are dysregulated in p.A53T neurons. a. Volcano plot of differentially expressed proteins between control and patient-derived p.A53T-neurons assessed by quantitative proteomics analysis. Each point represents the difference in expression (fold-change) between the two groups plotted against the level of statistical significance. Blue dots correspond to proteins downregulated in p.A53T neurons while red dots show proteins upregulated in p.A53T neurons (FDR=0.05, S0 = 0.1, as indicated by black lines). b. GO enrichment analysis for biological processes of the differentially expressed proteins was performed using DAVID software (p<0.05).

**Table S1:**
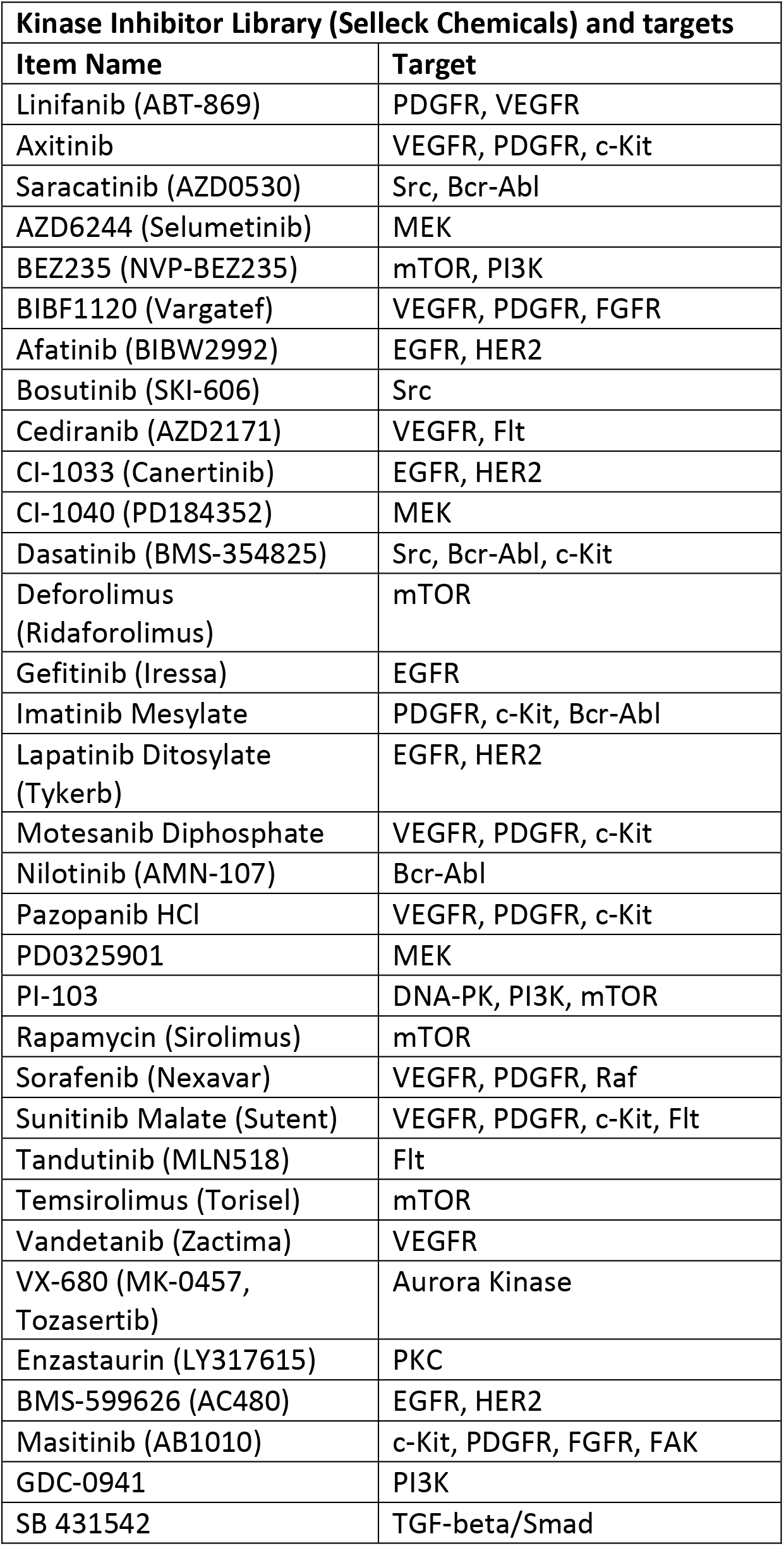
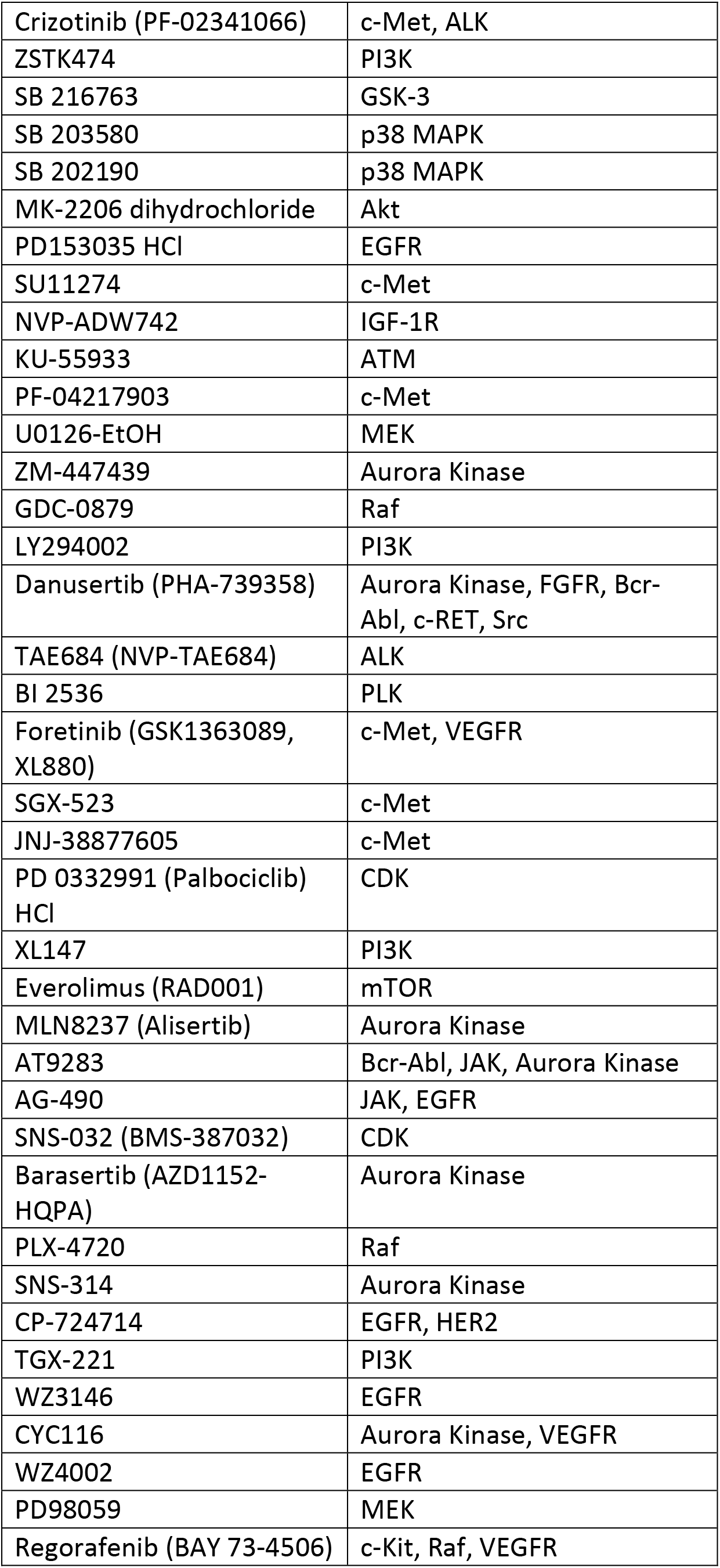
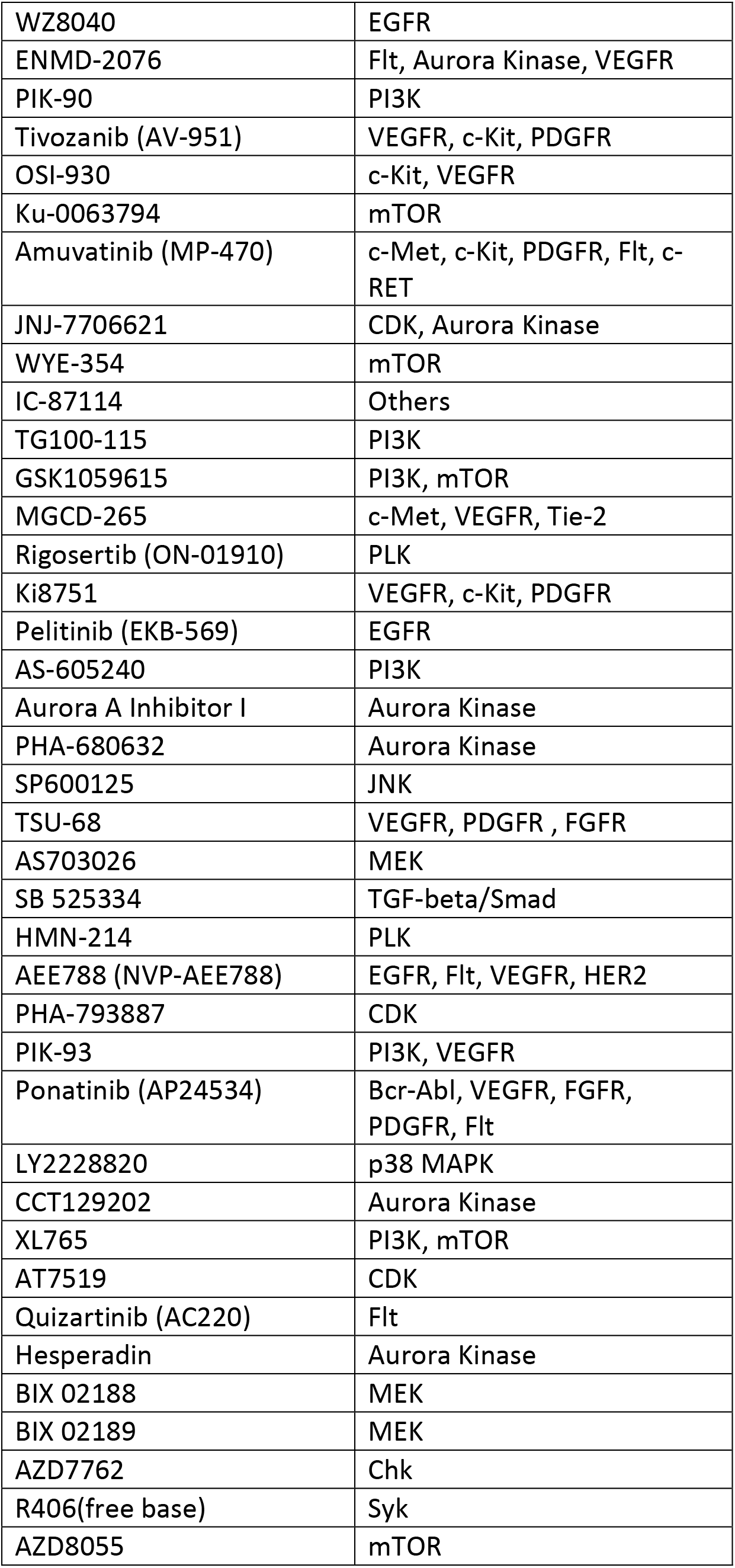
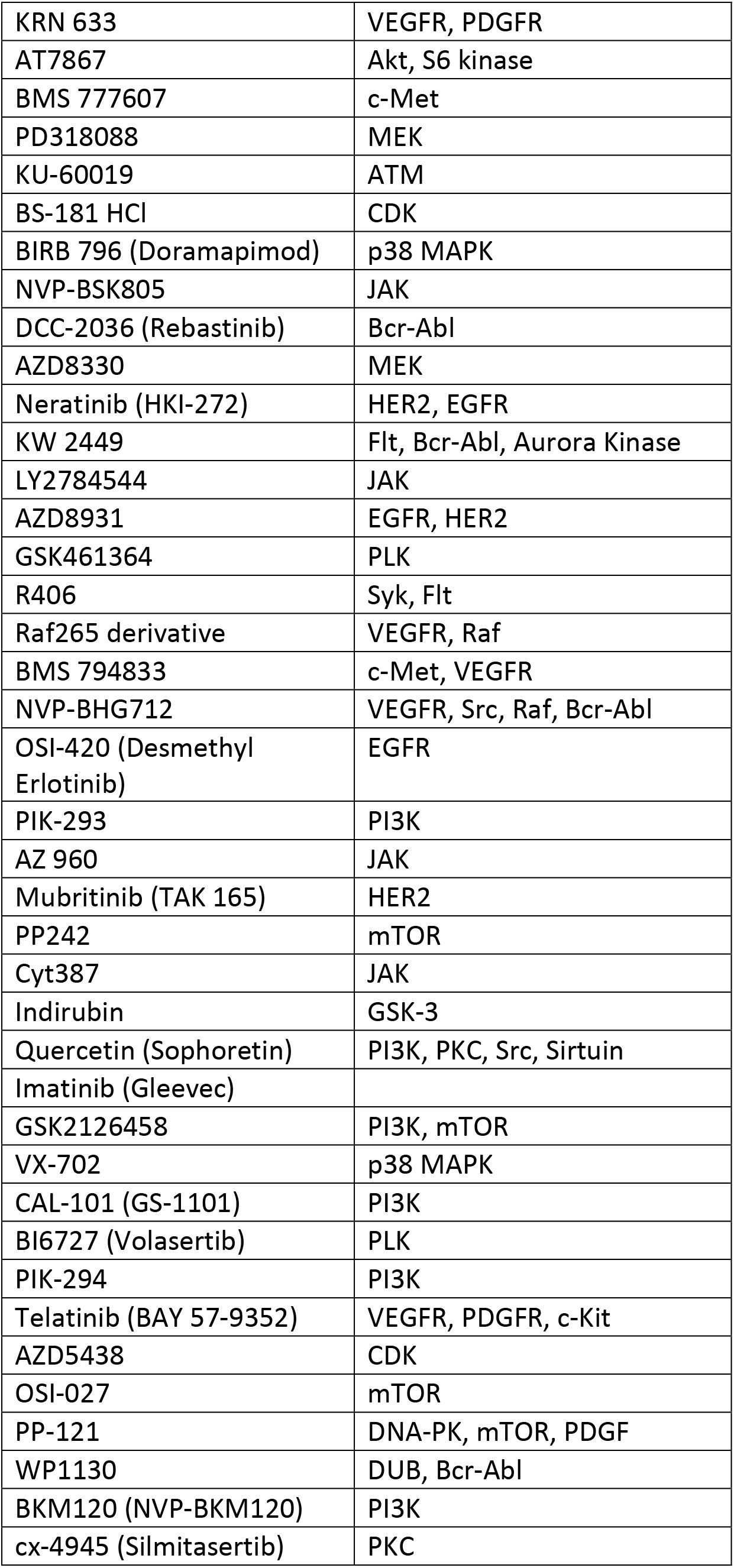
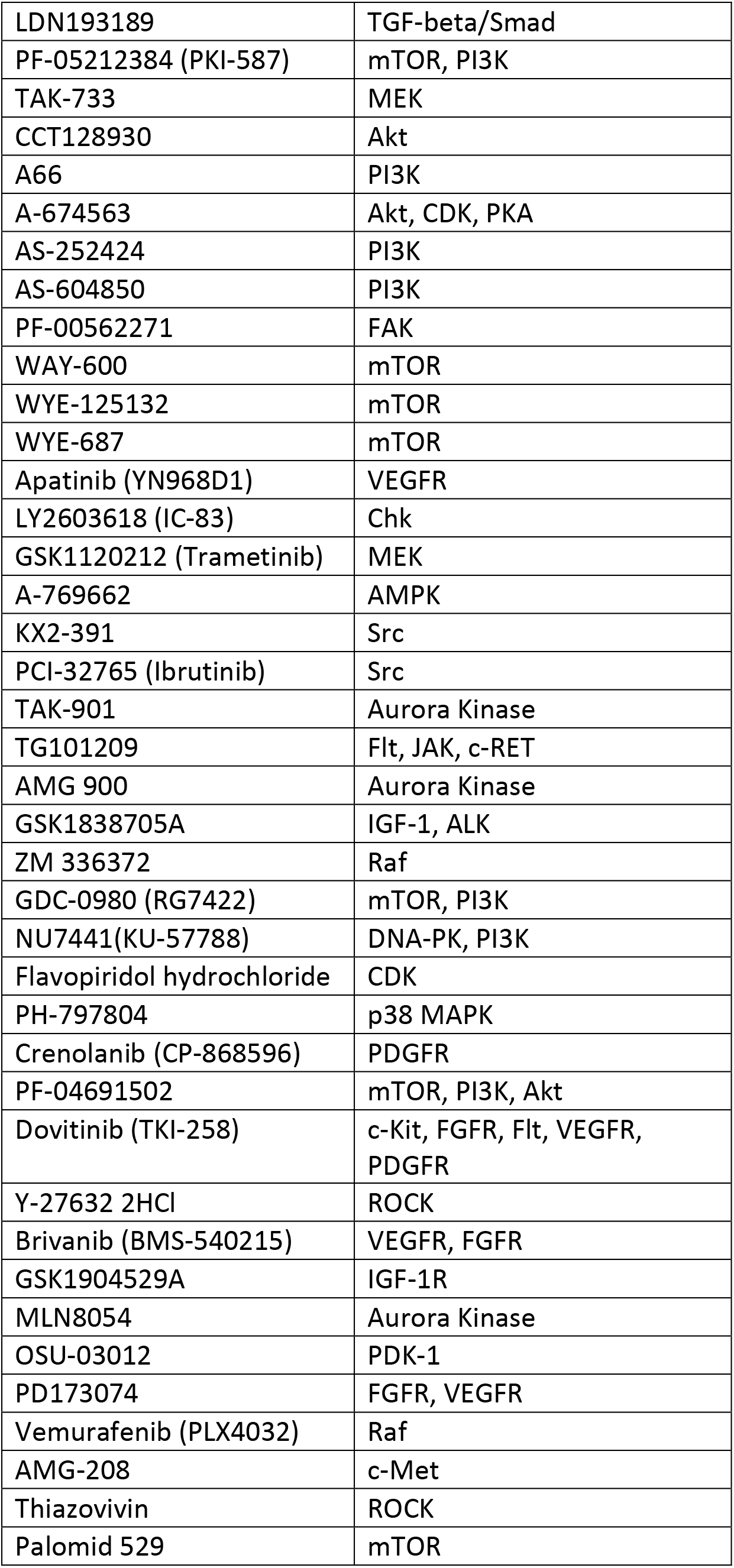
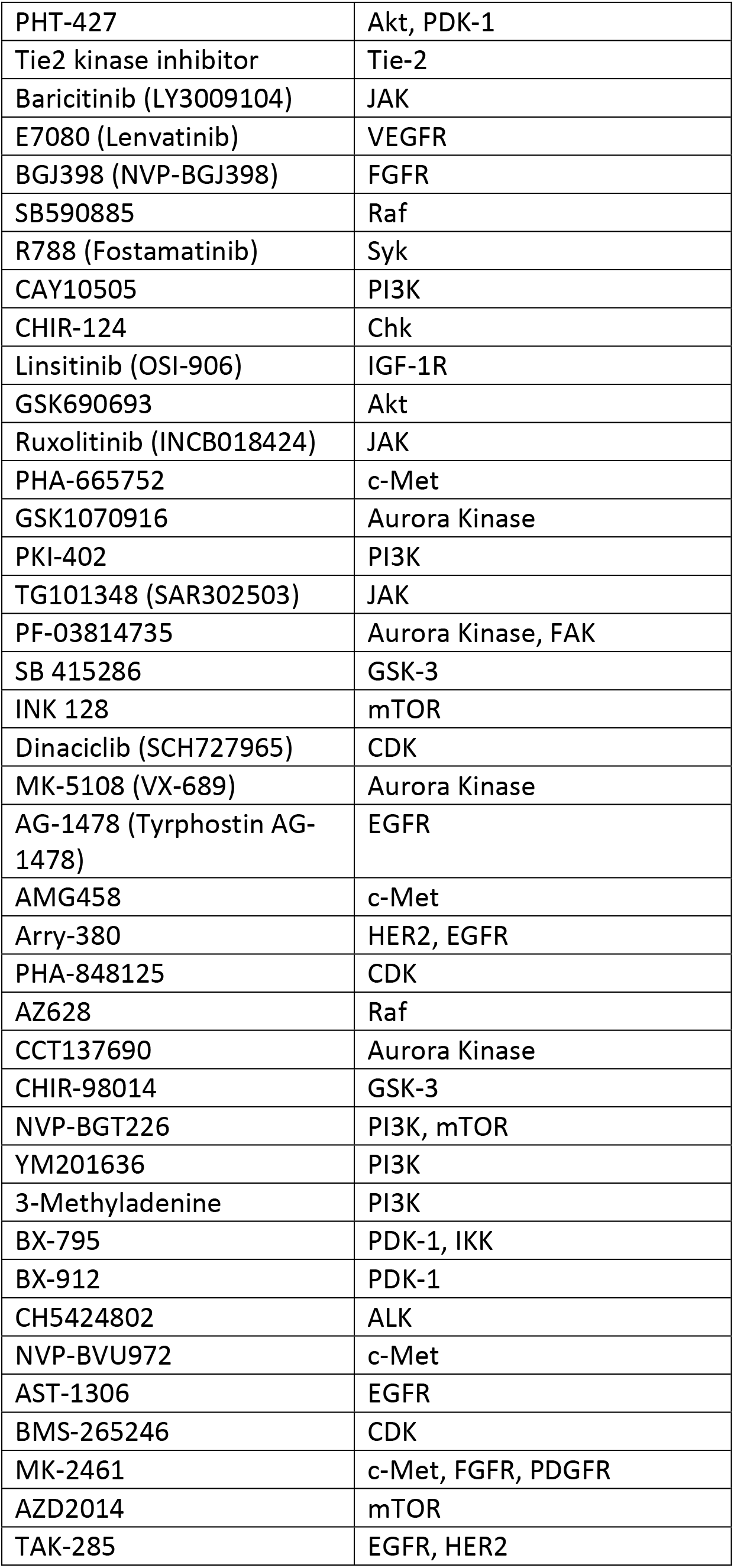
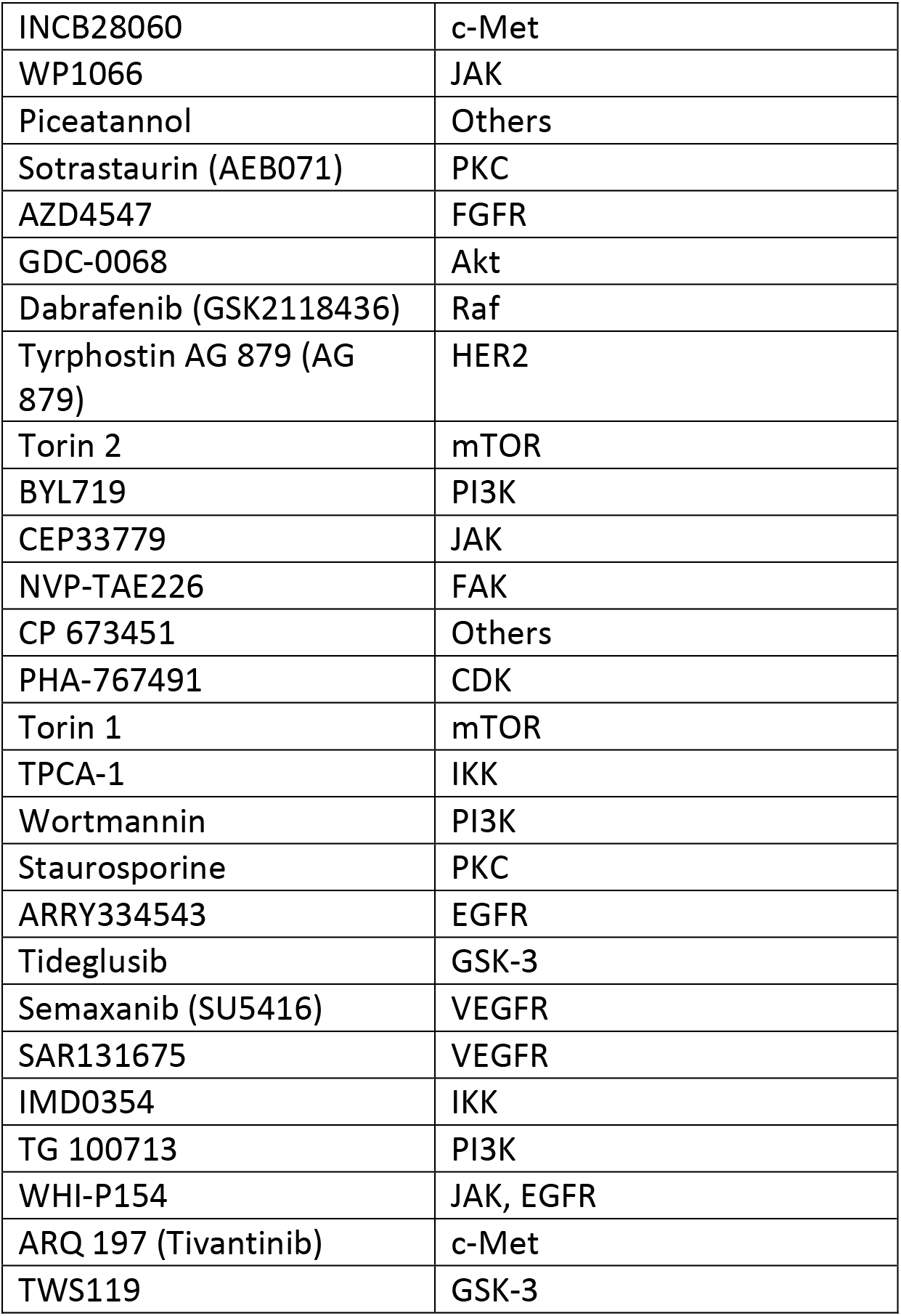
Kinase Inhibitor Library (Selleck Chemicals) and targets.

**Table S2:**
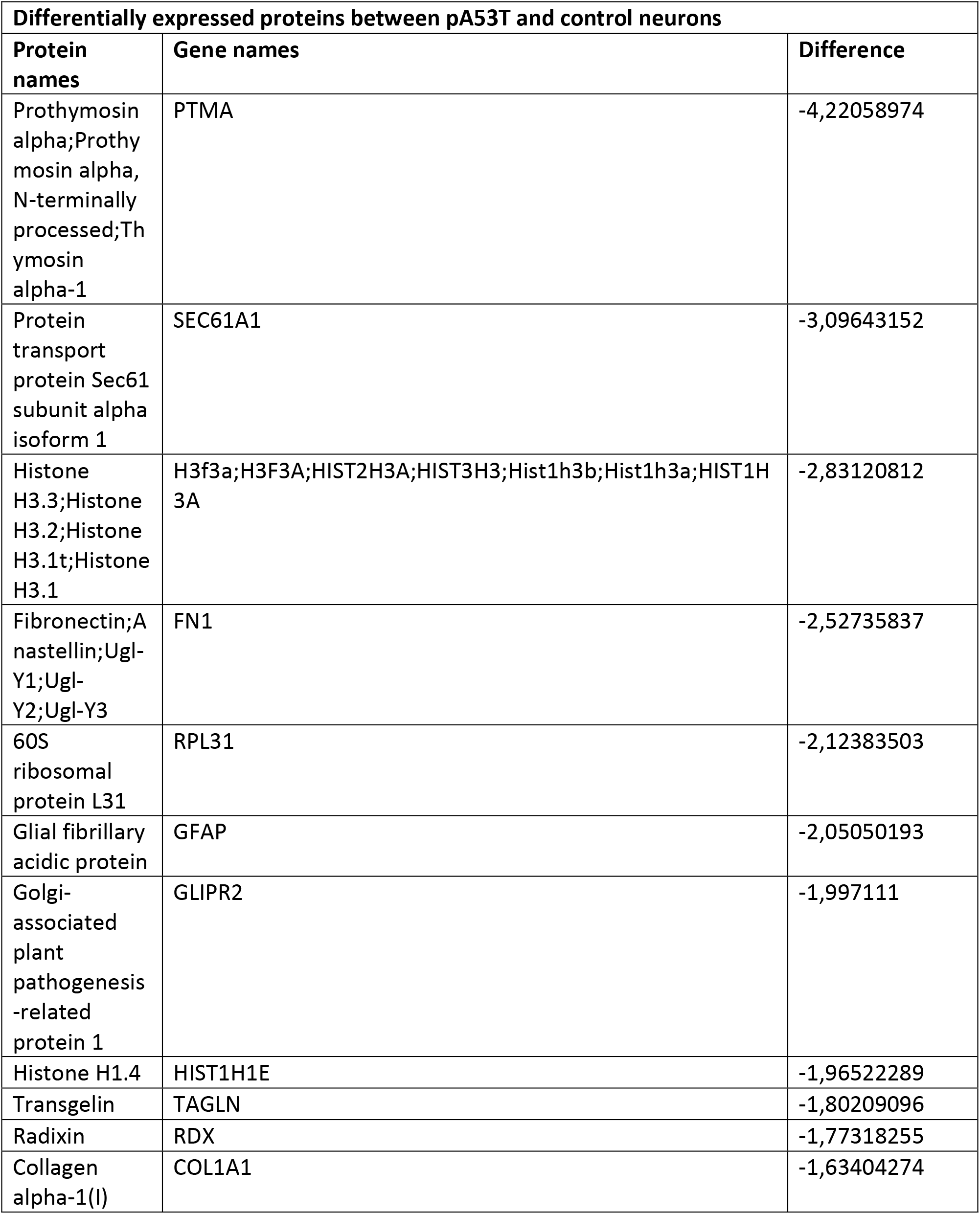
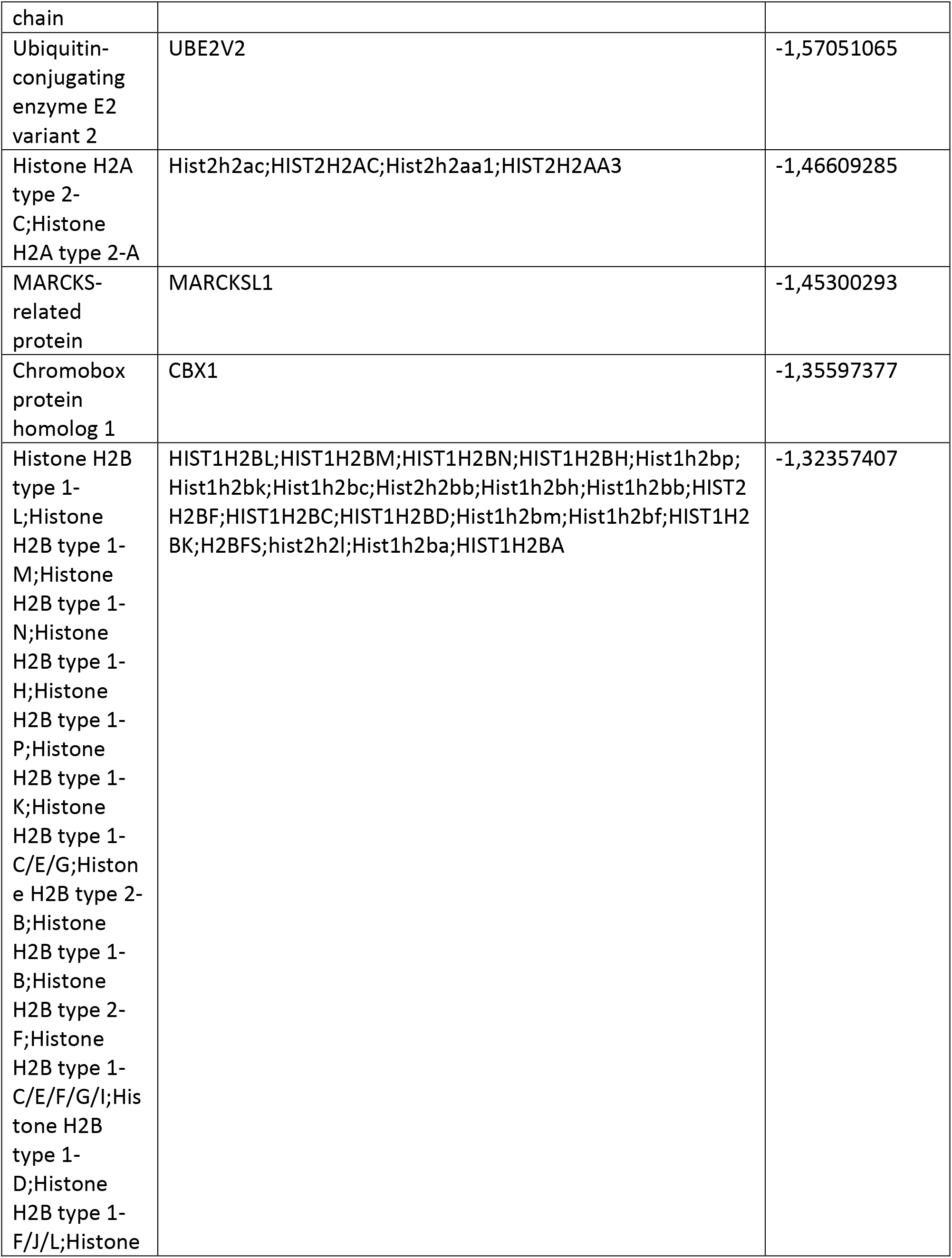
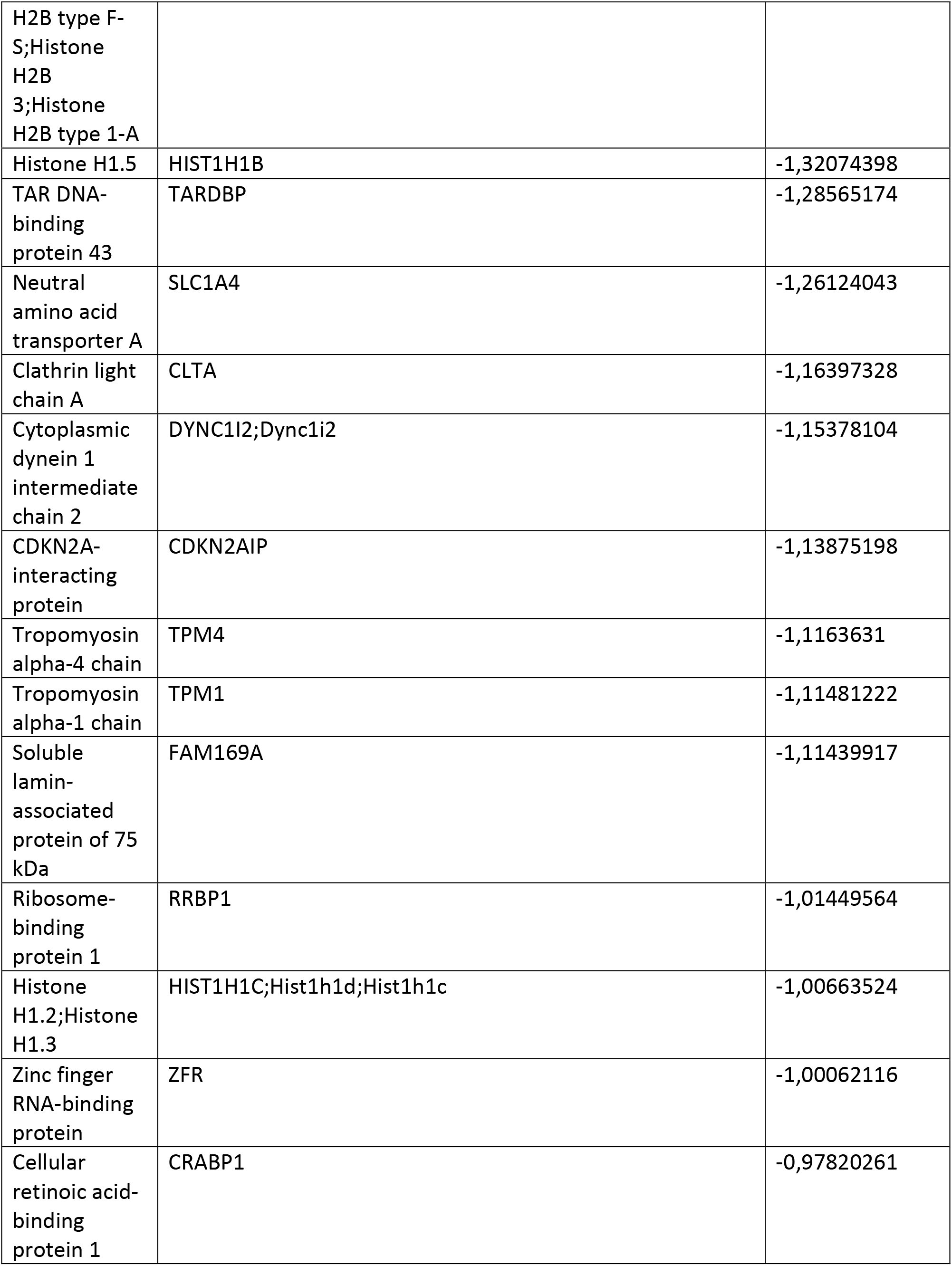
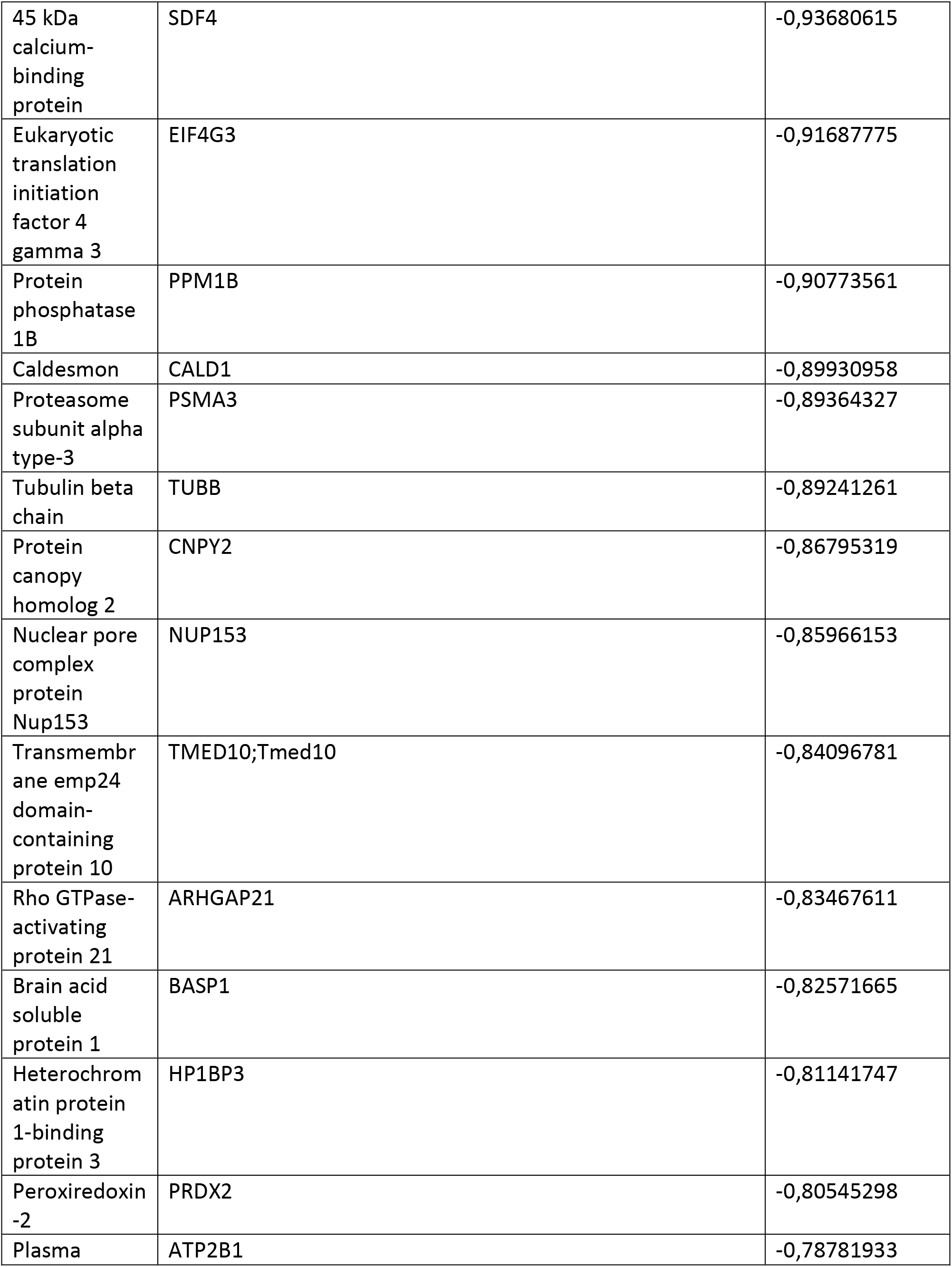
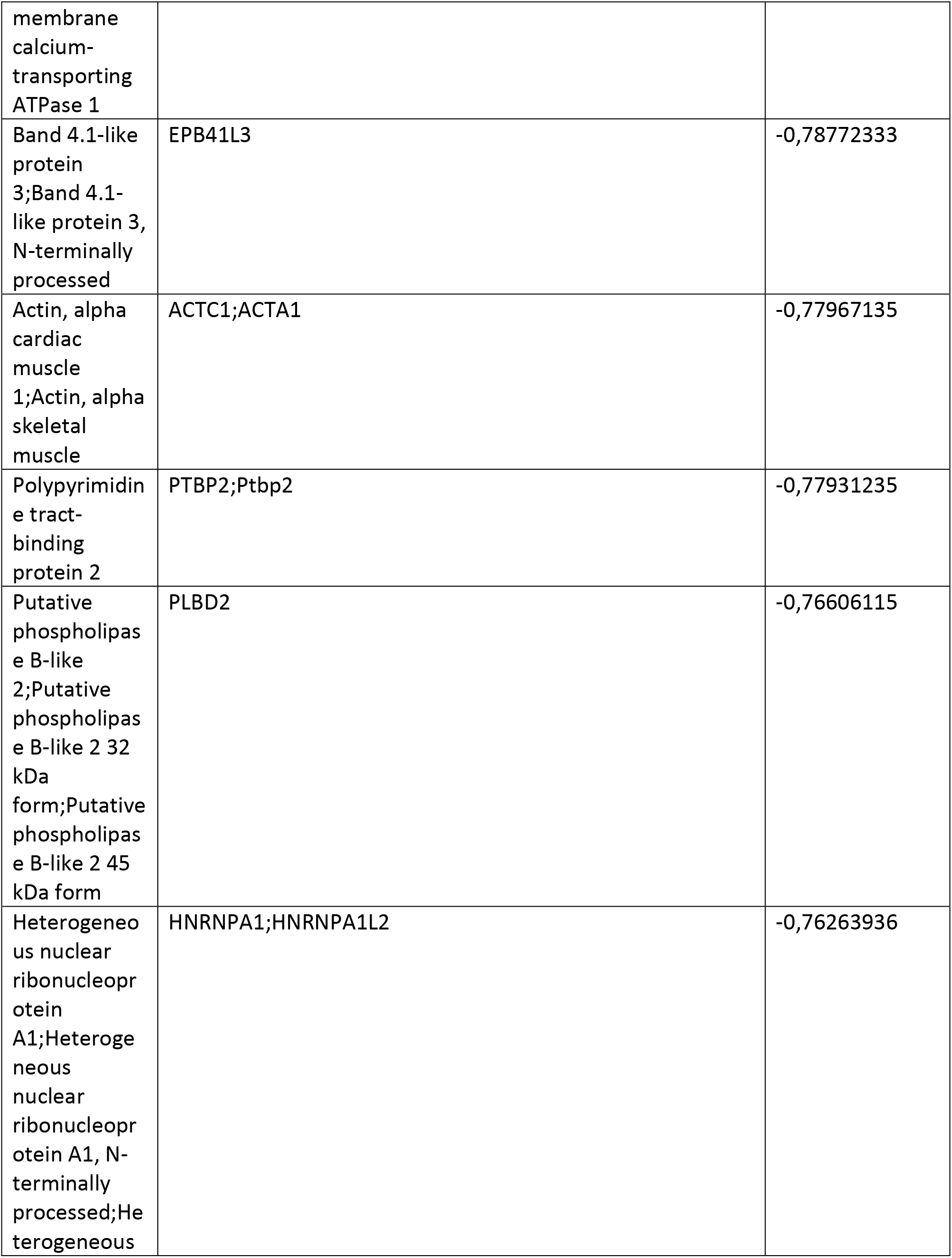
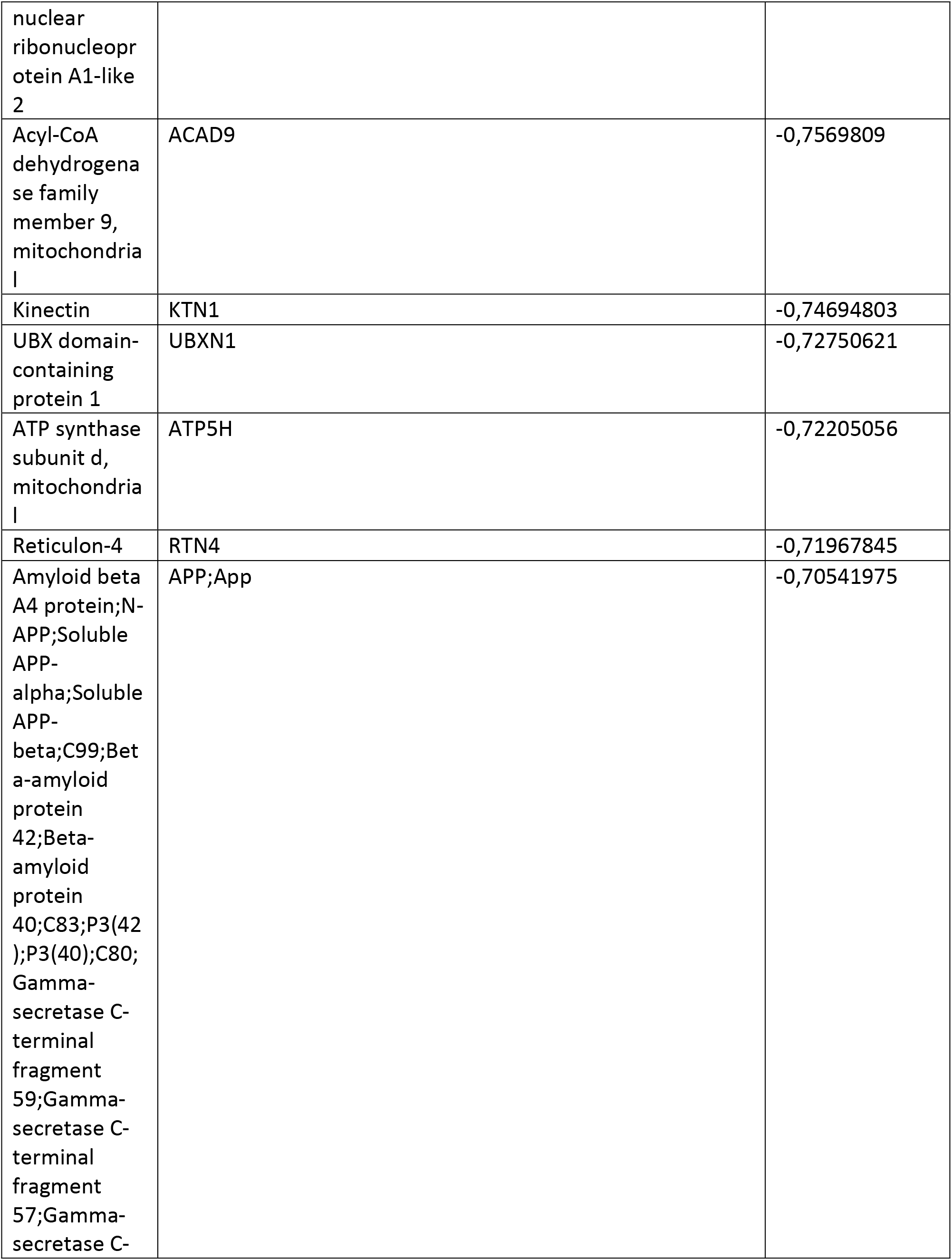
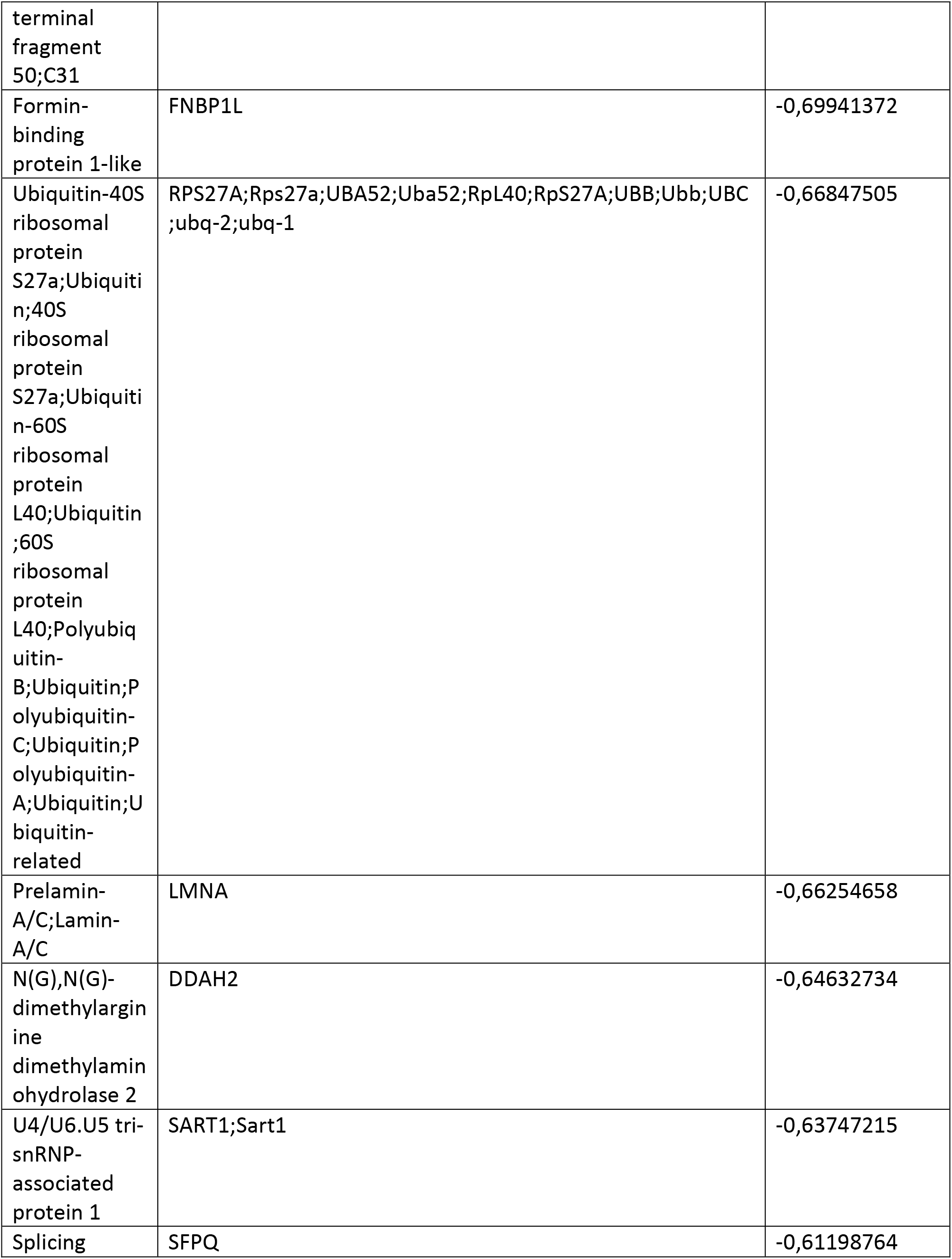
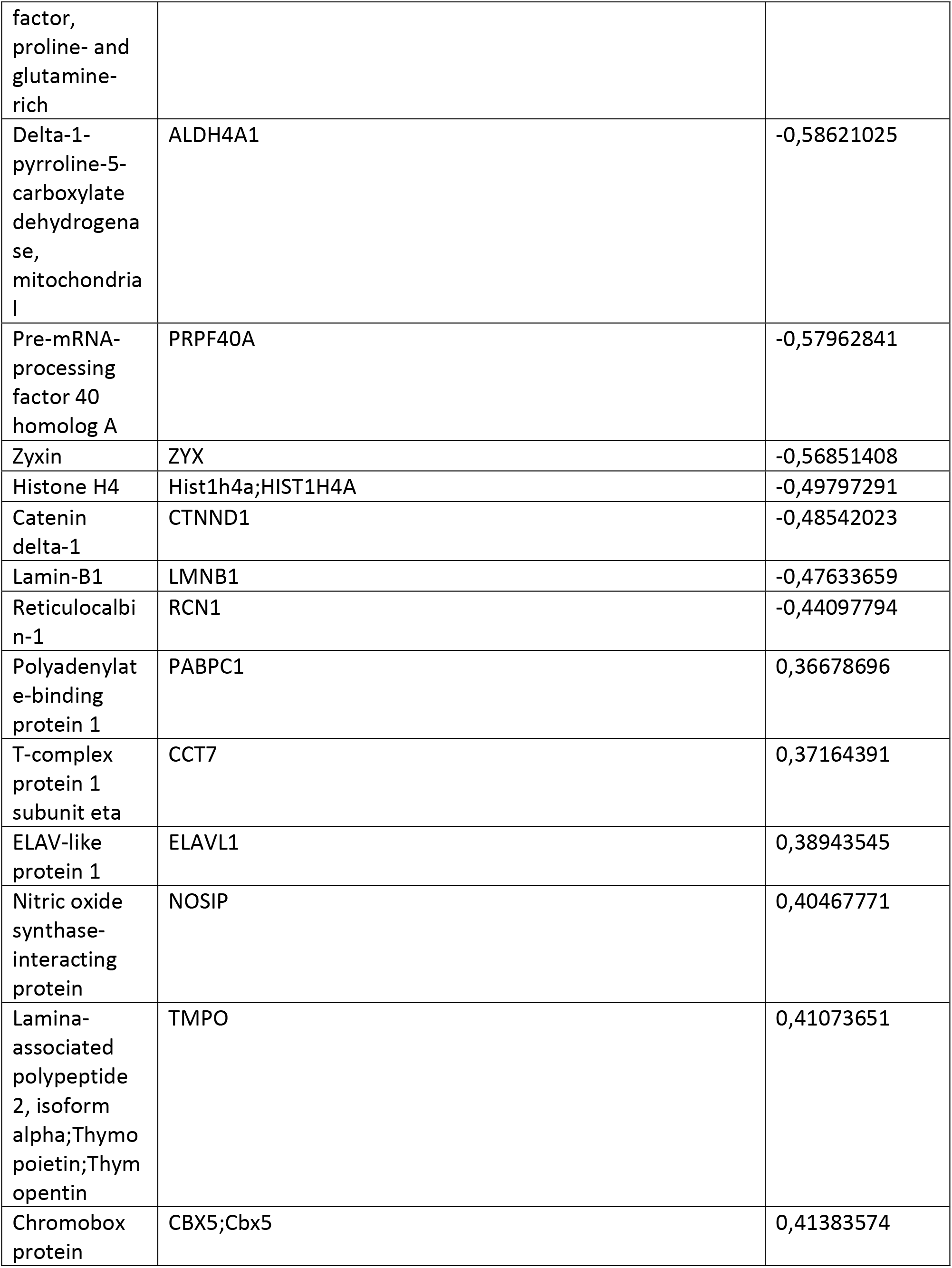
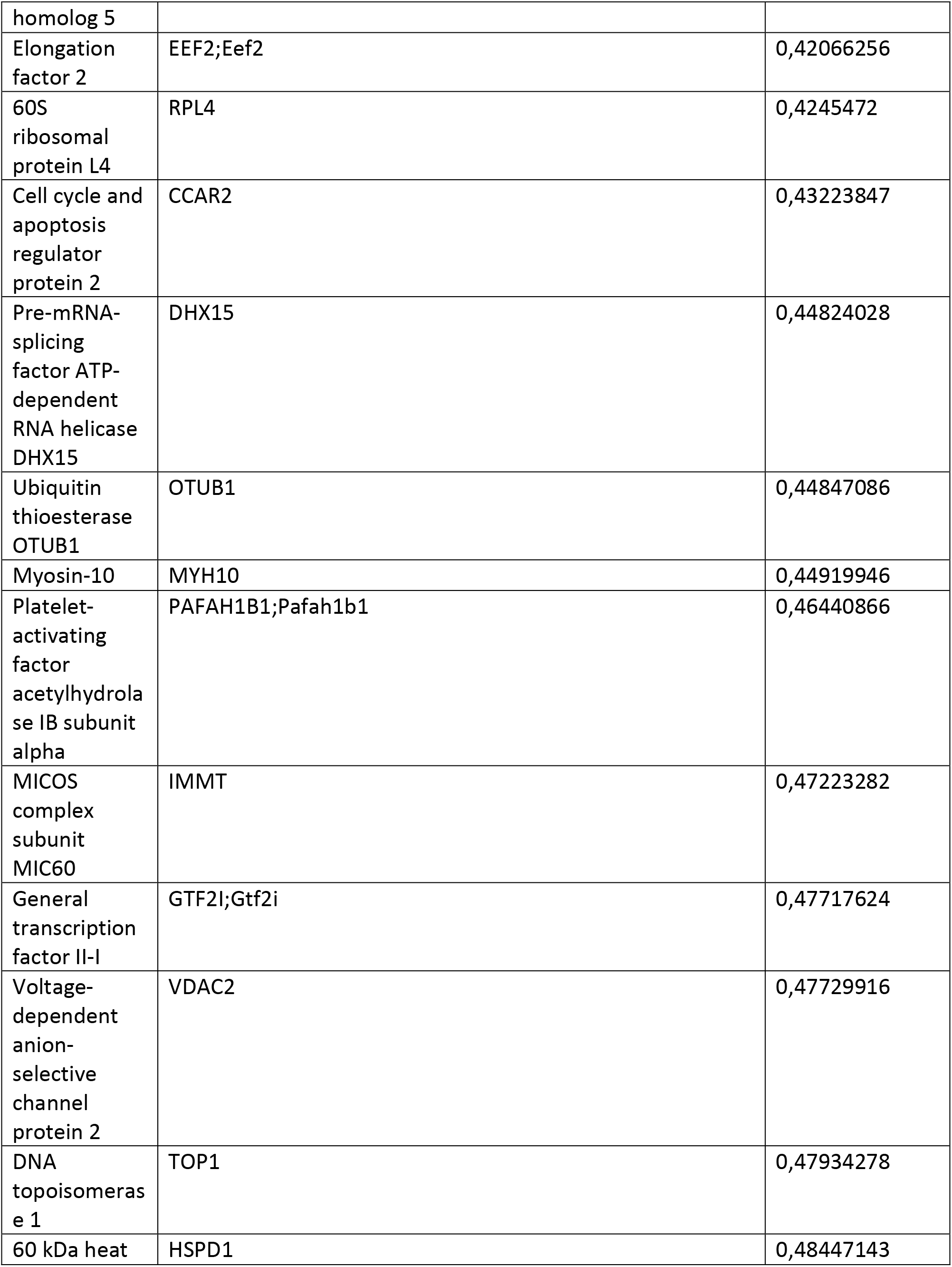
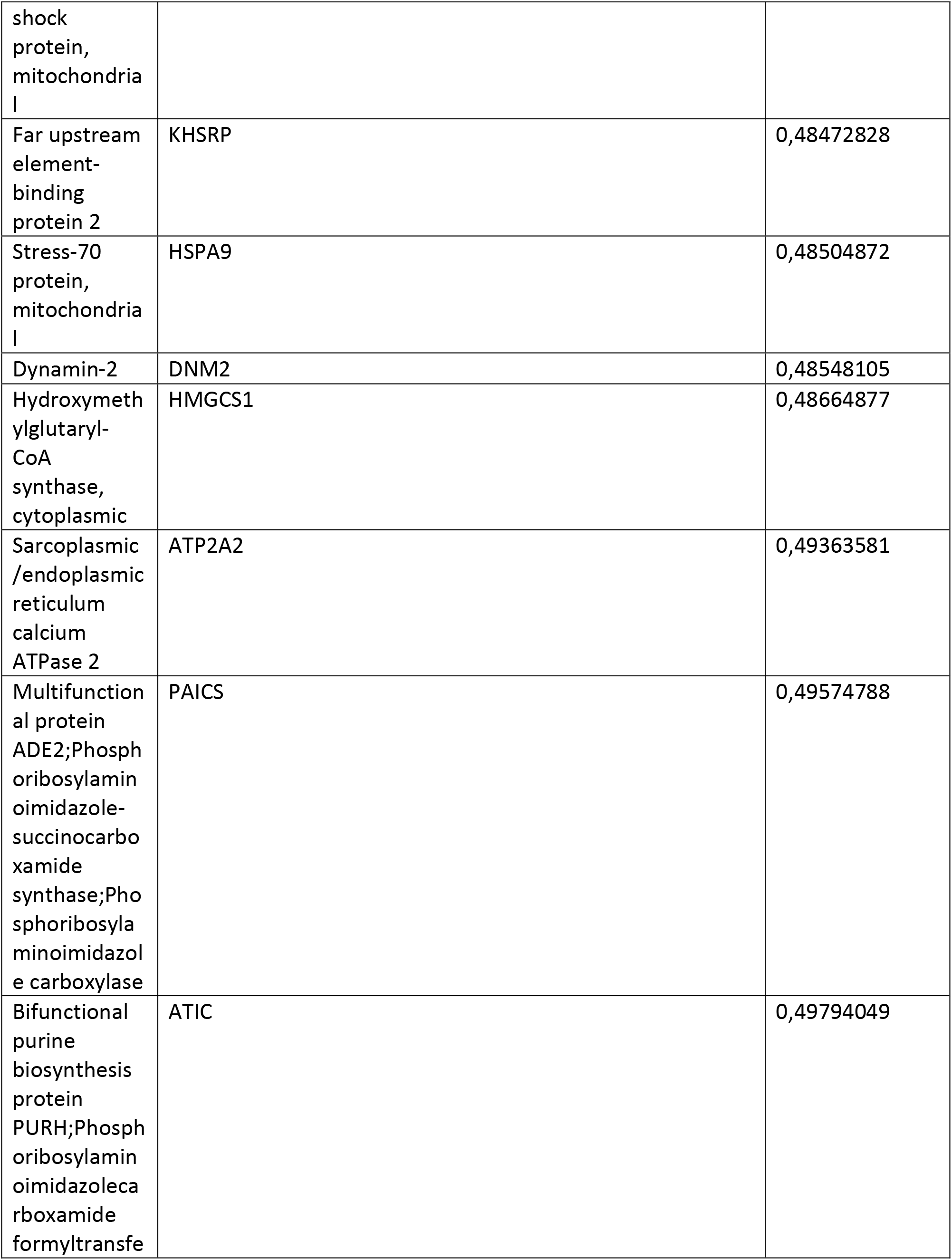
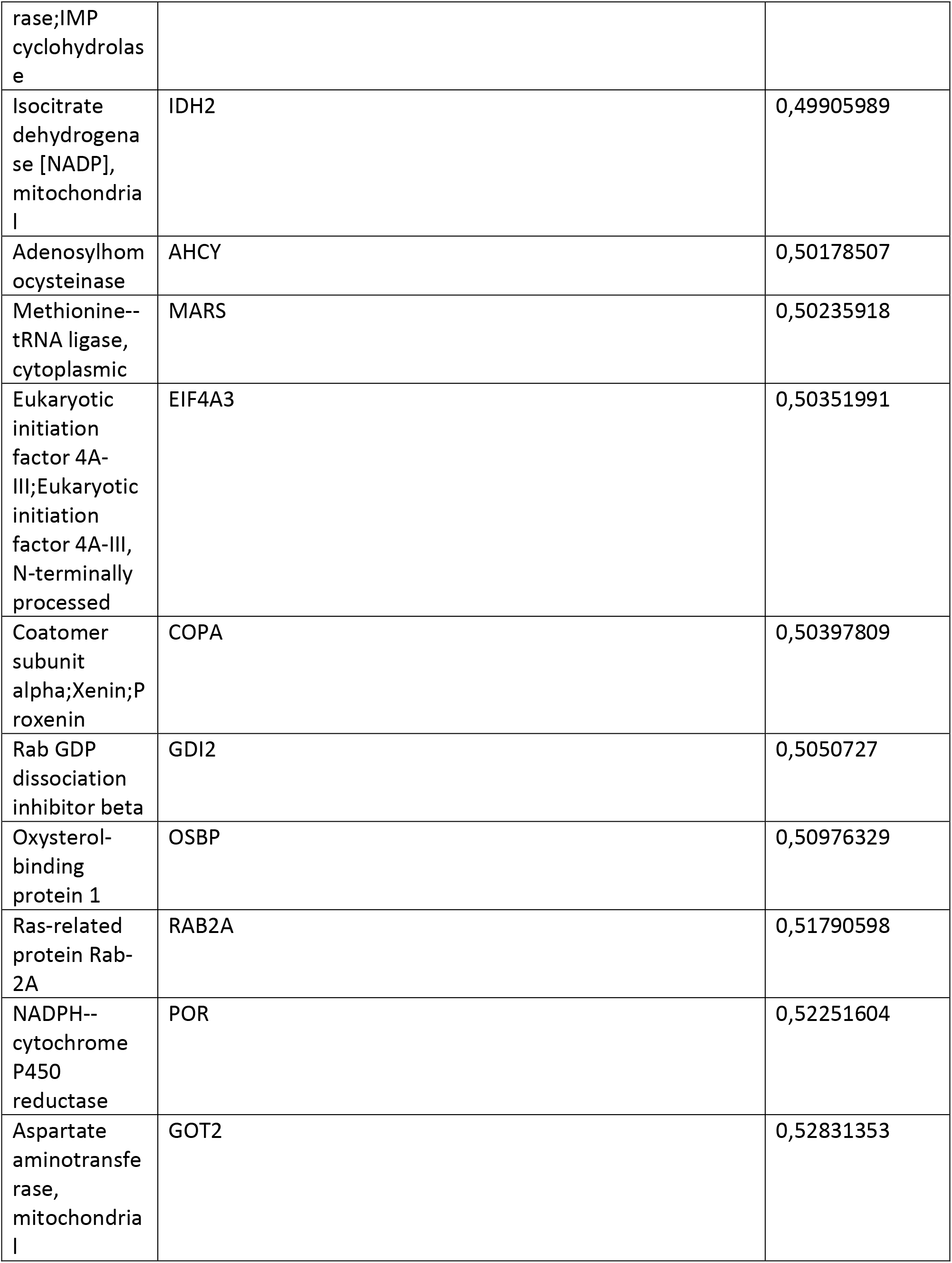
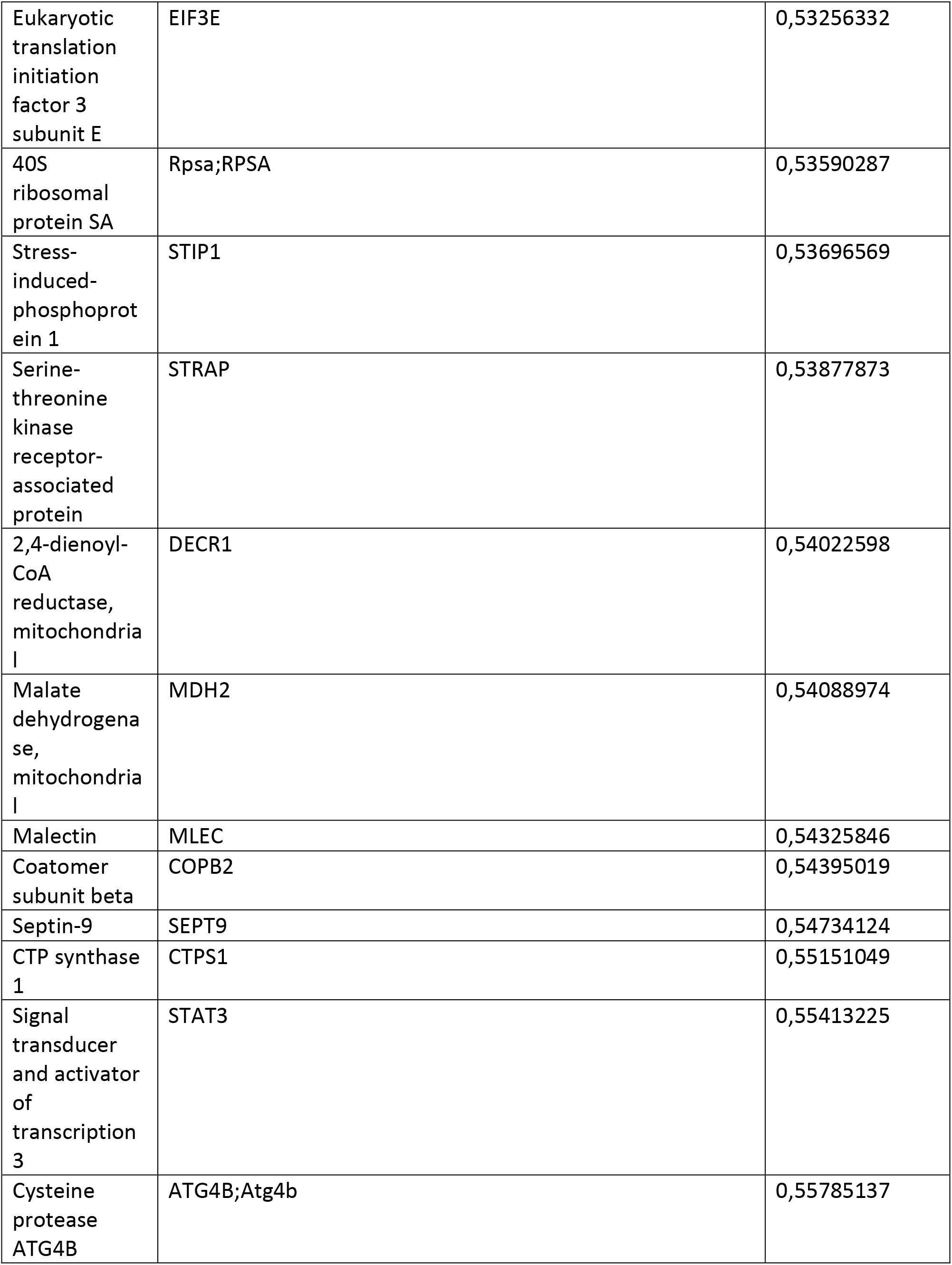
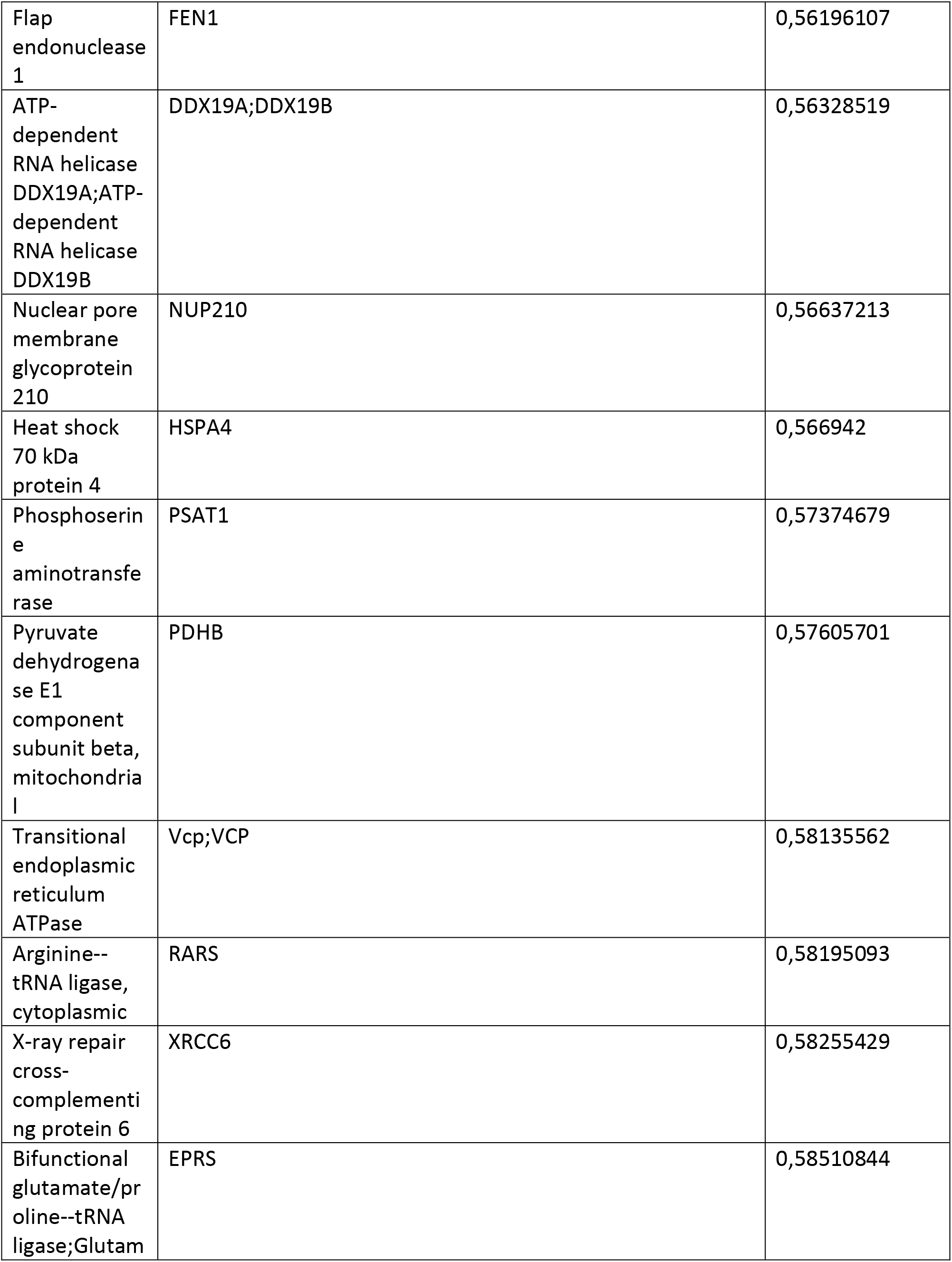
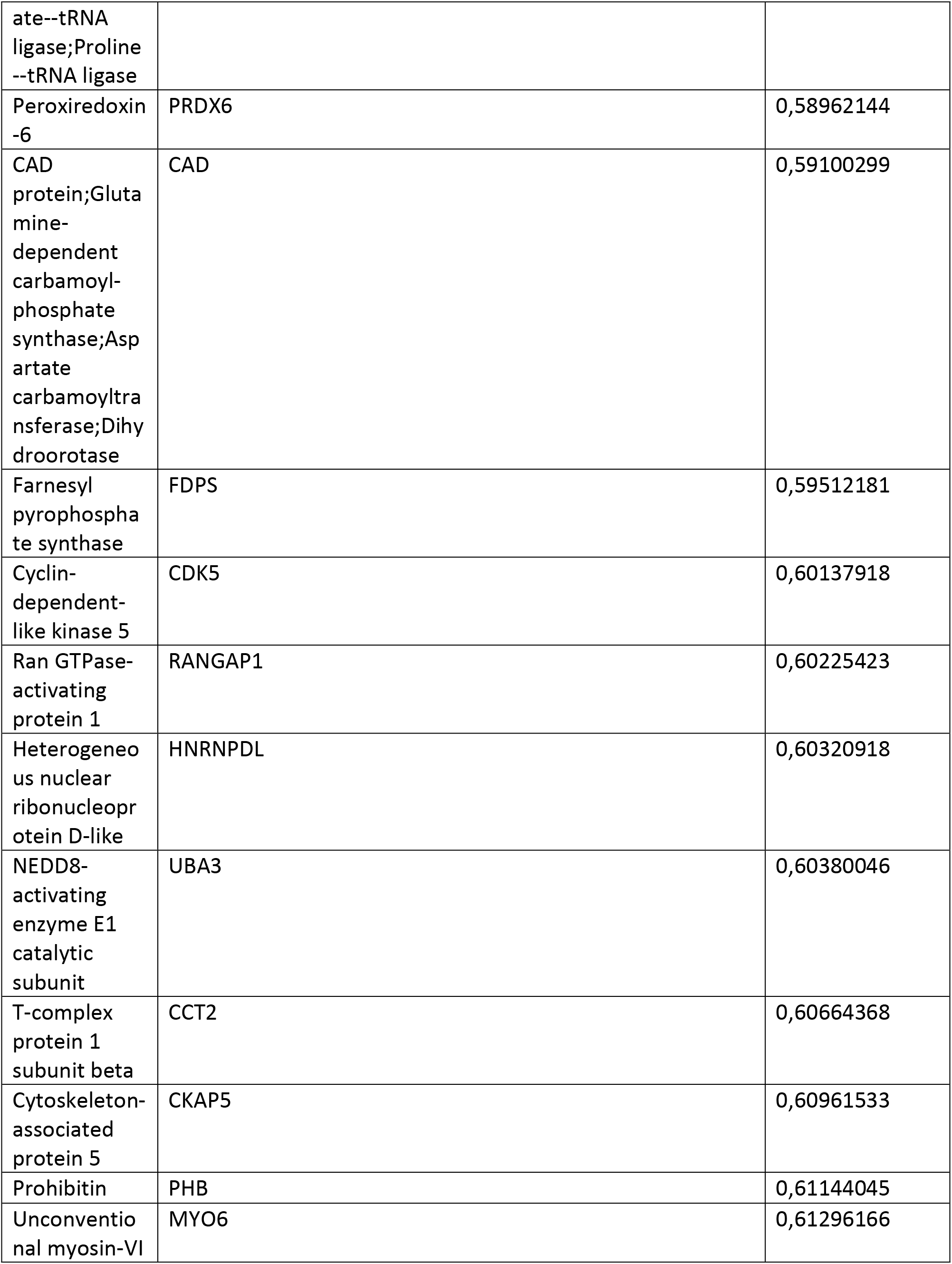
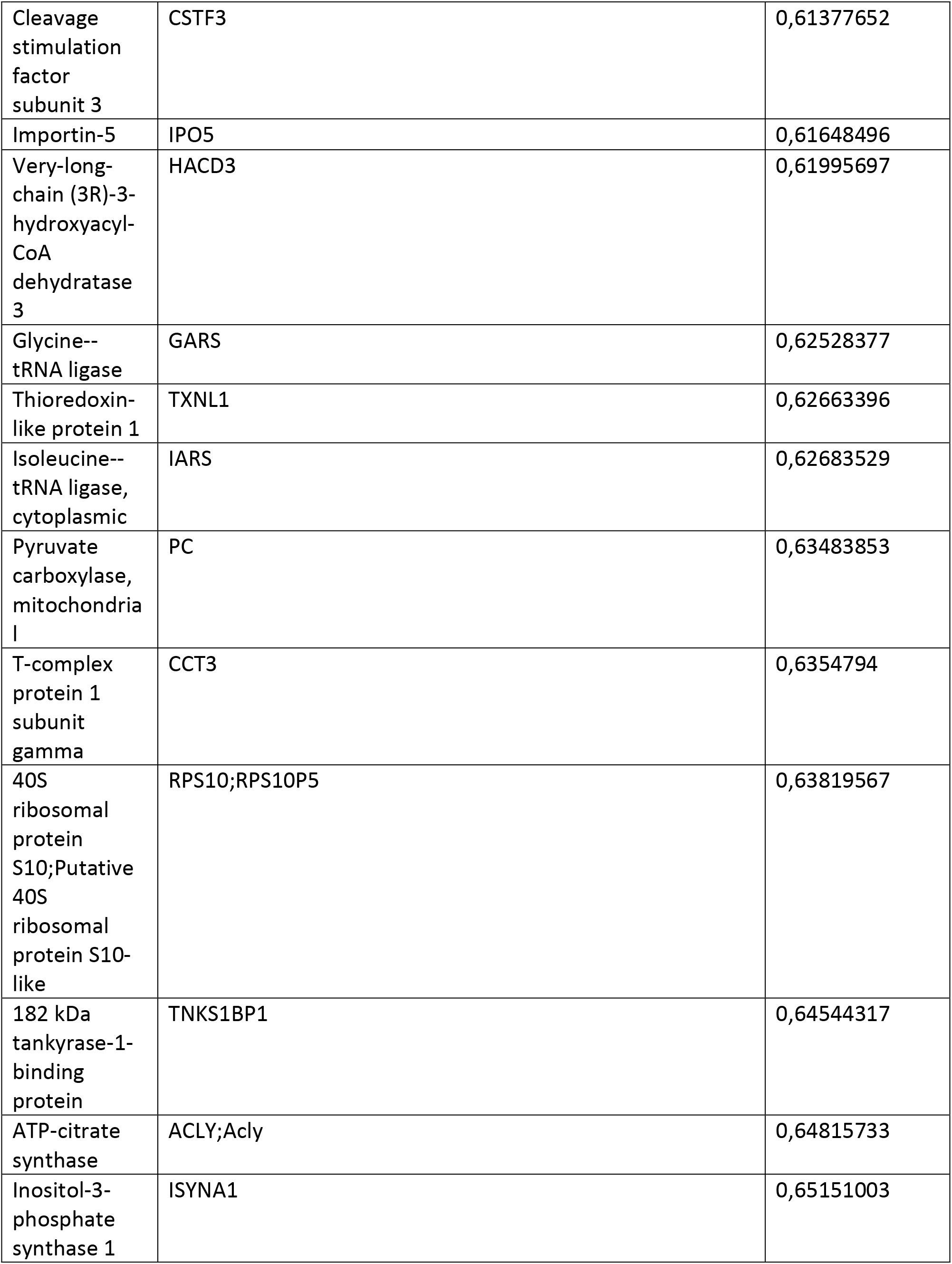
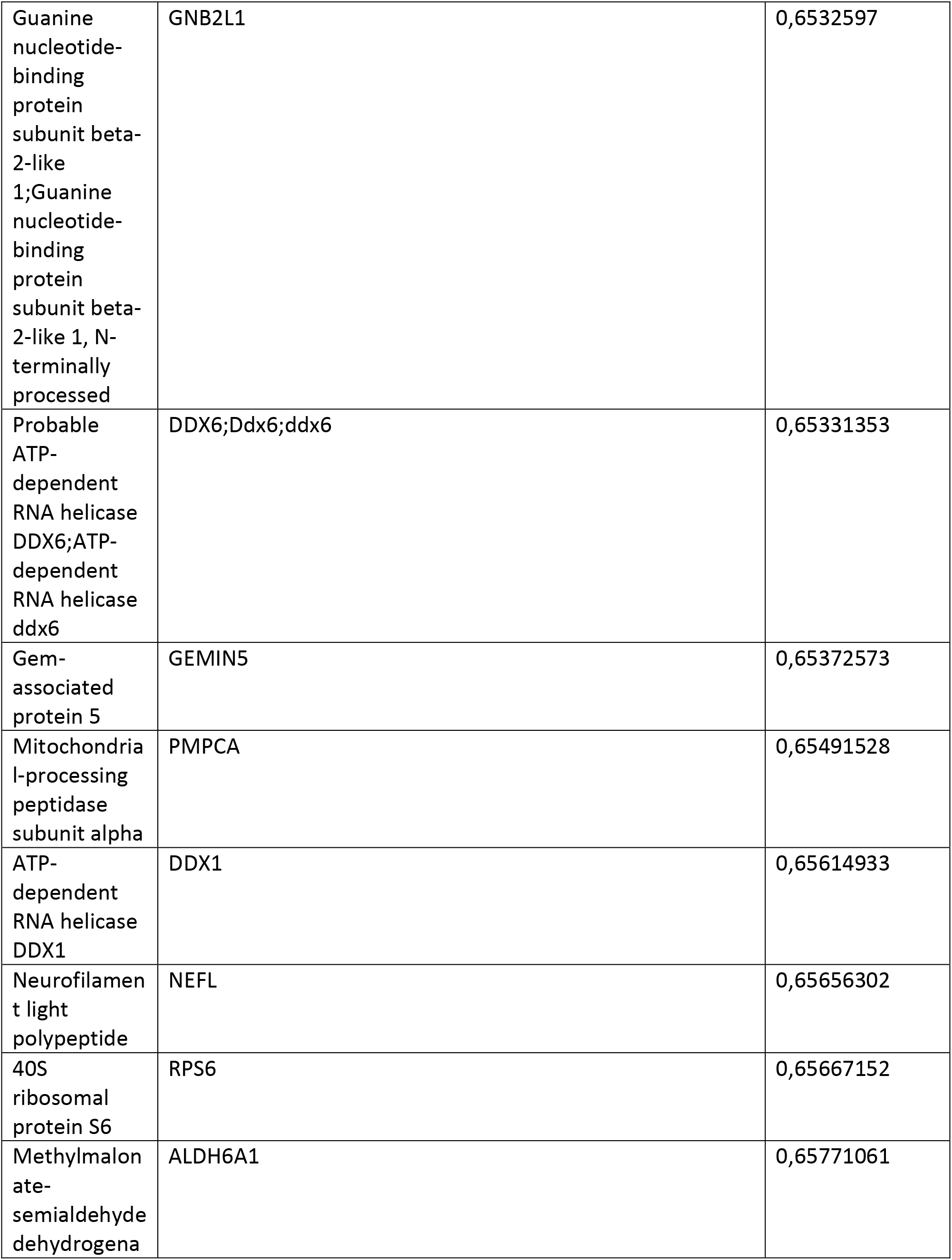
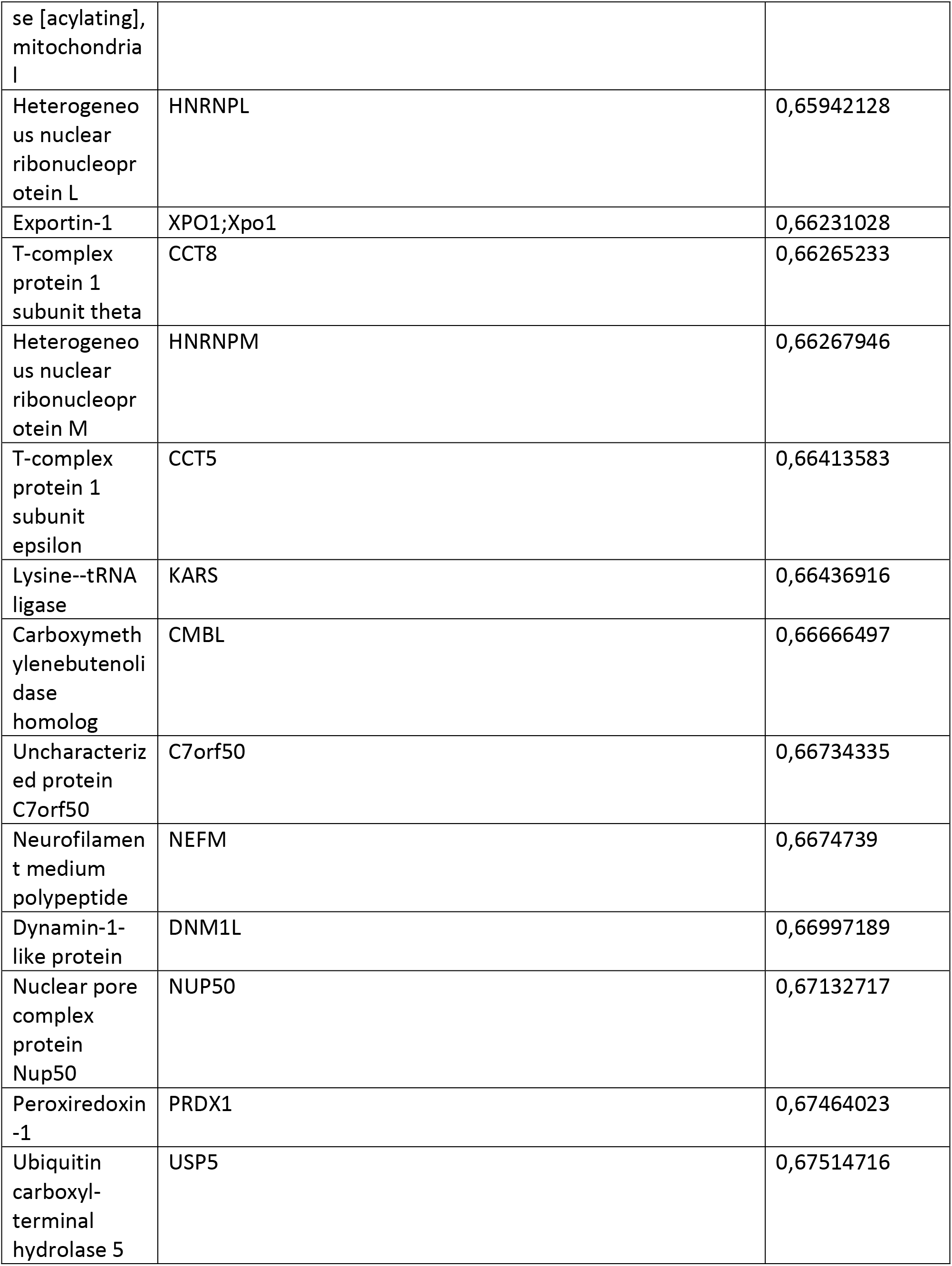
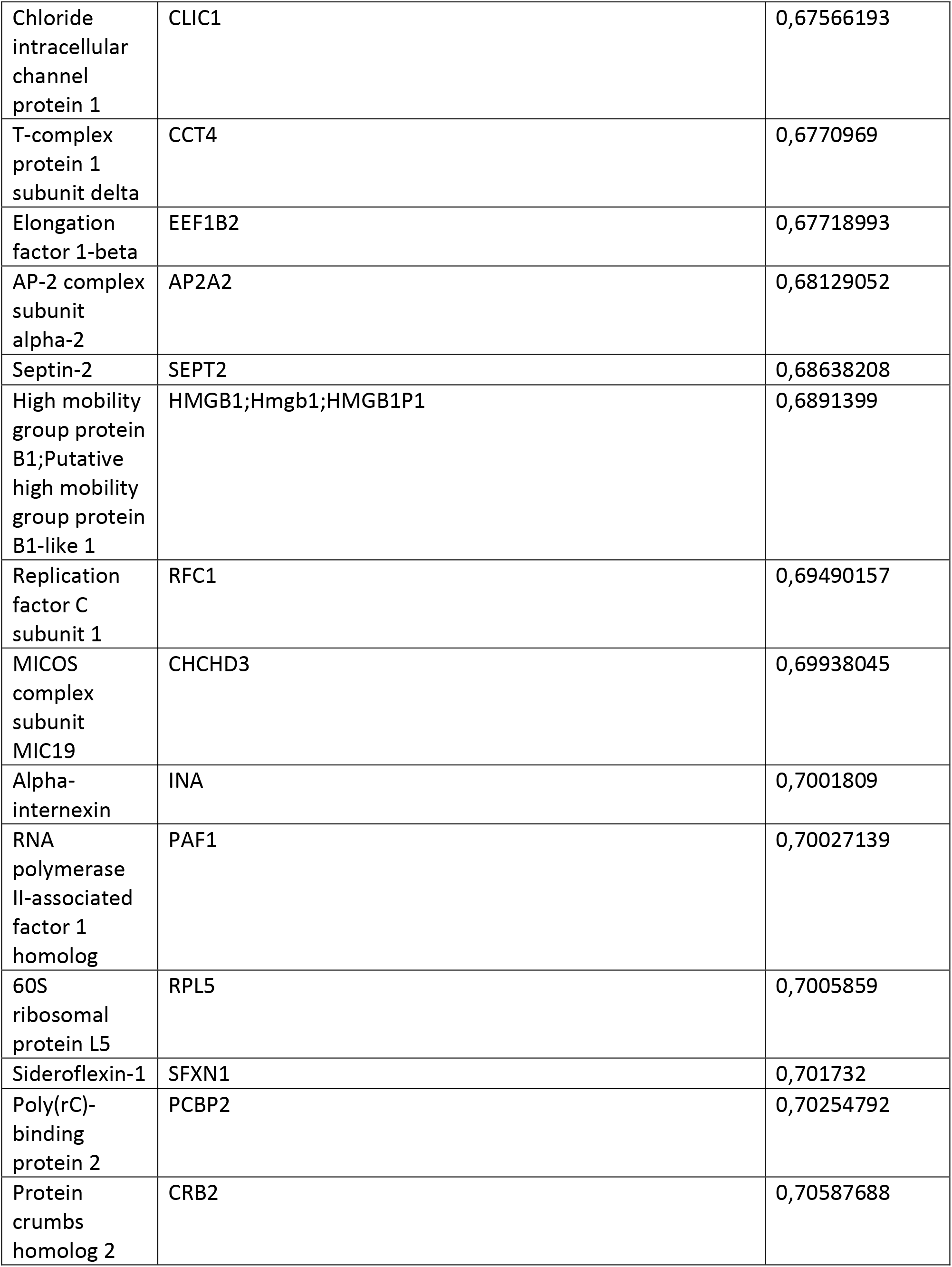
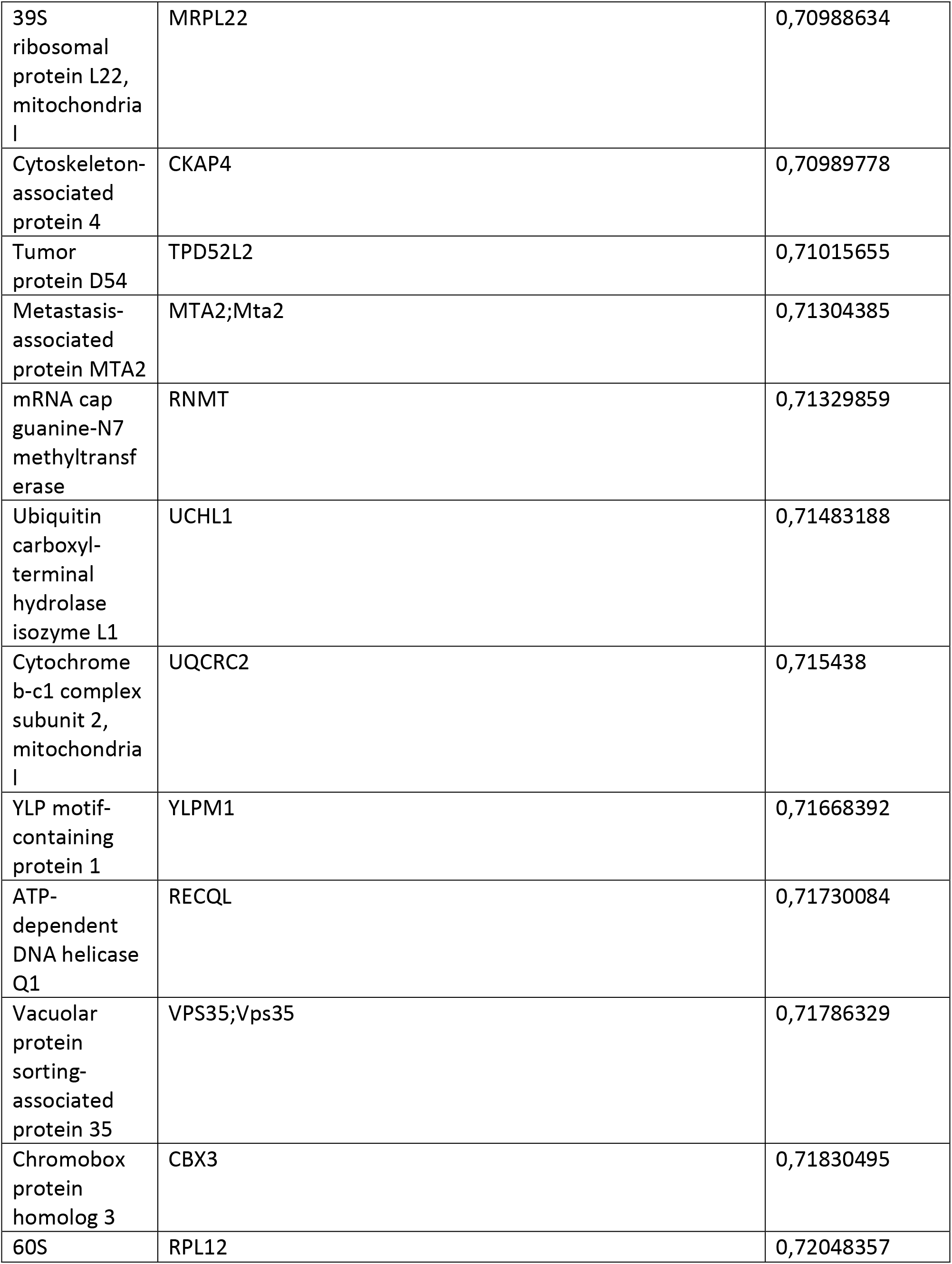
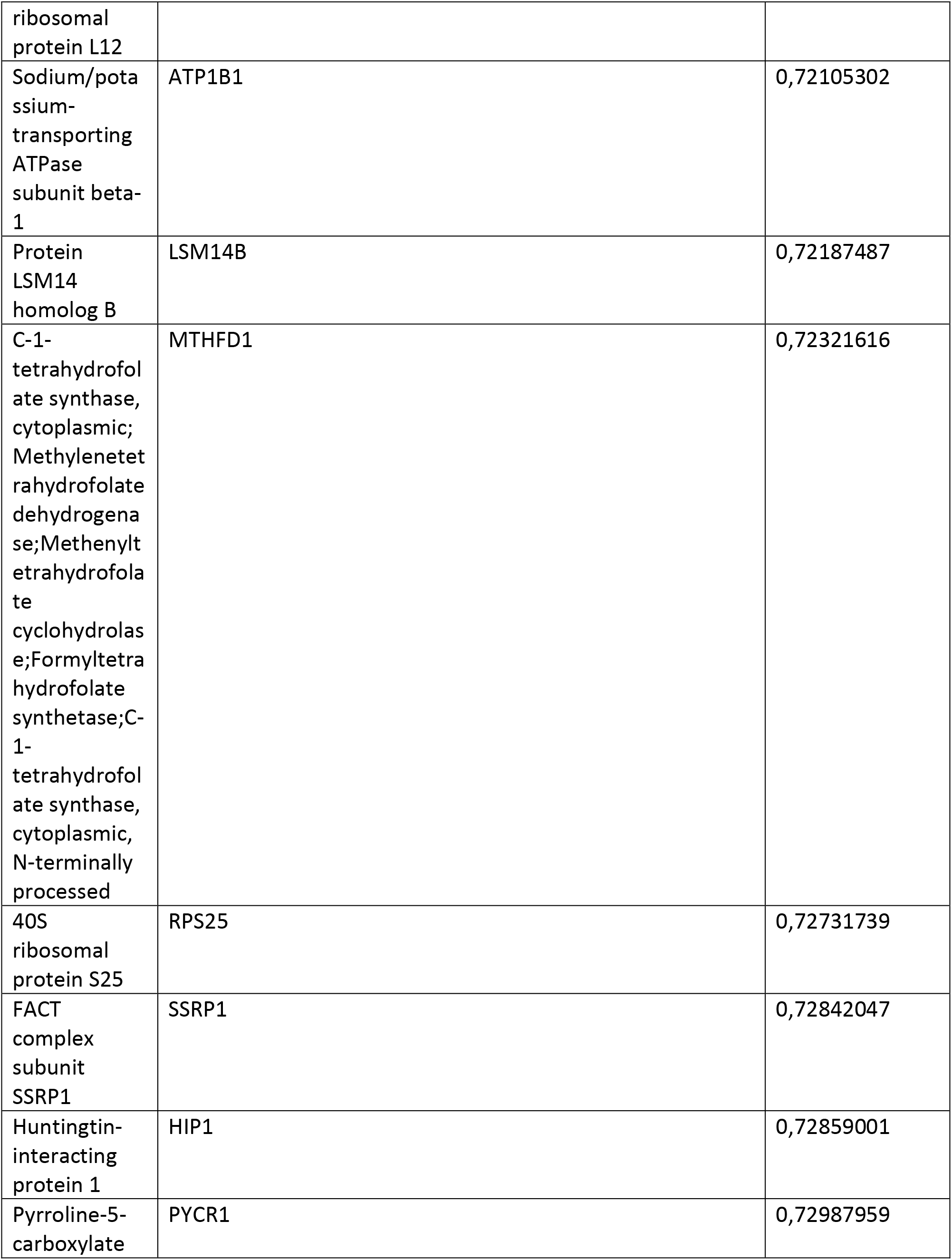
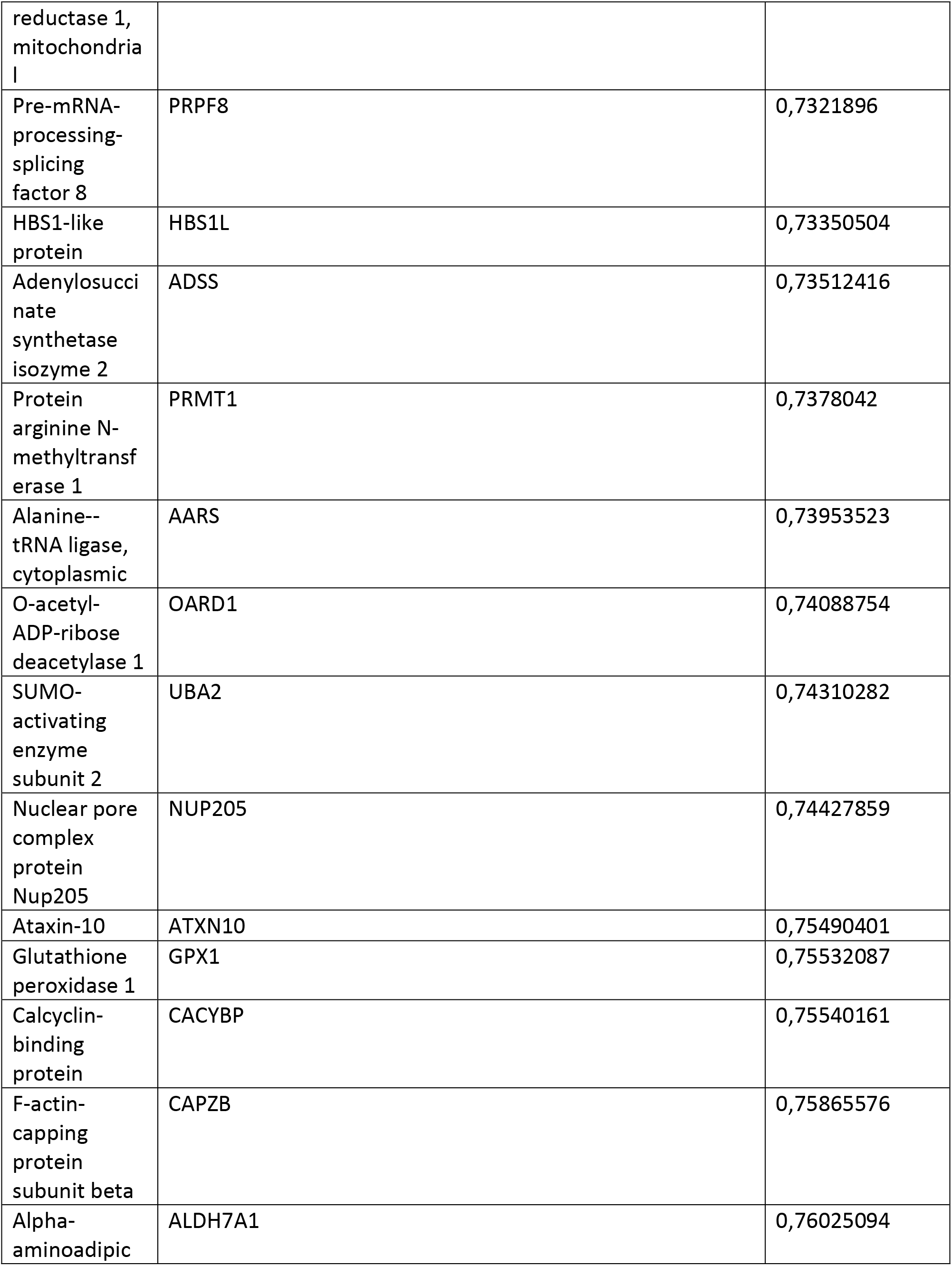
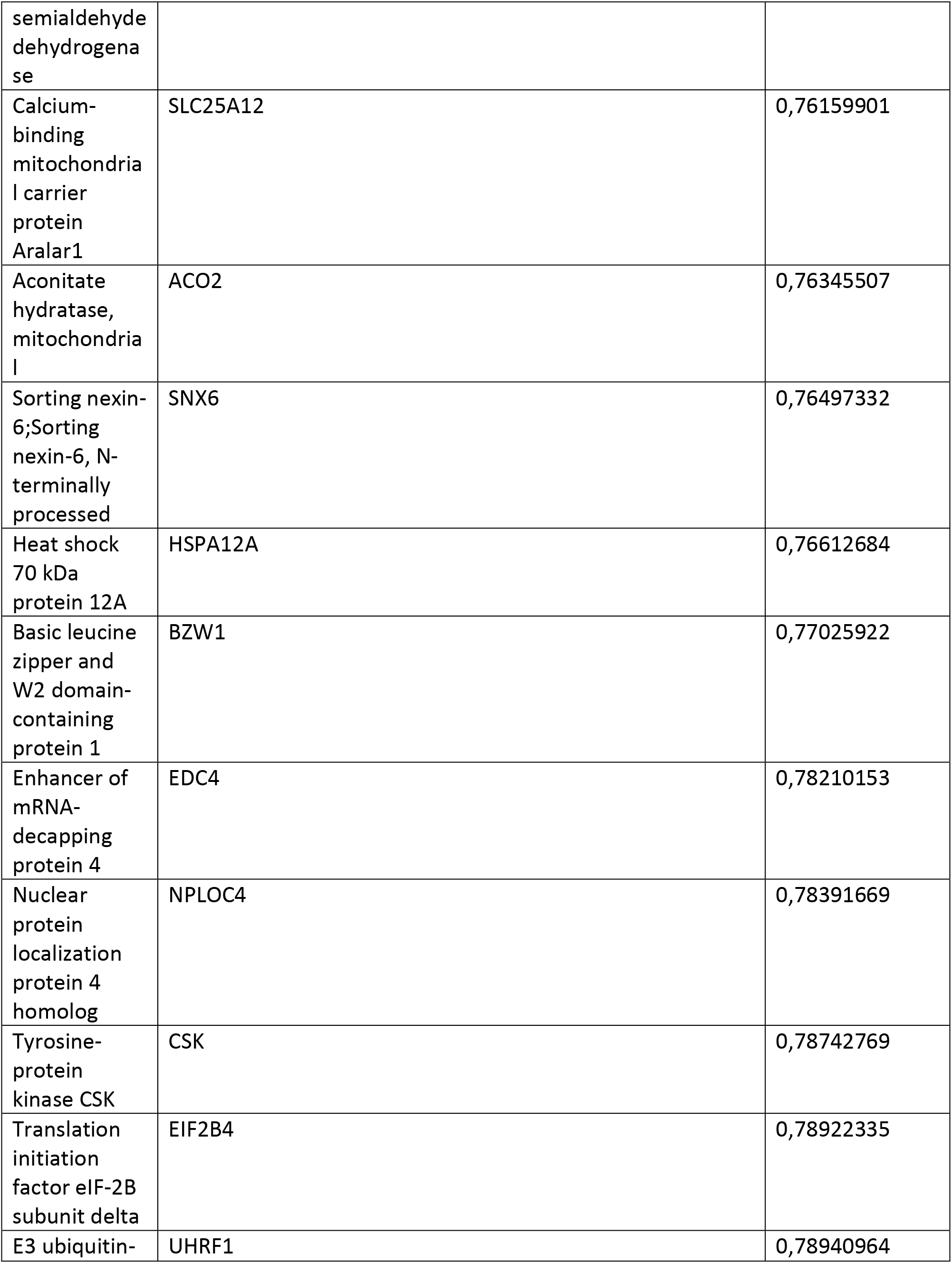
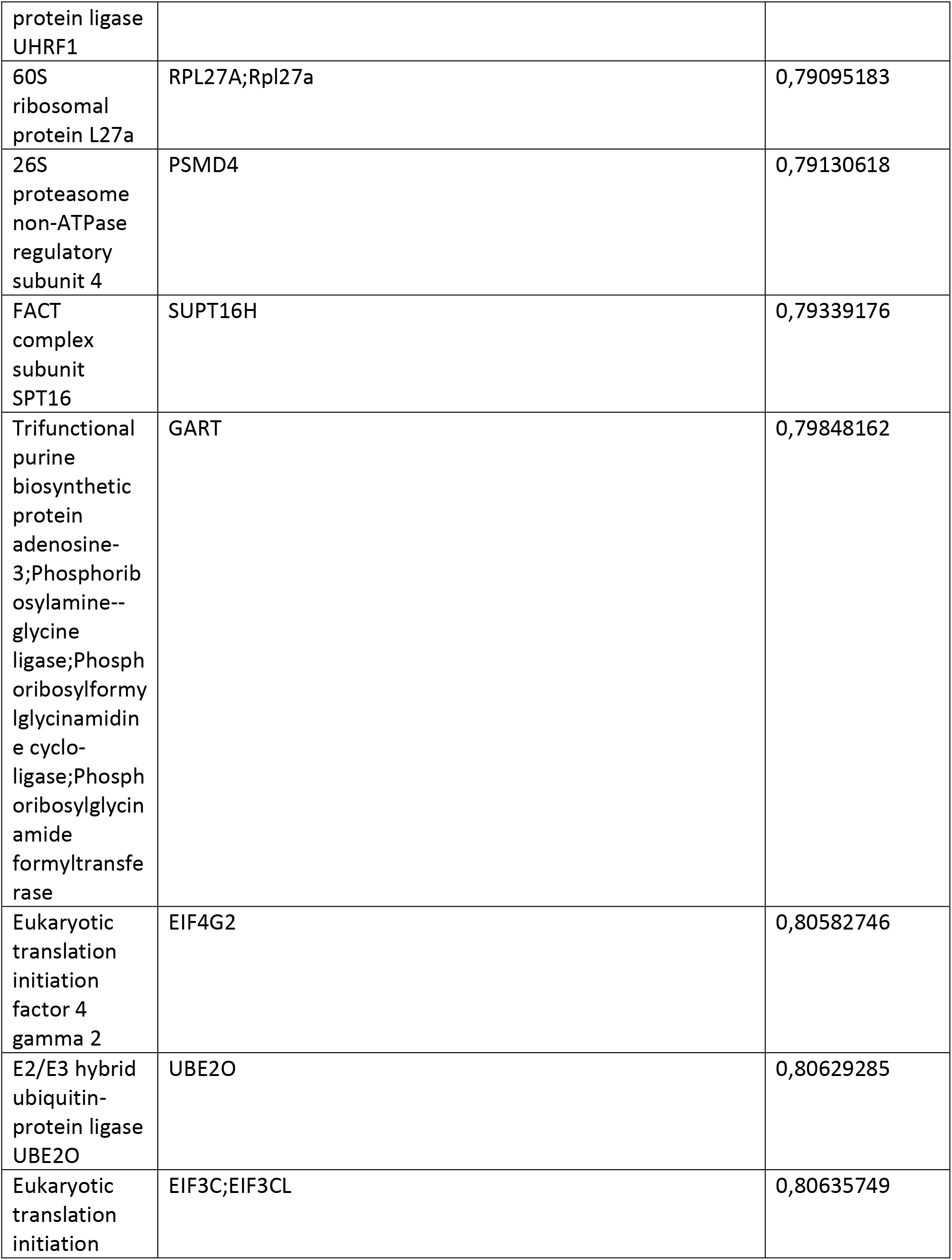
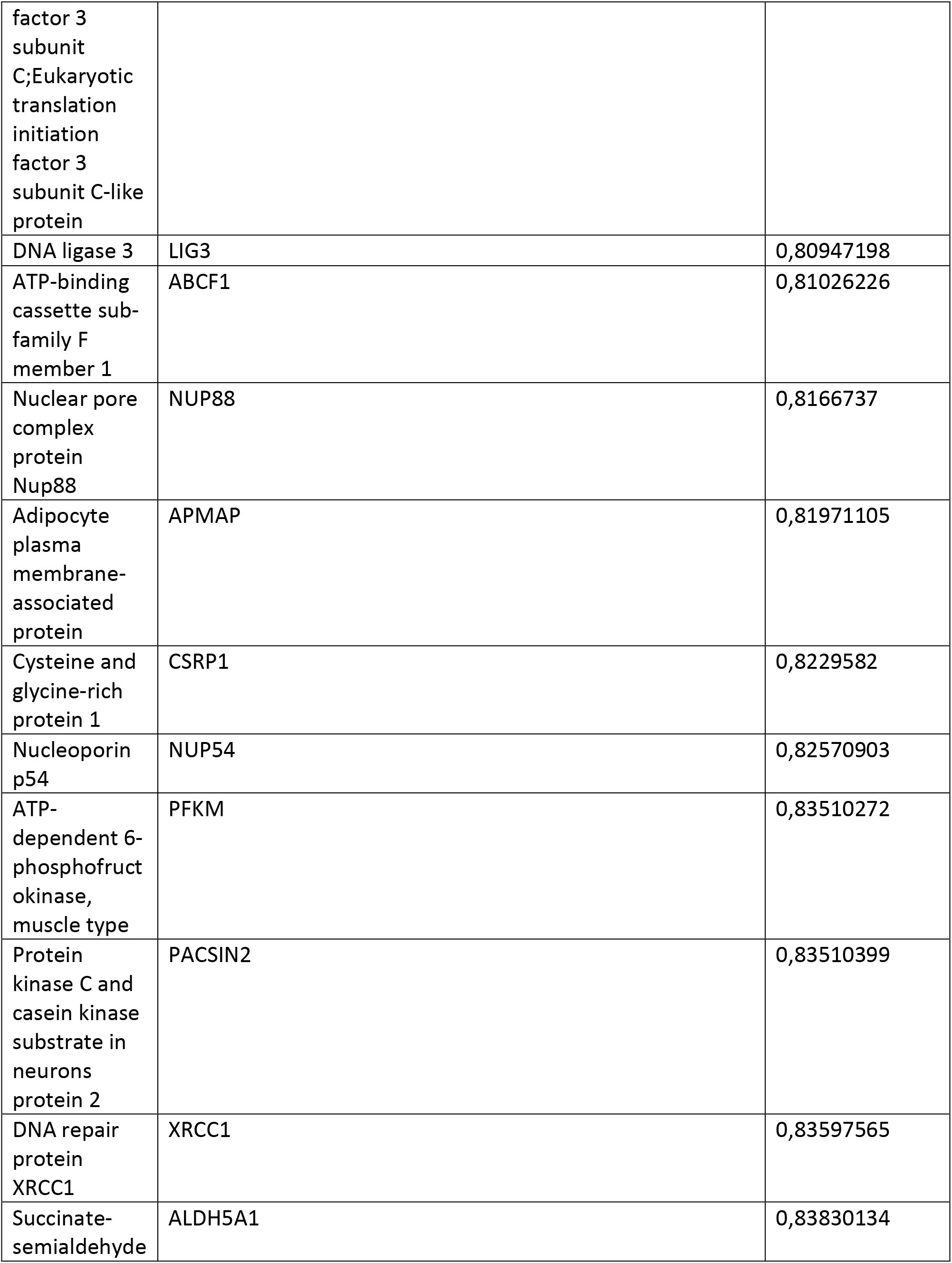
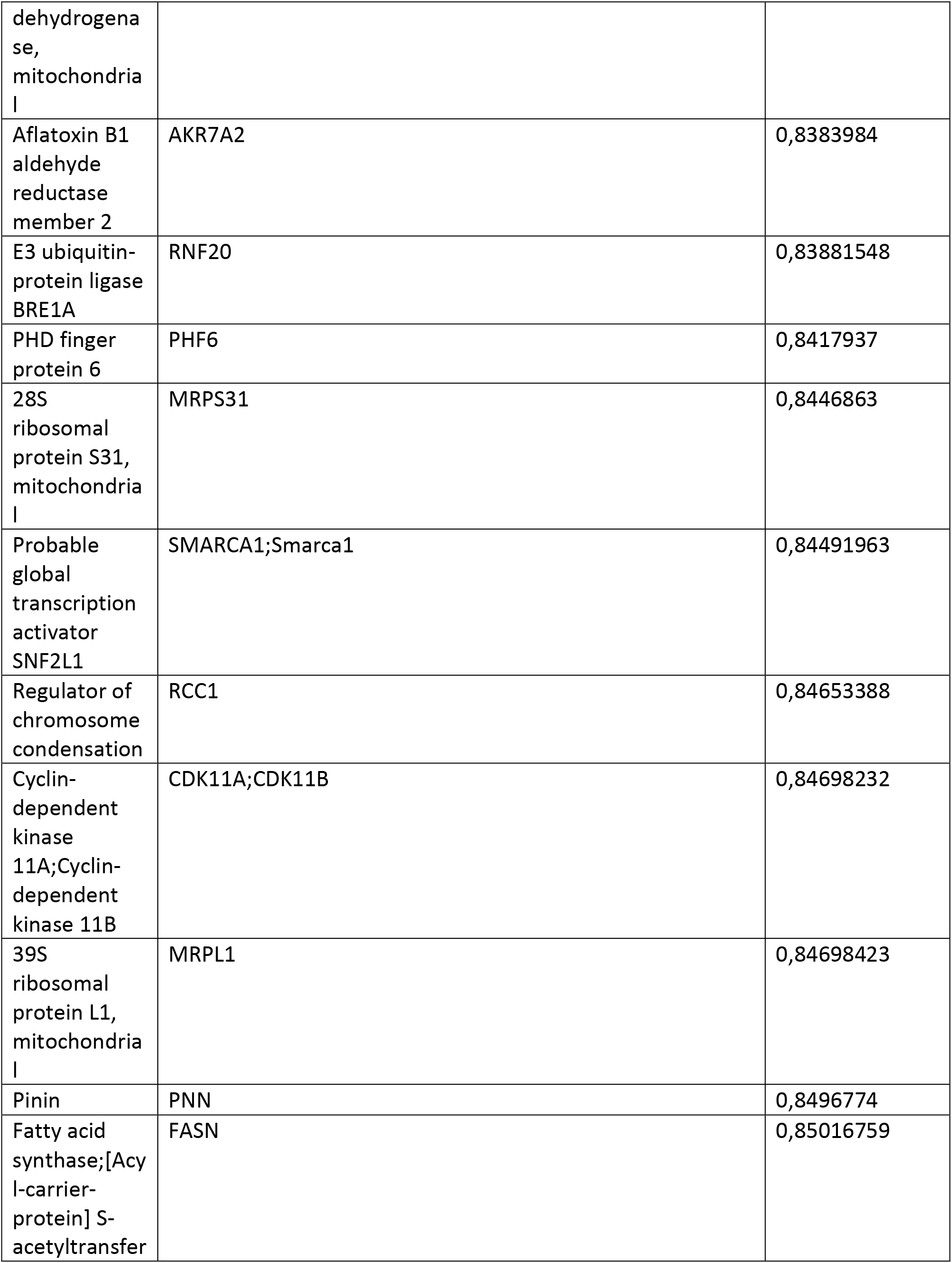
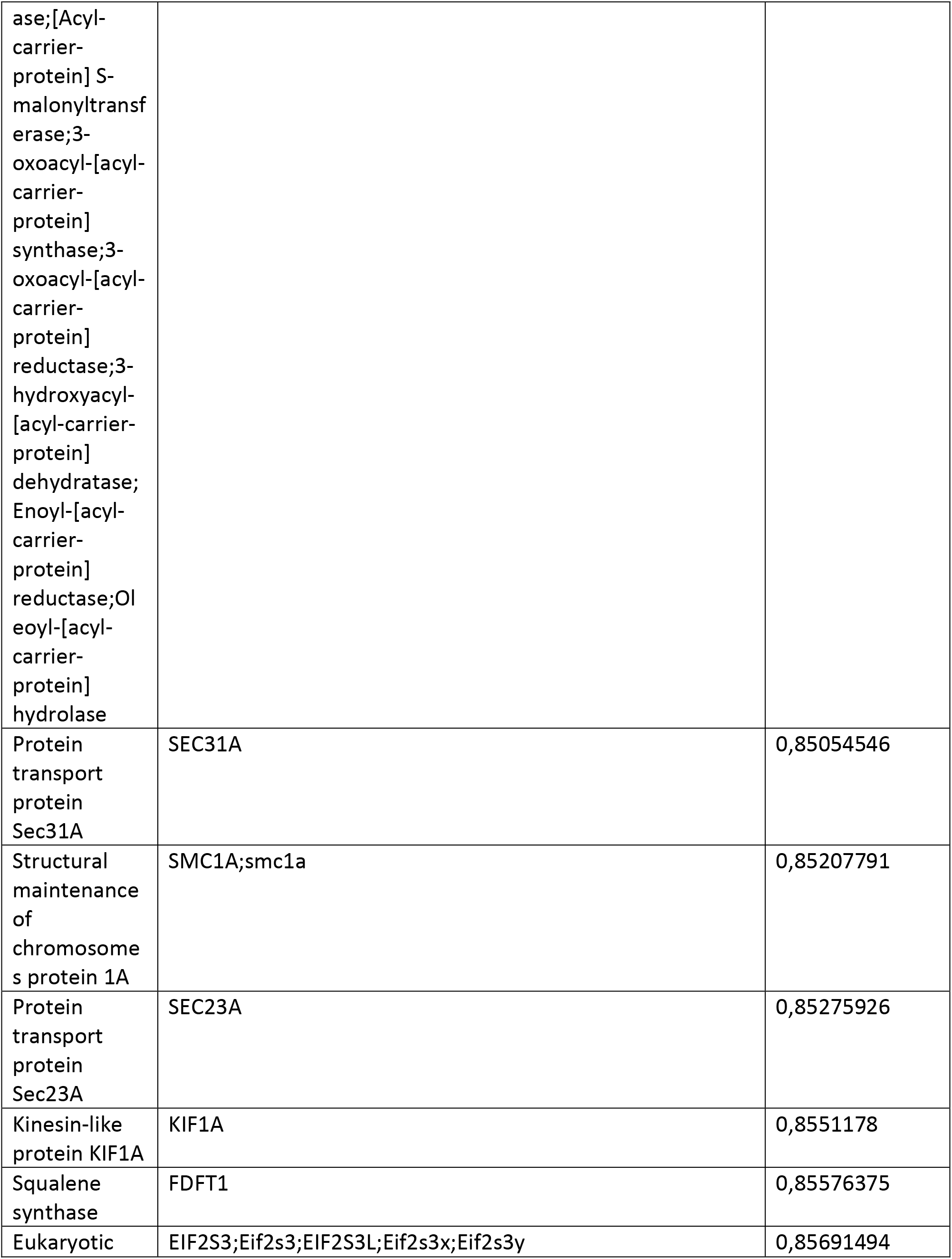
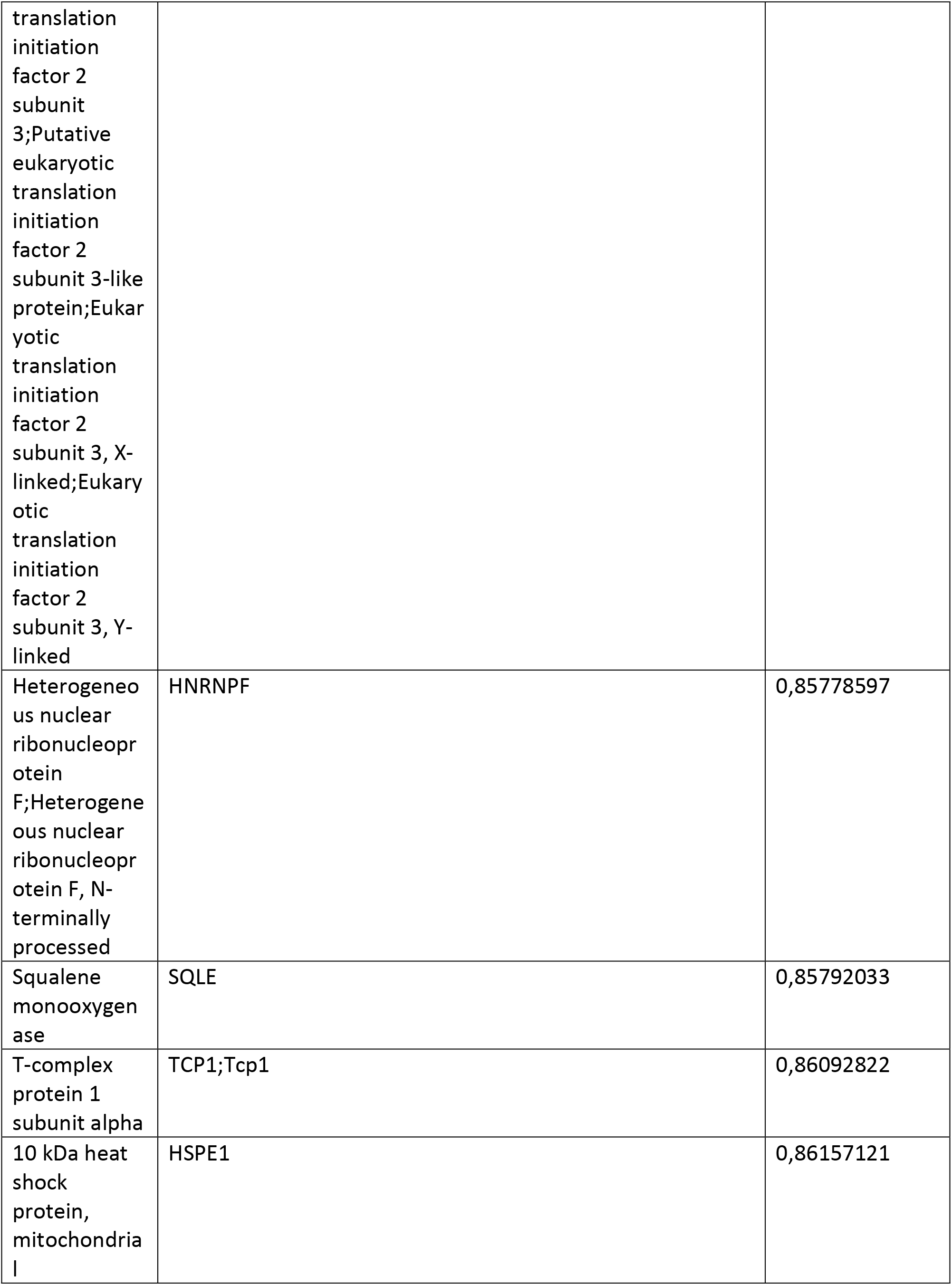
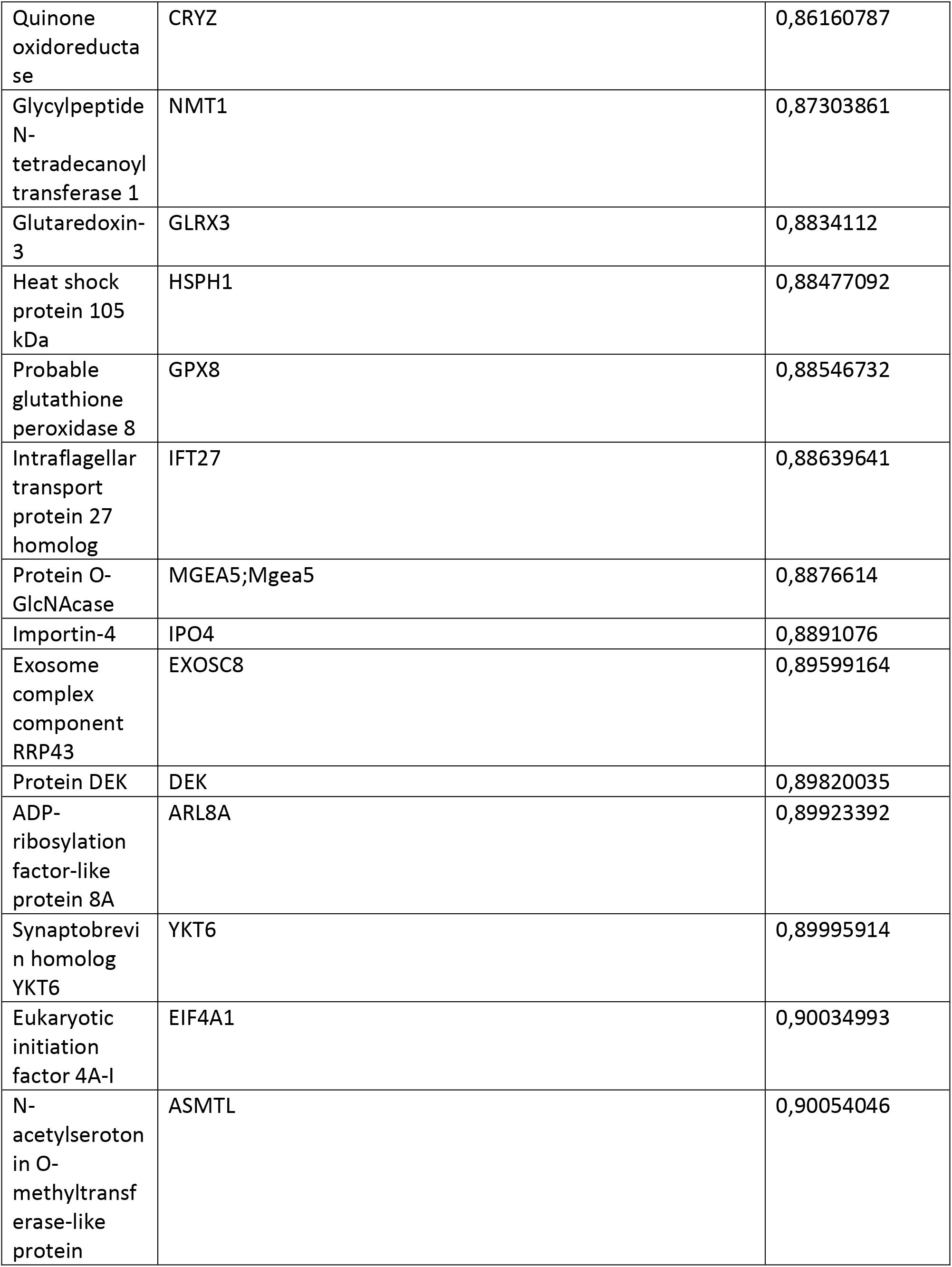
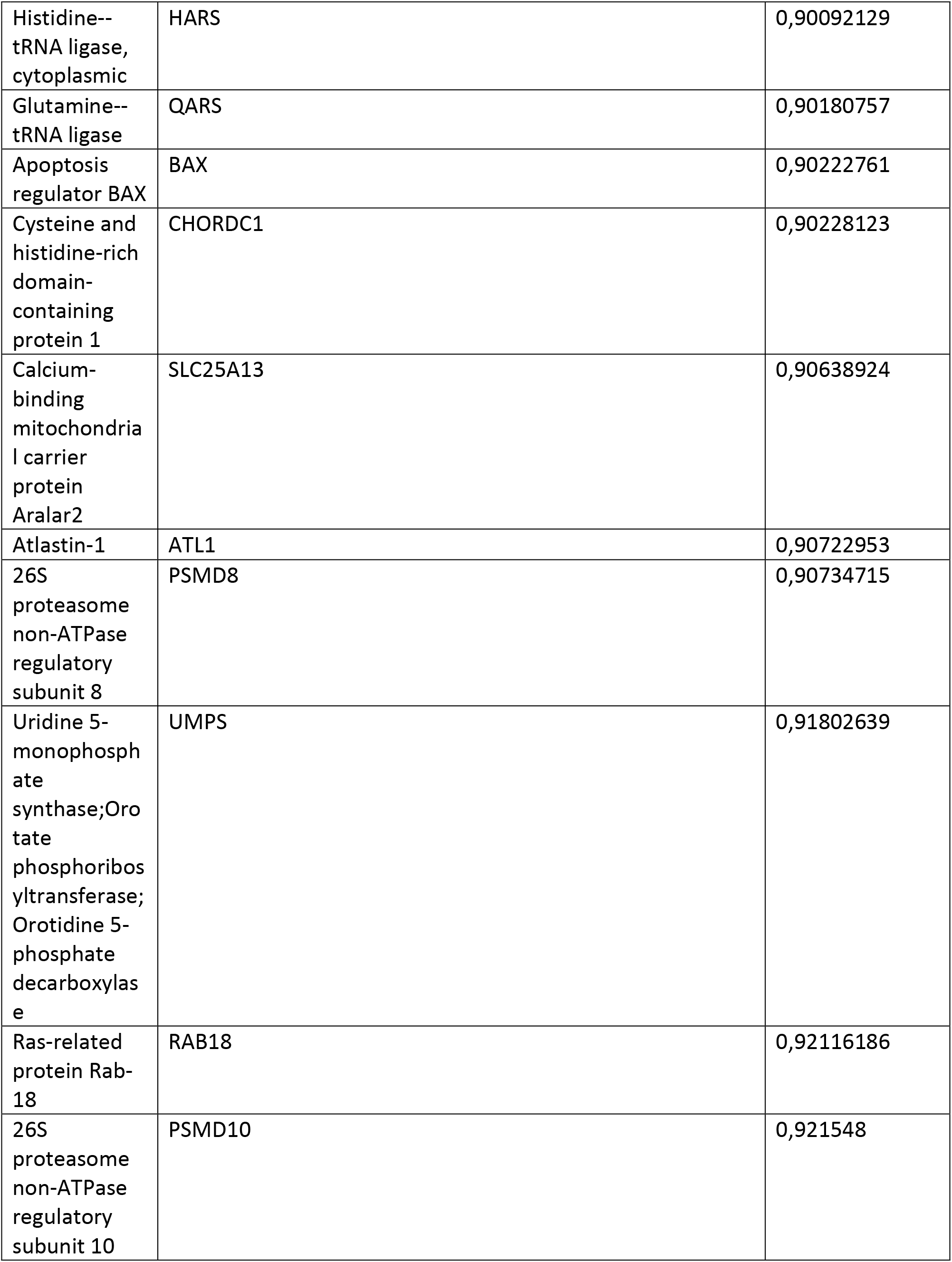
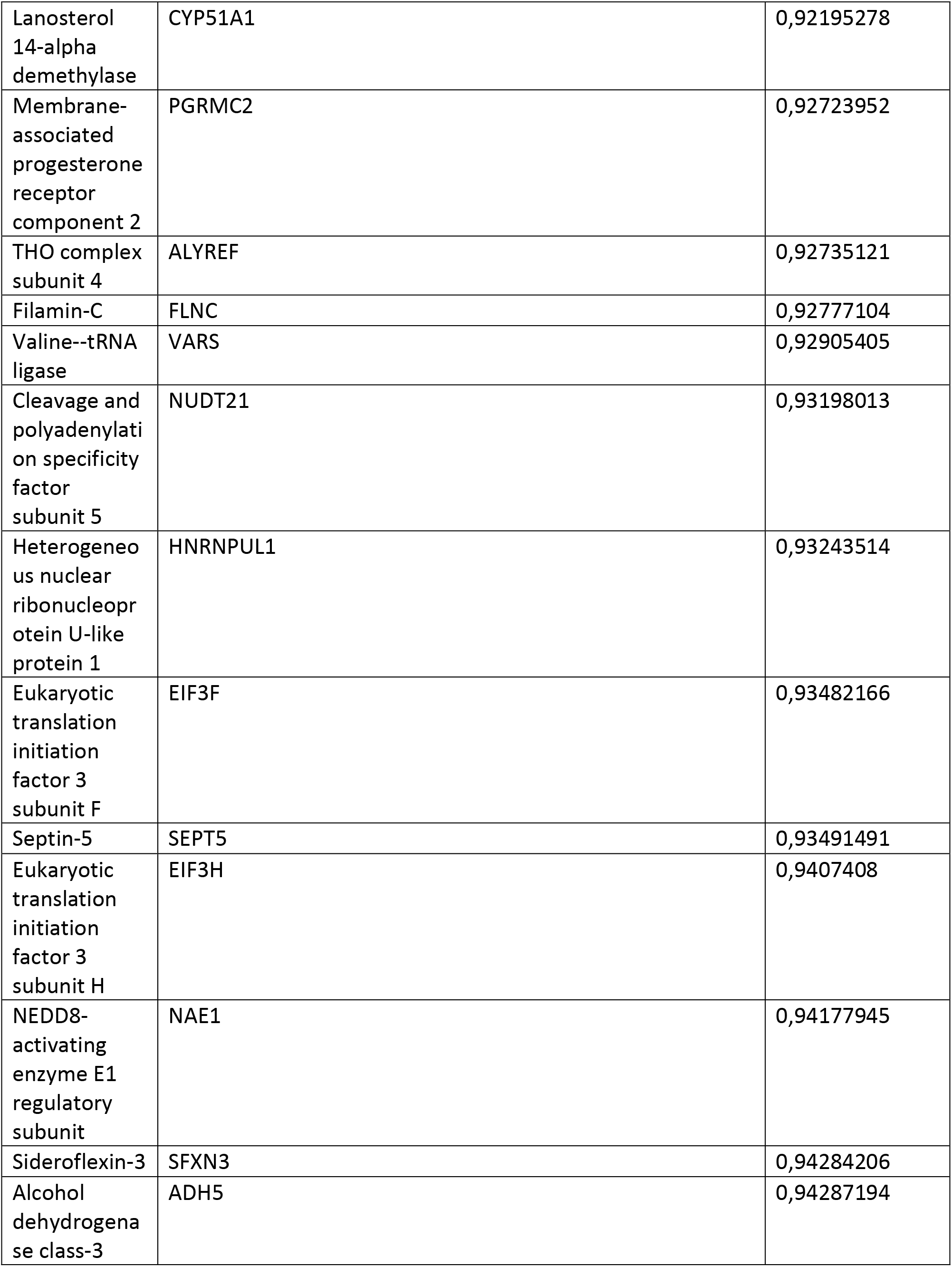
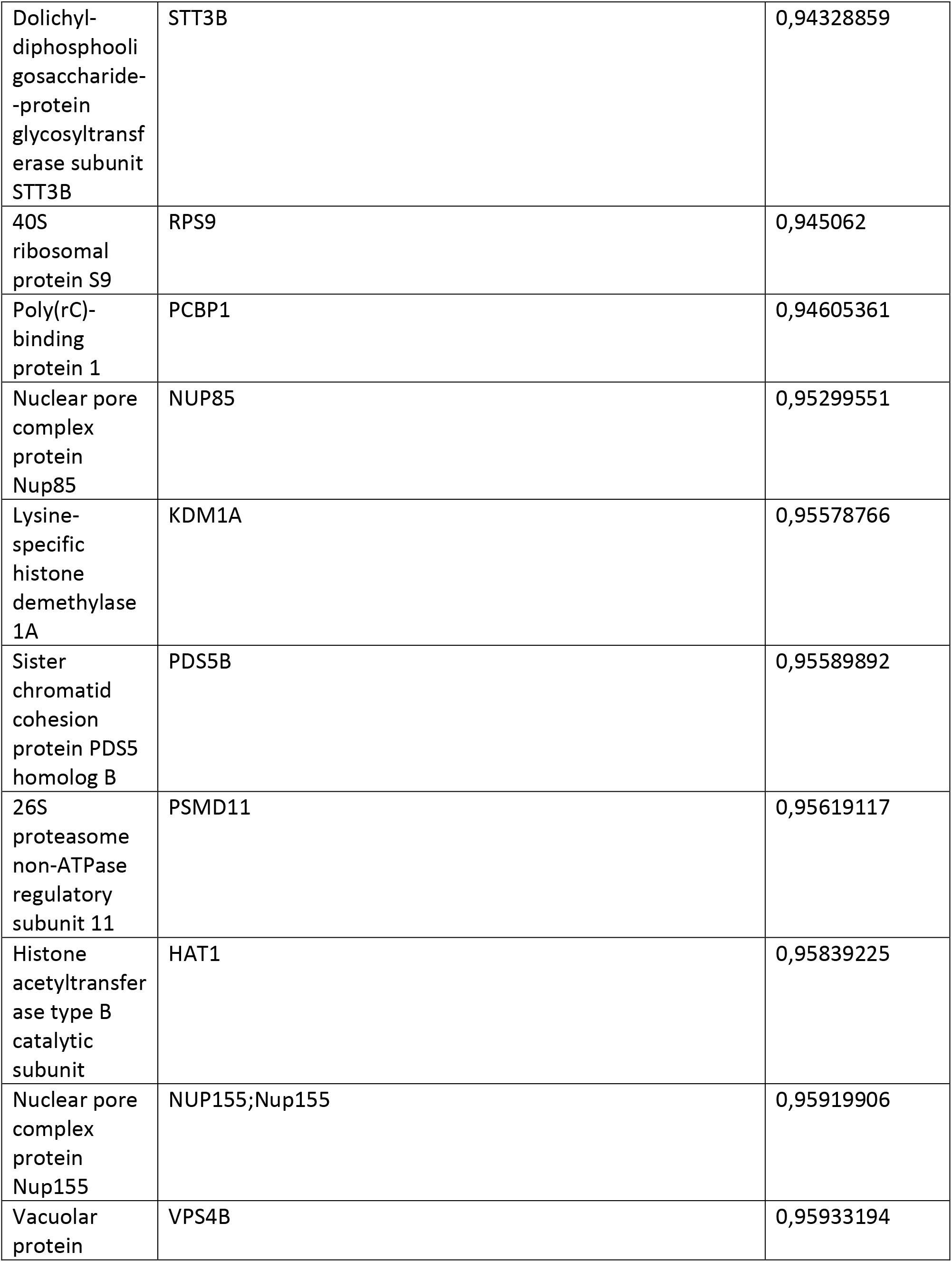
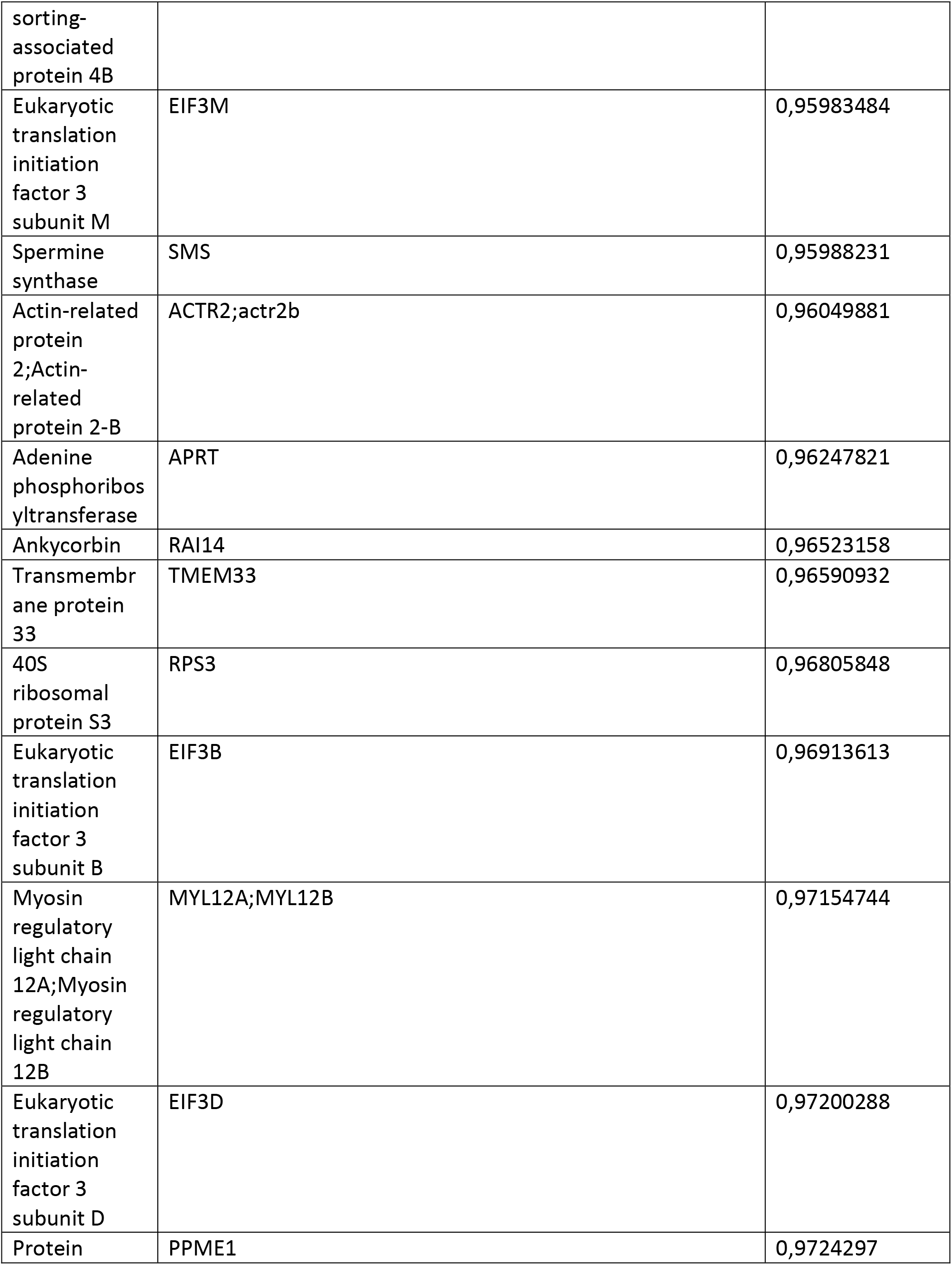
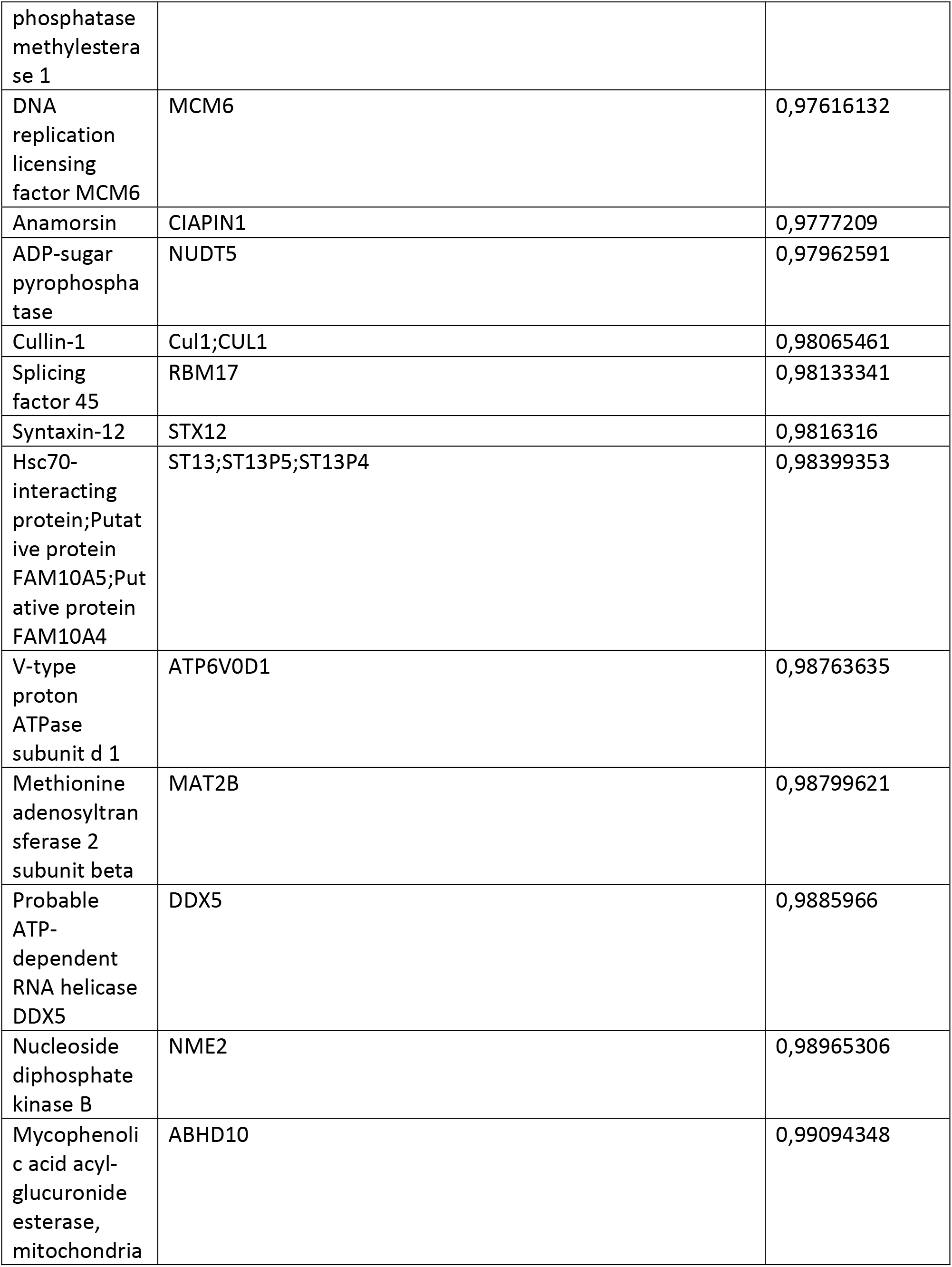
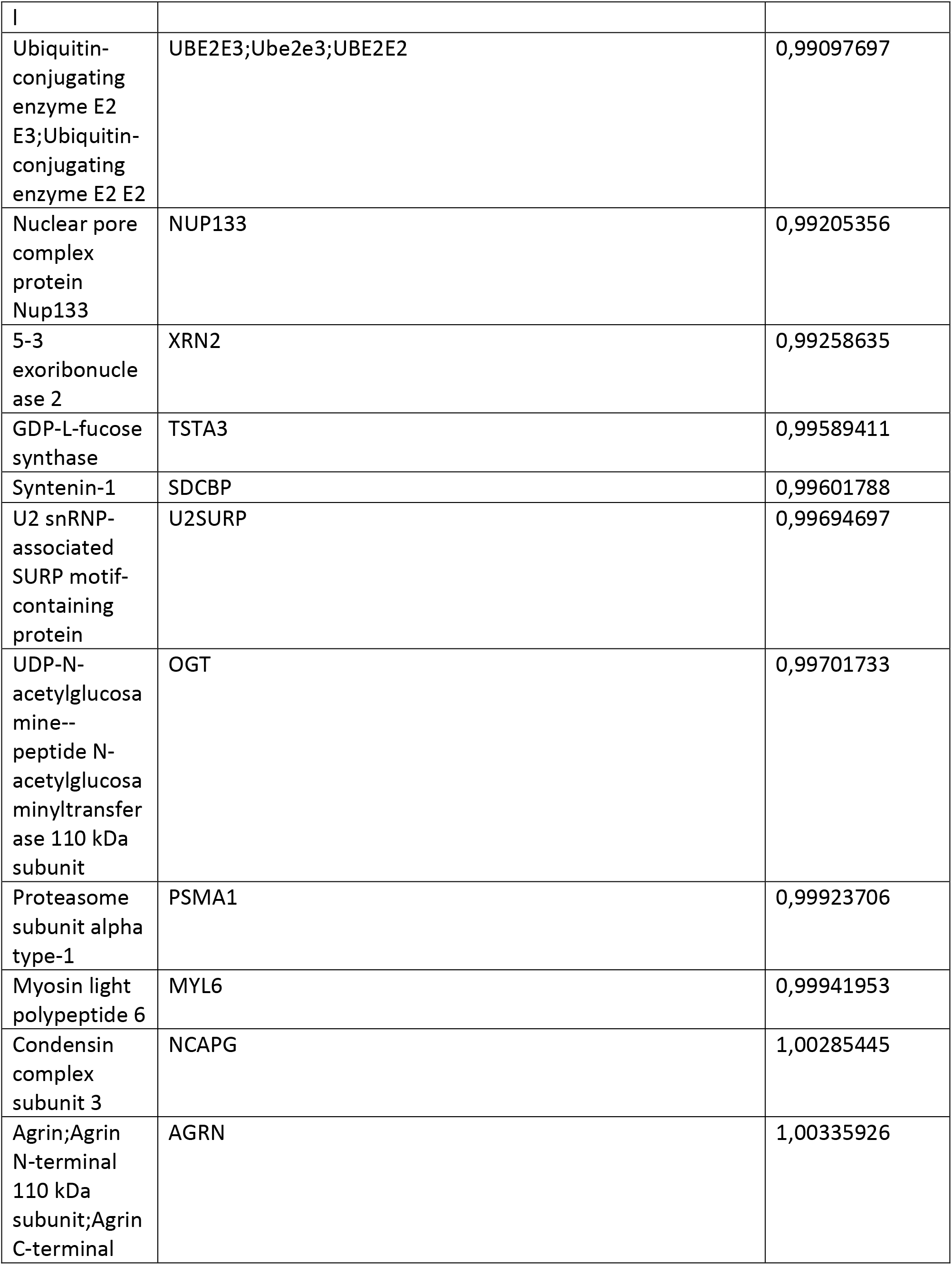
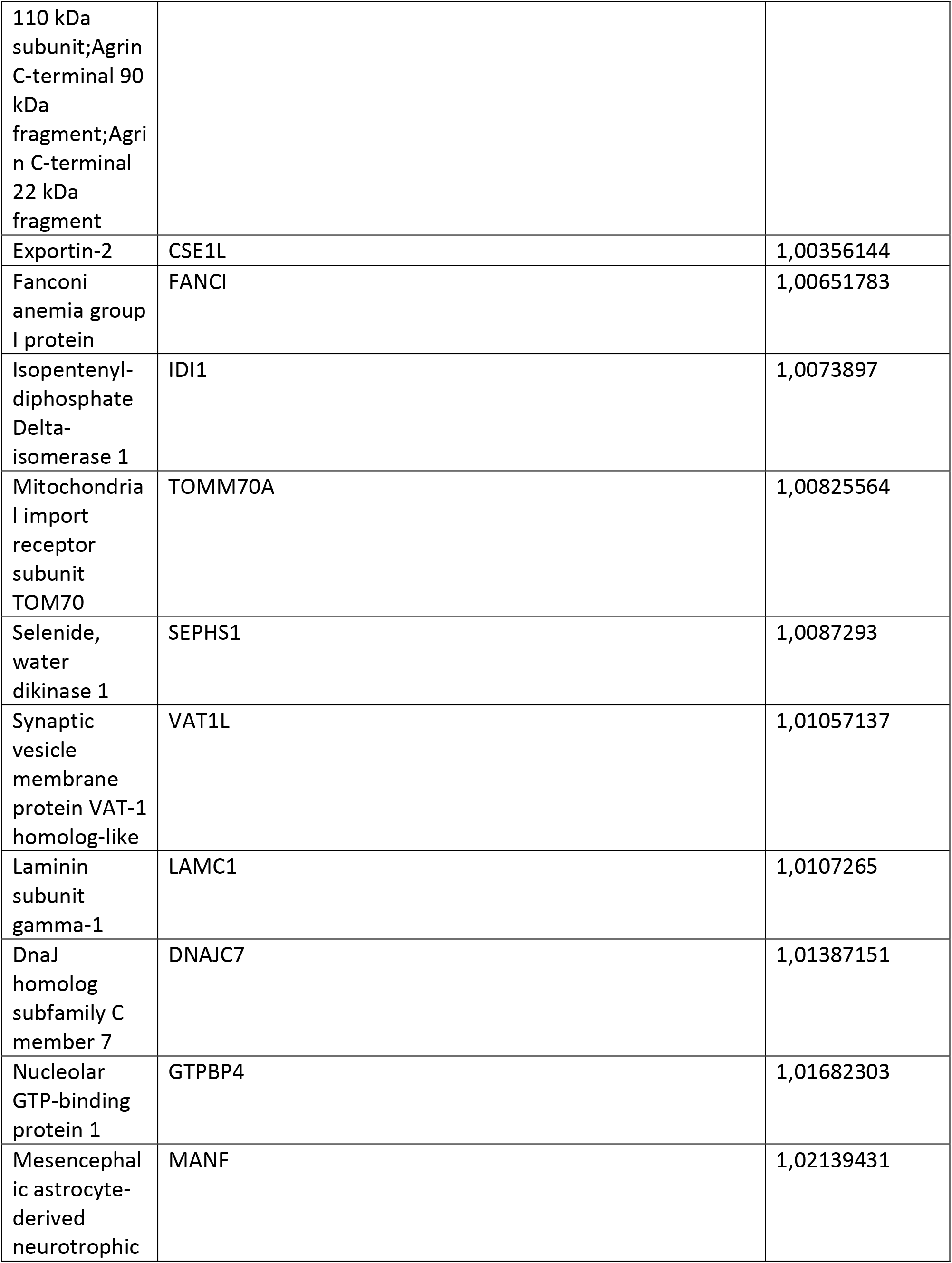
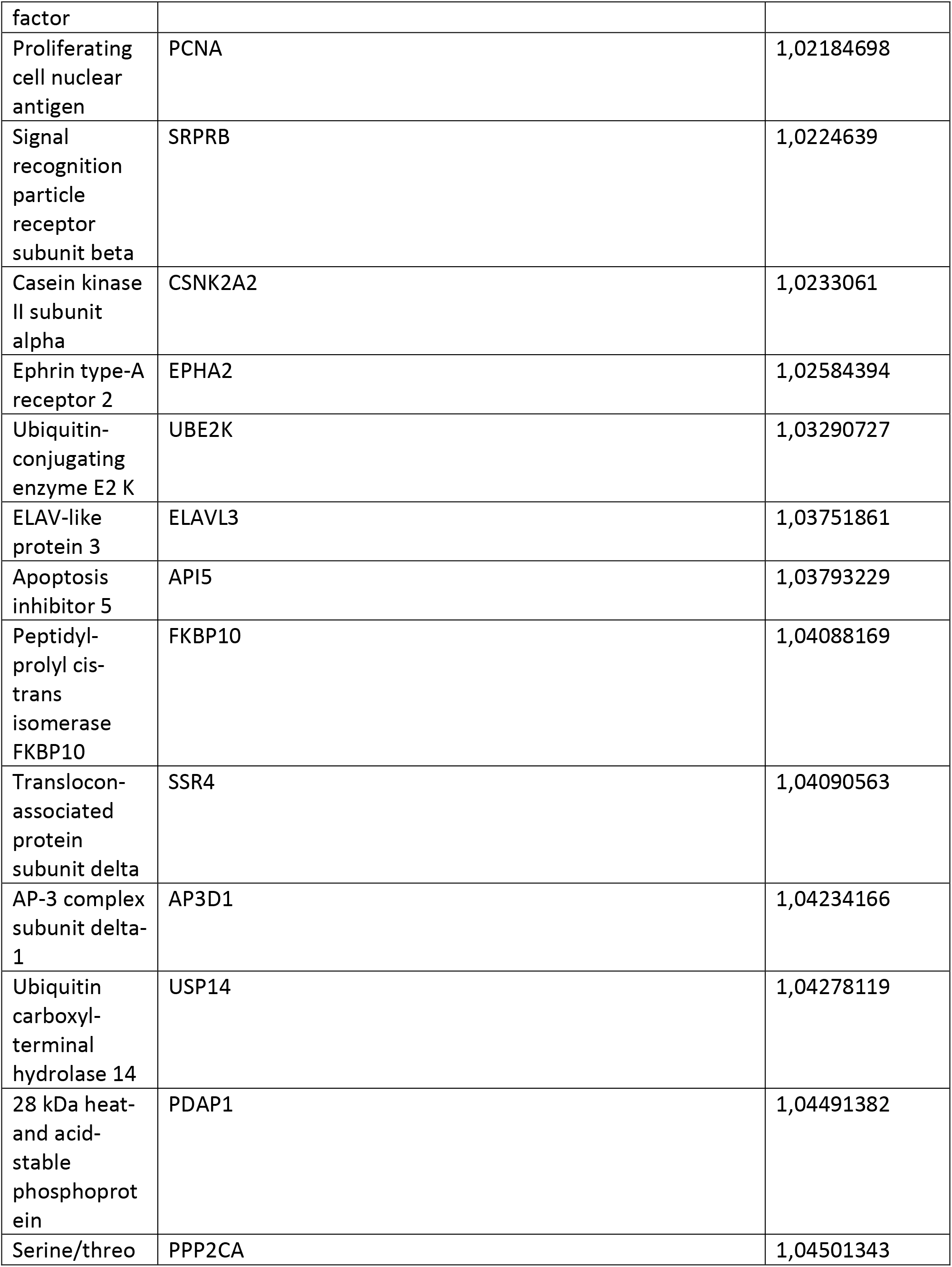
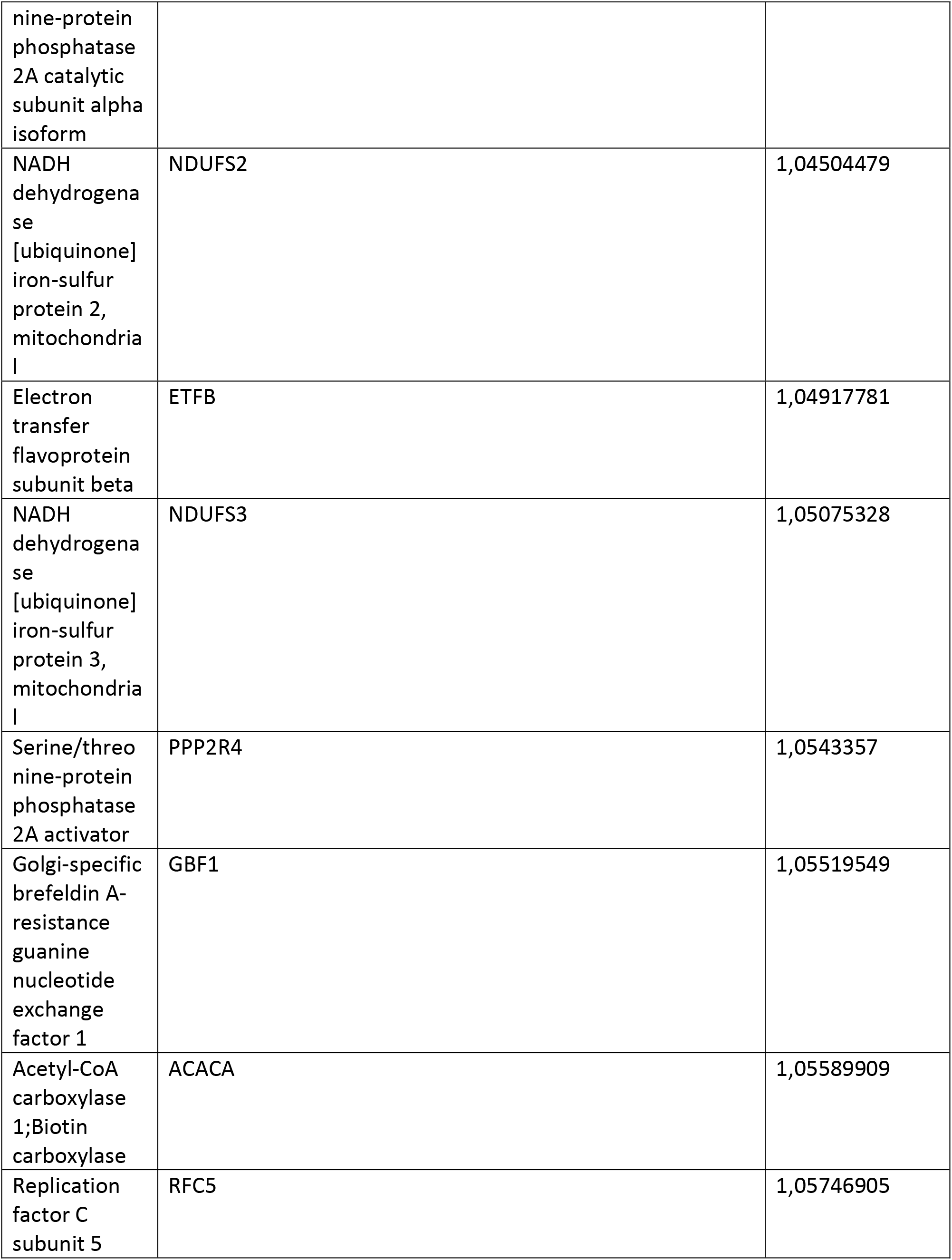
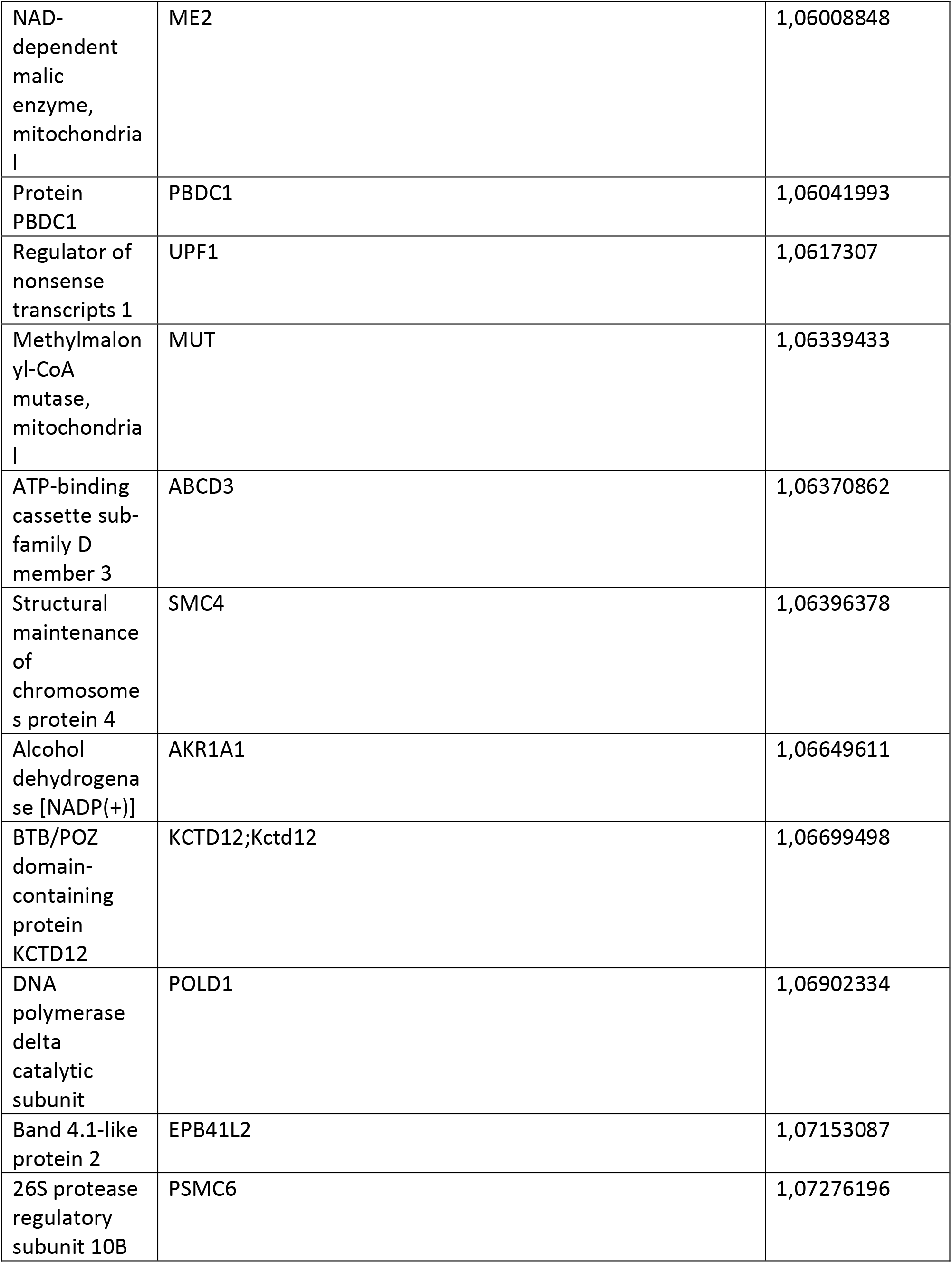
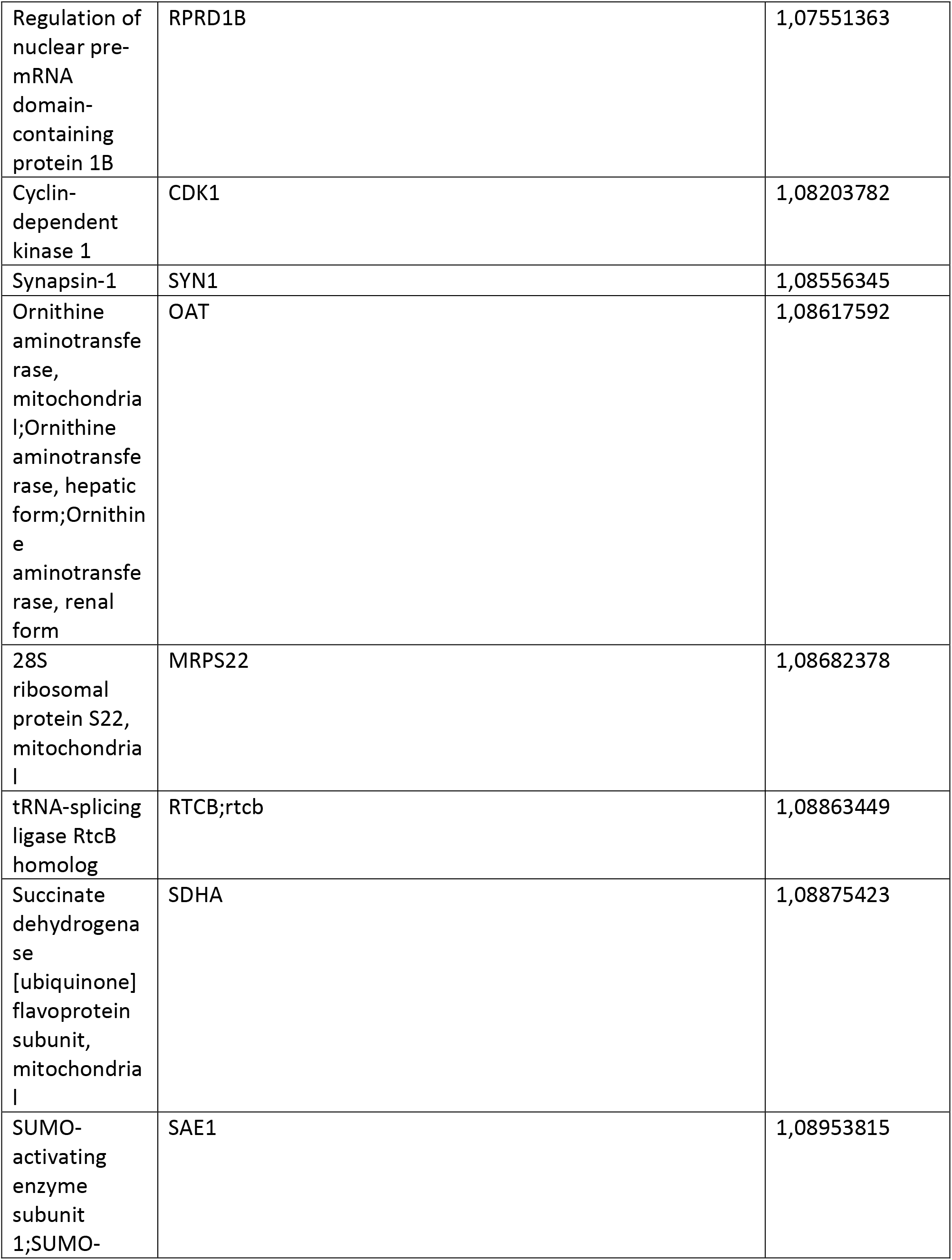
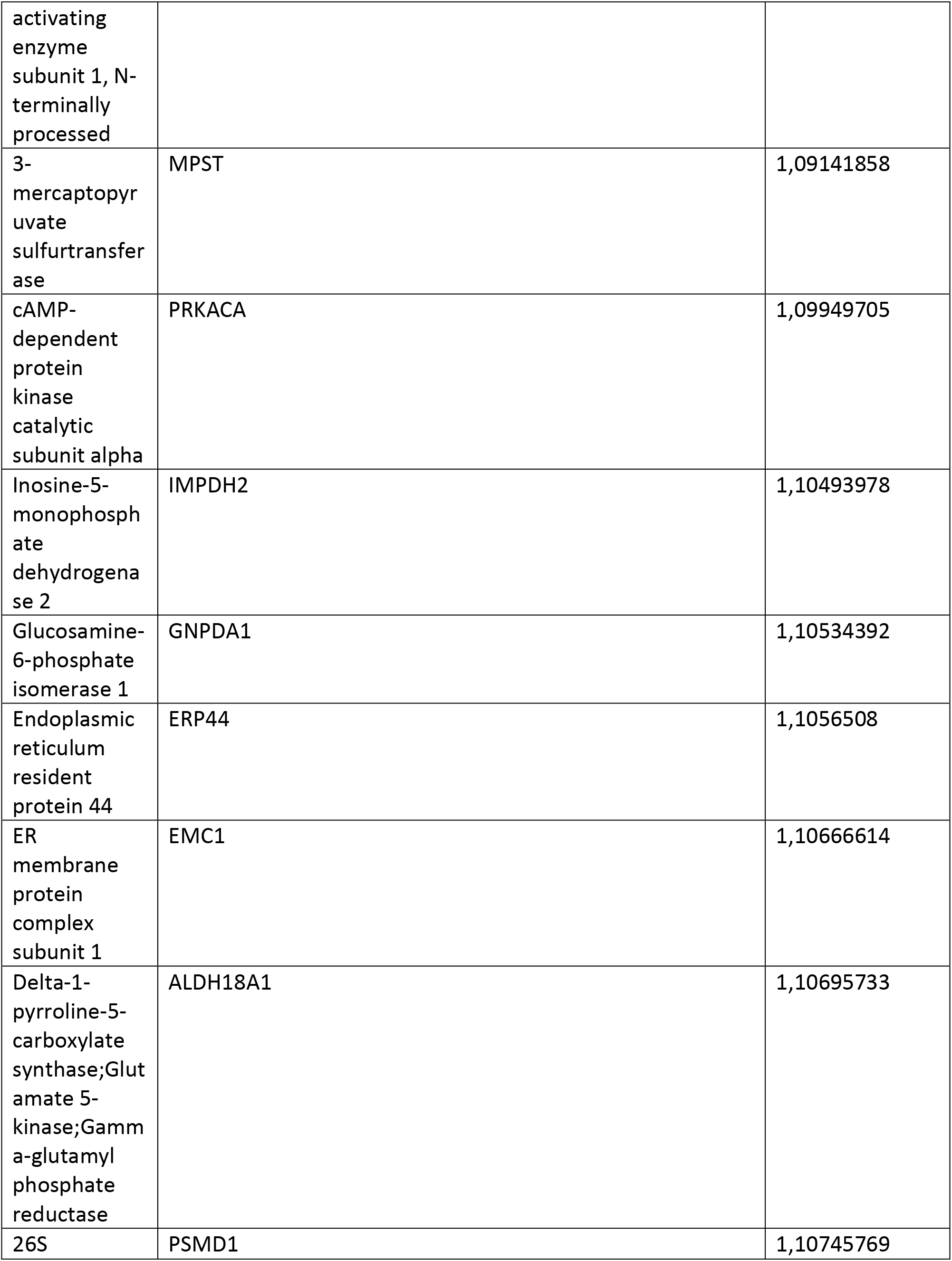
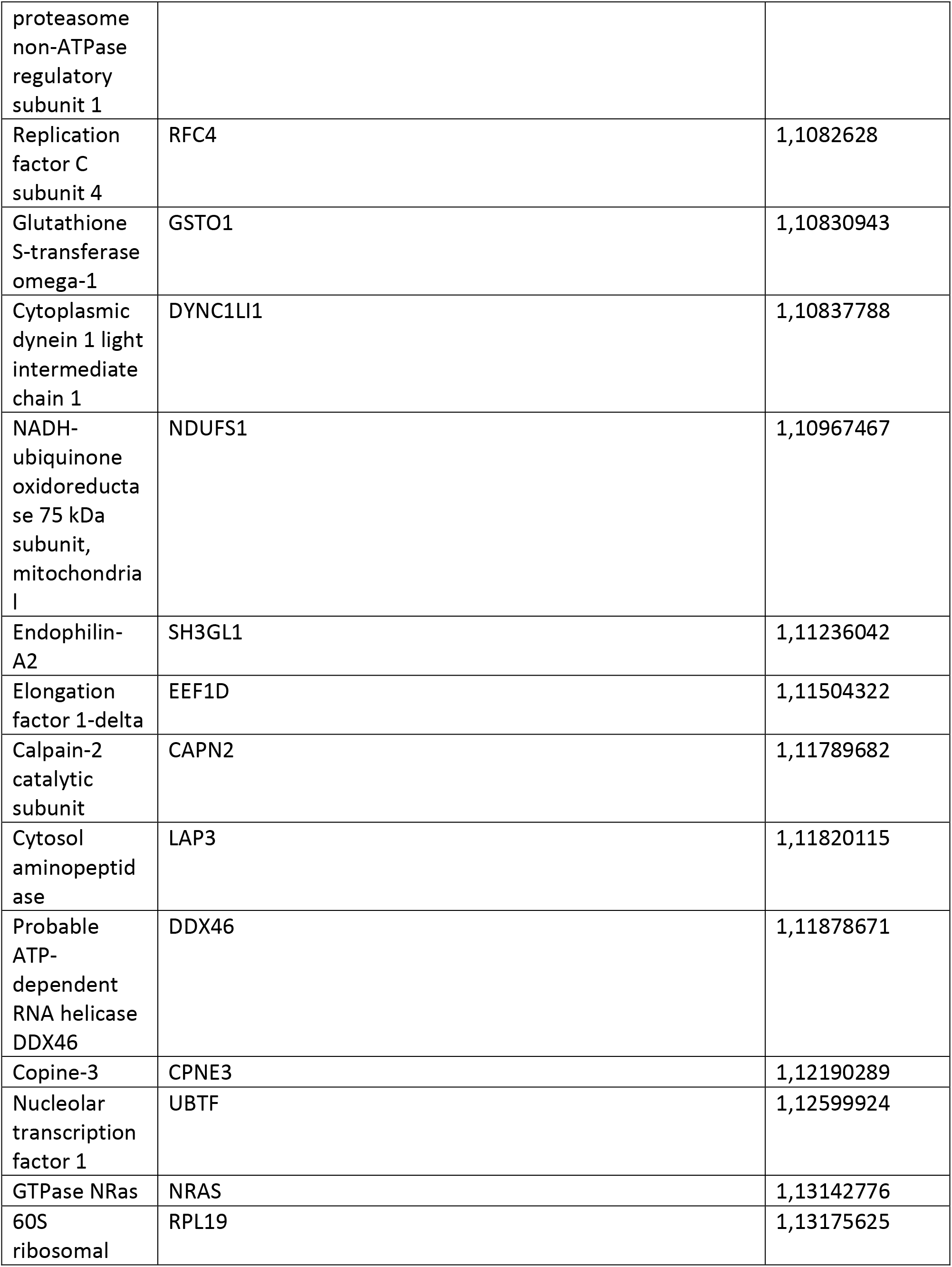
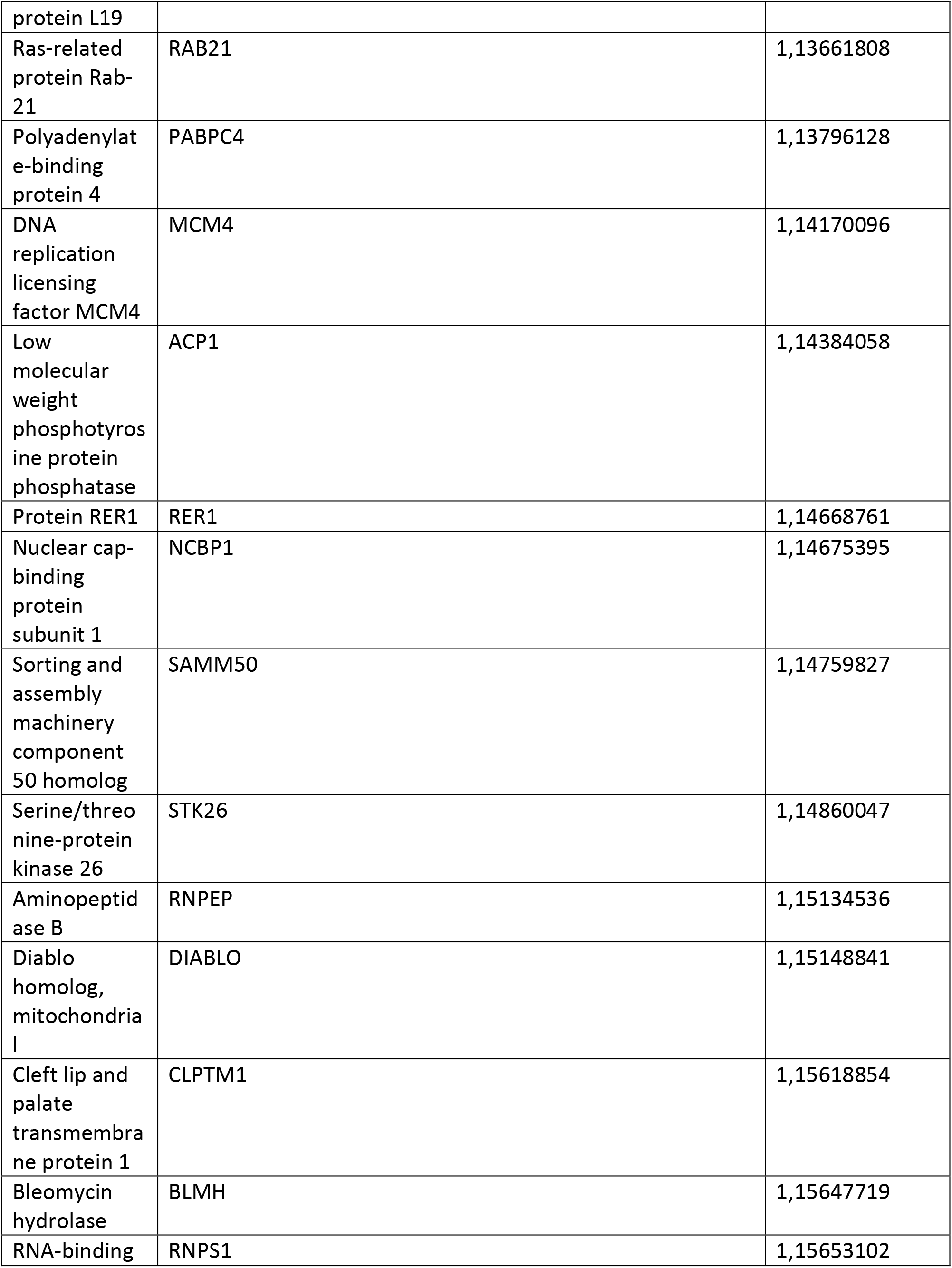
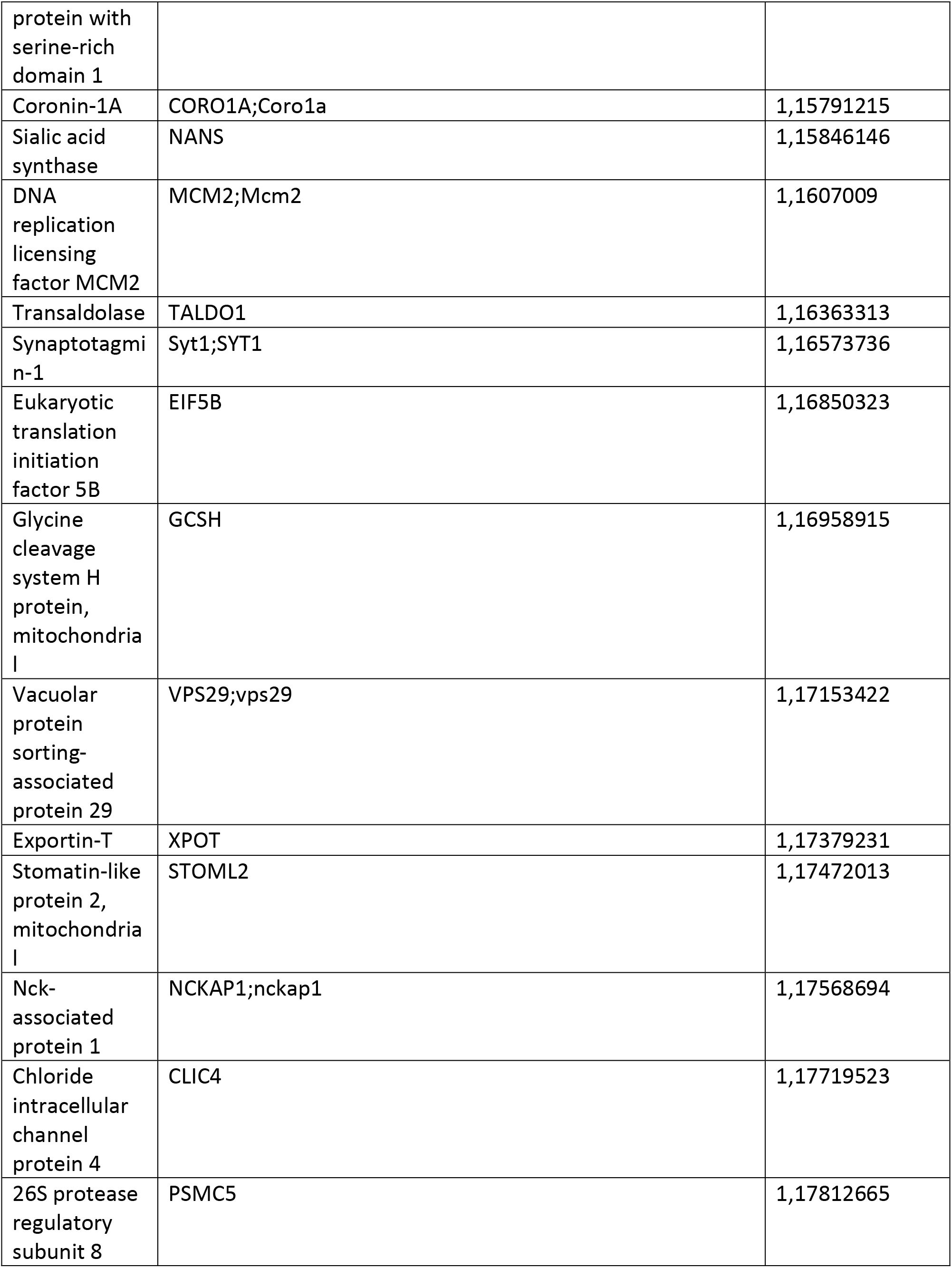
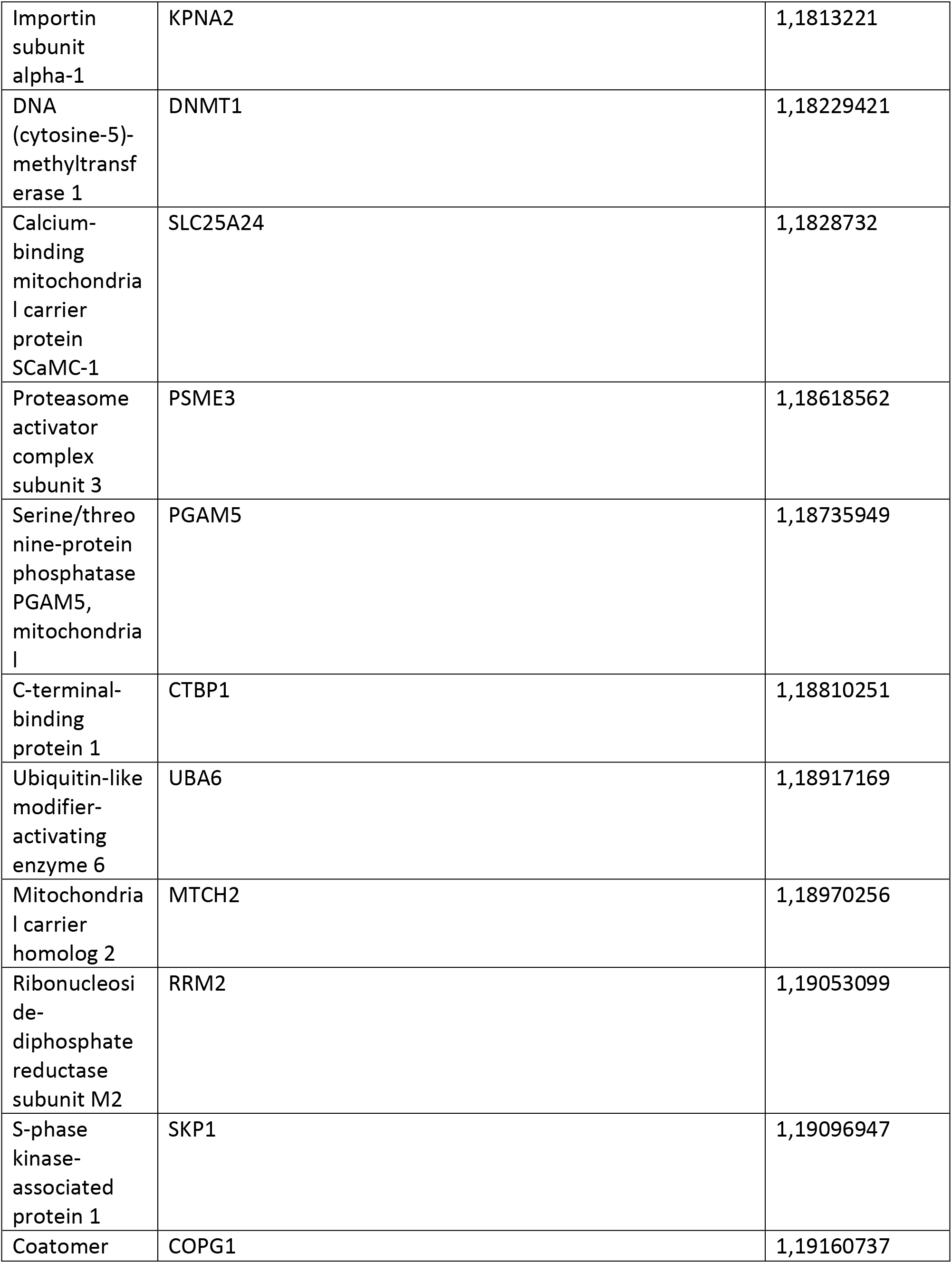
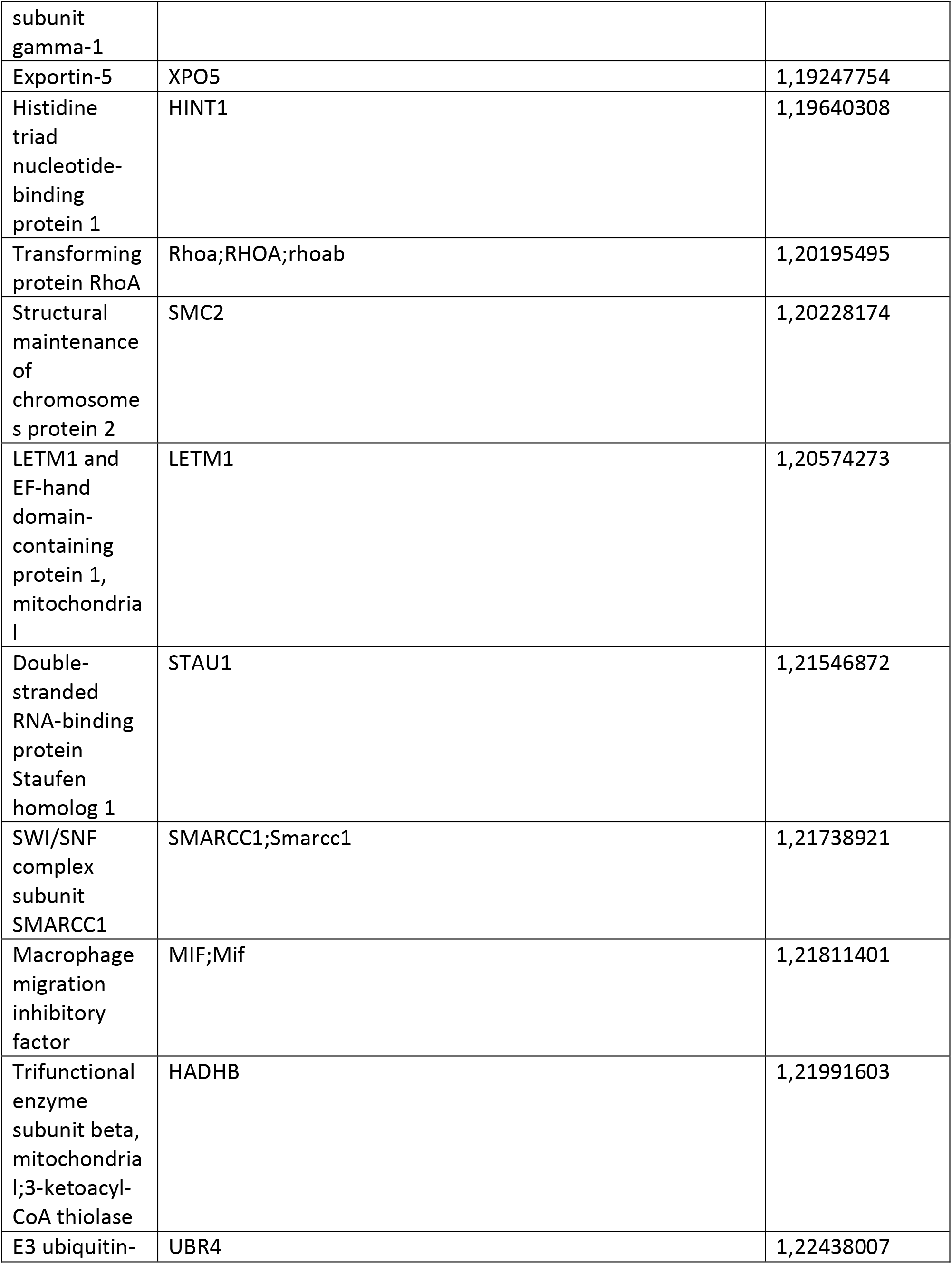
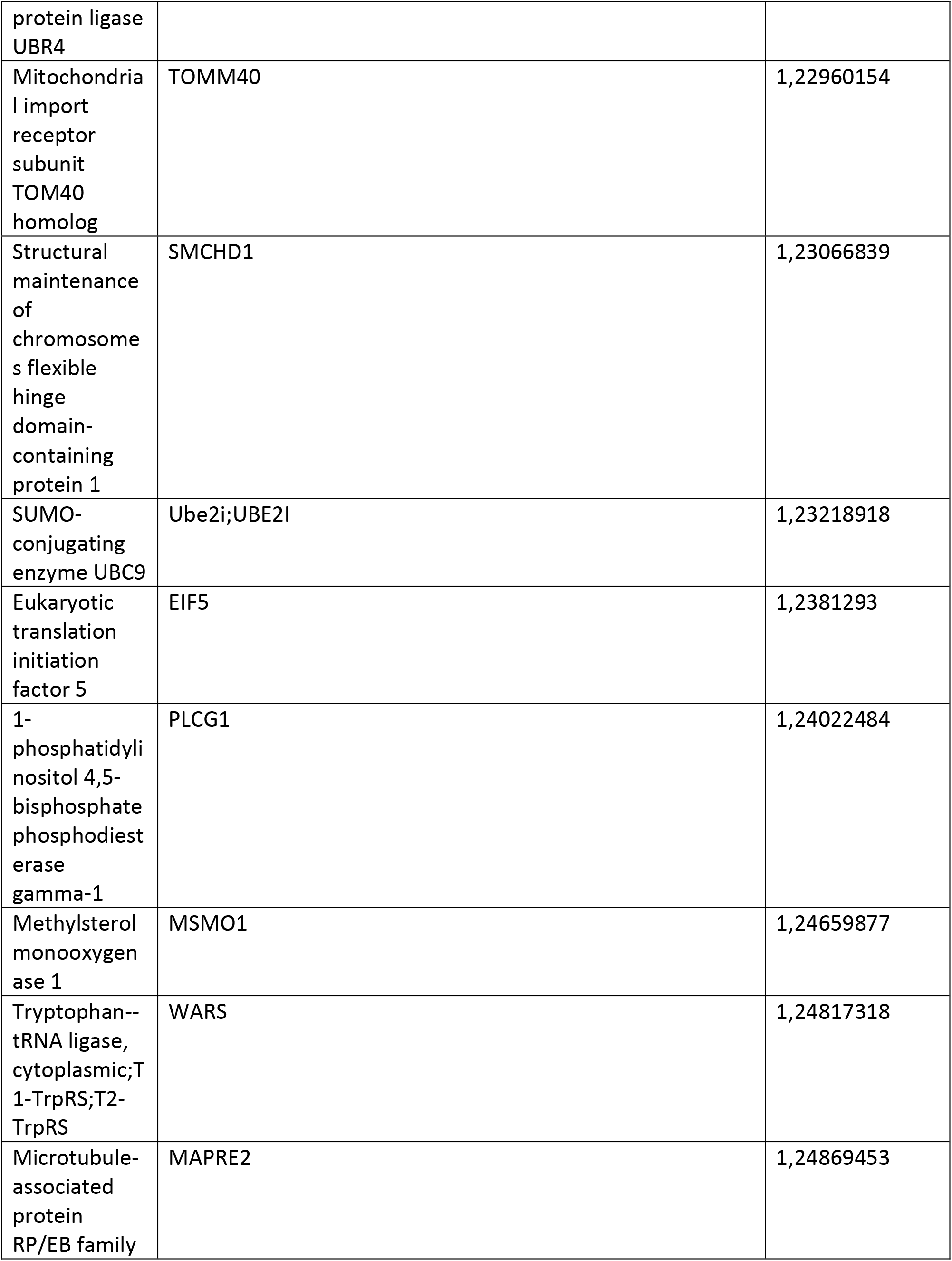
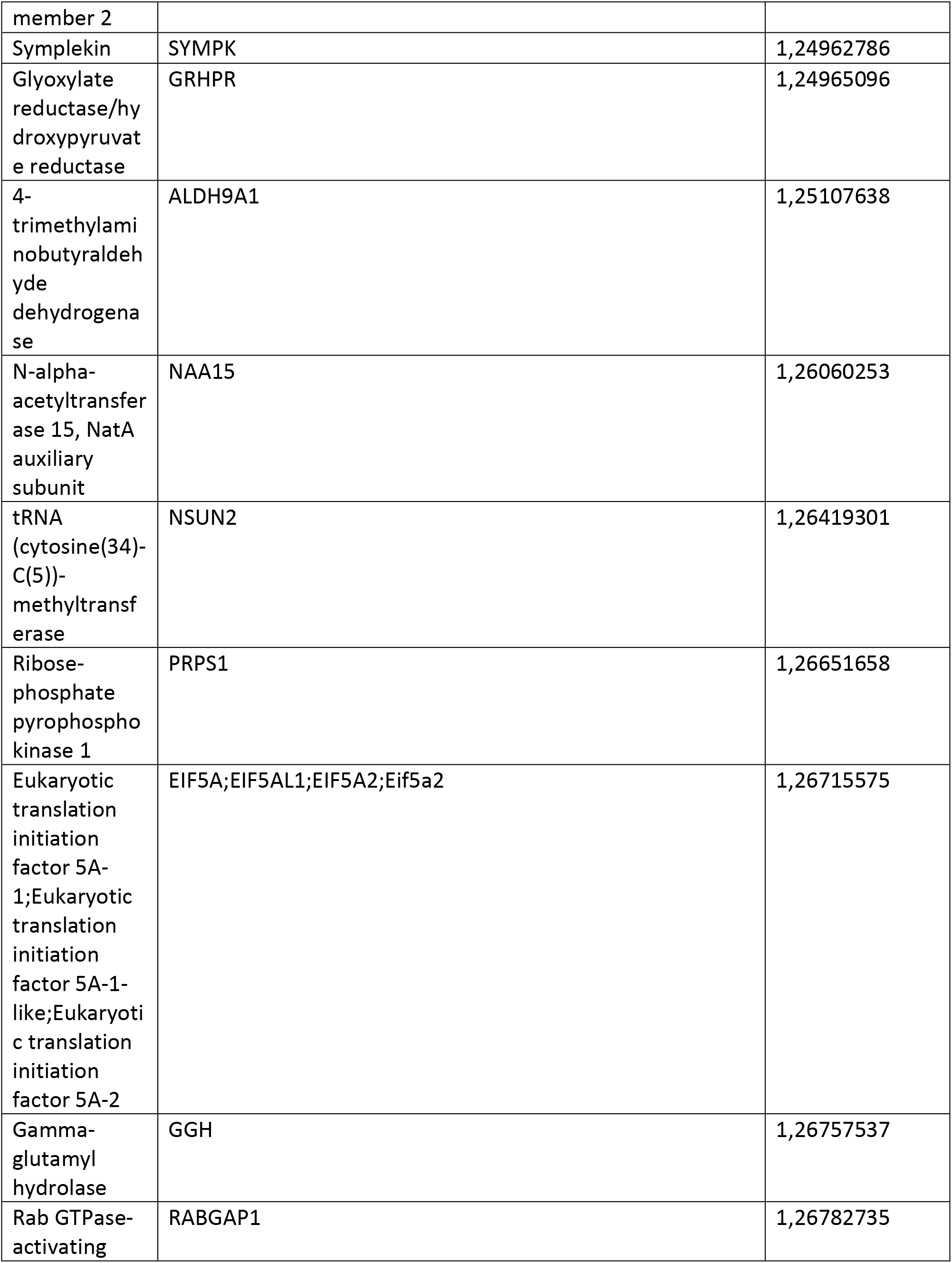
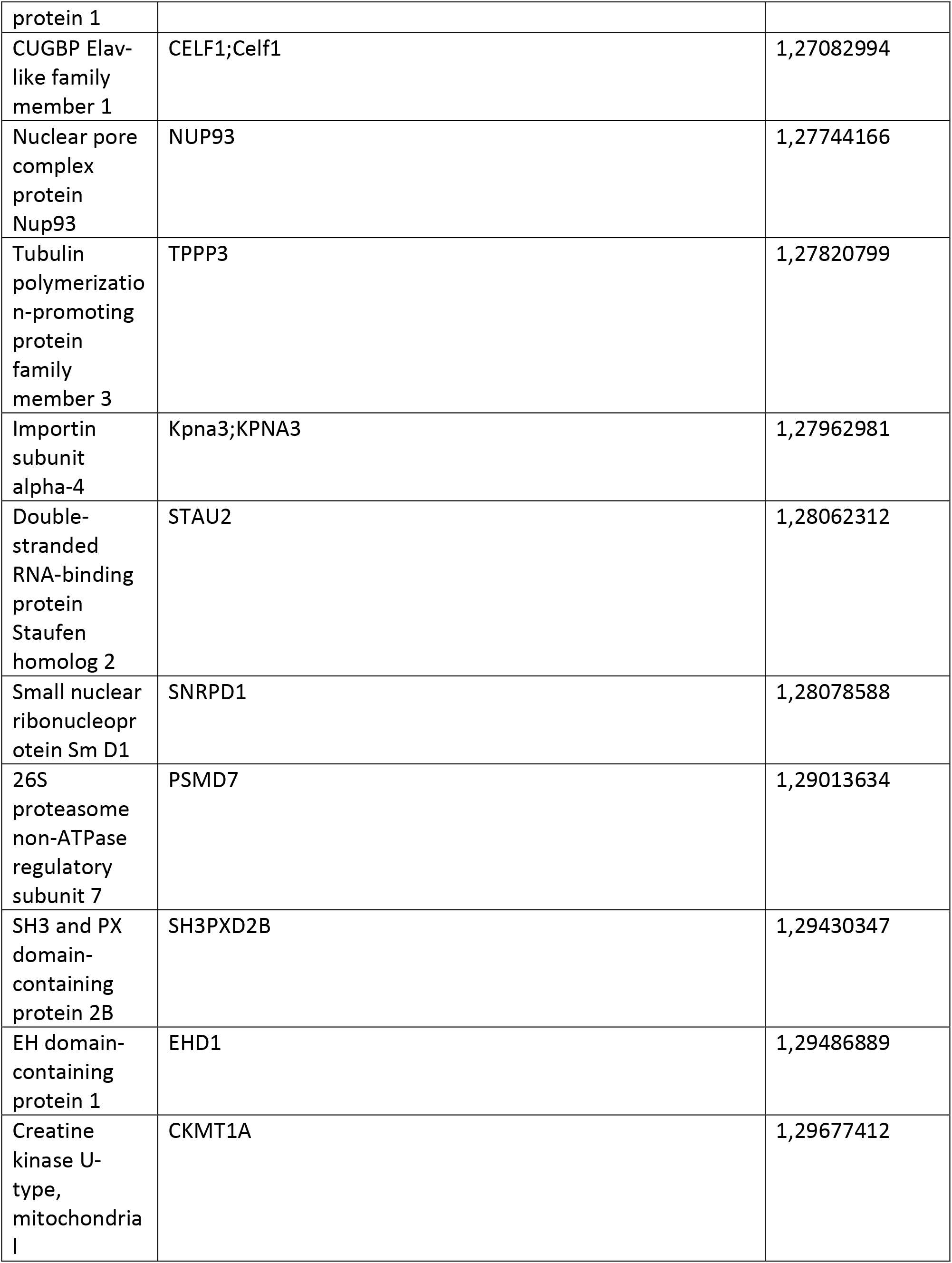
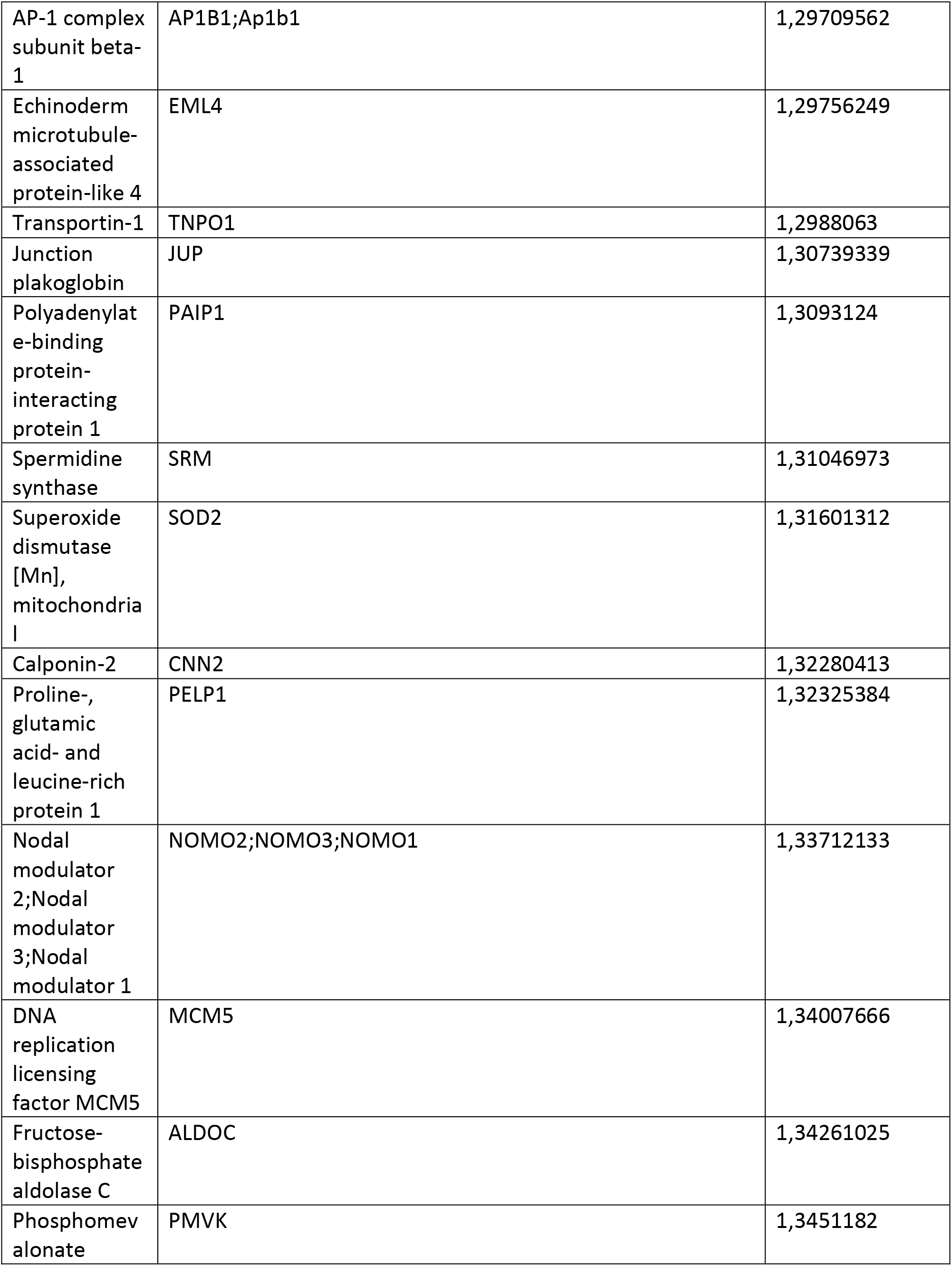
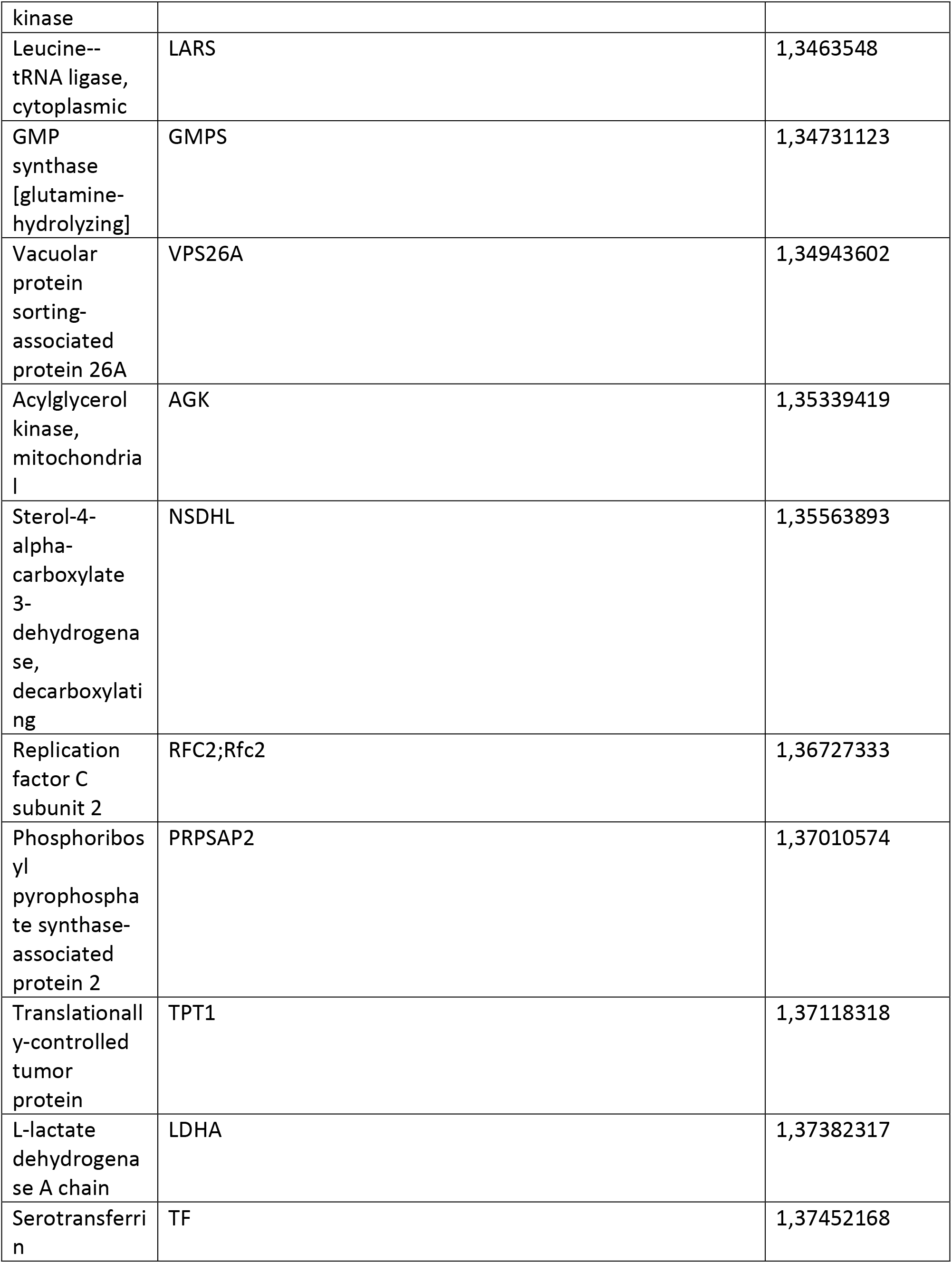
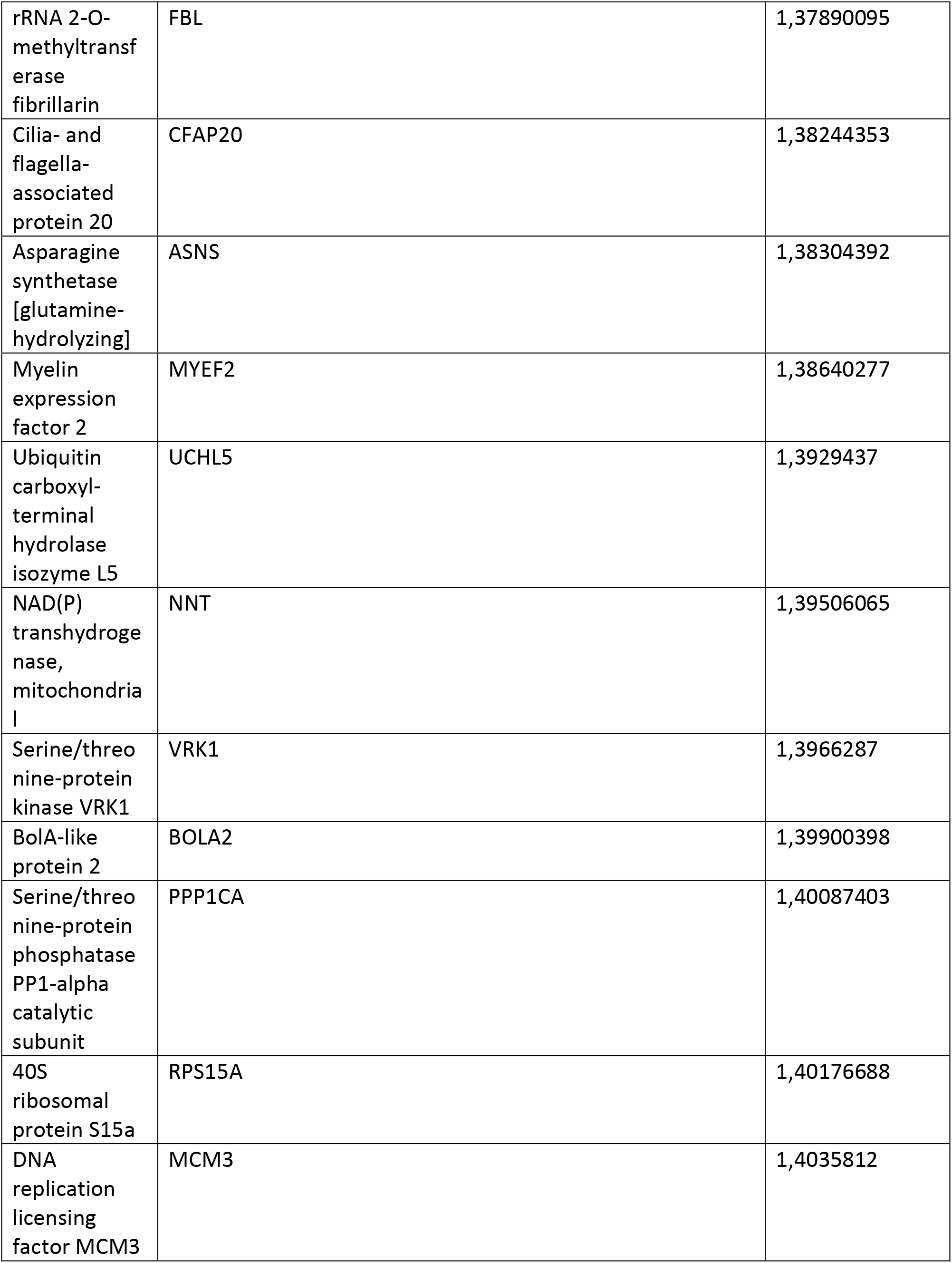
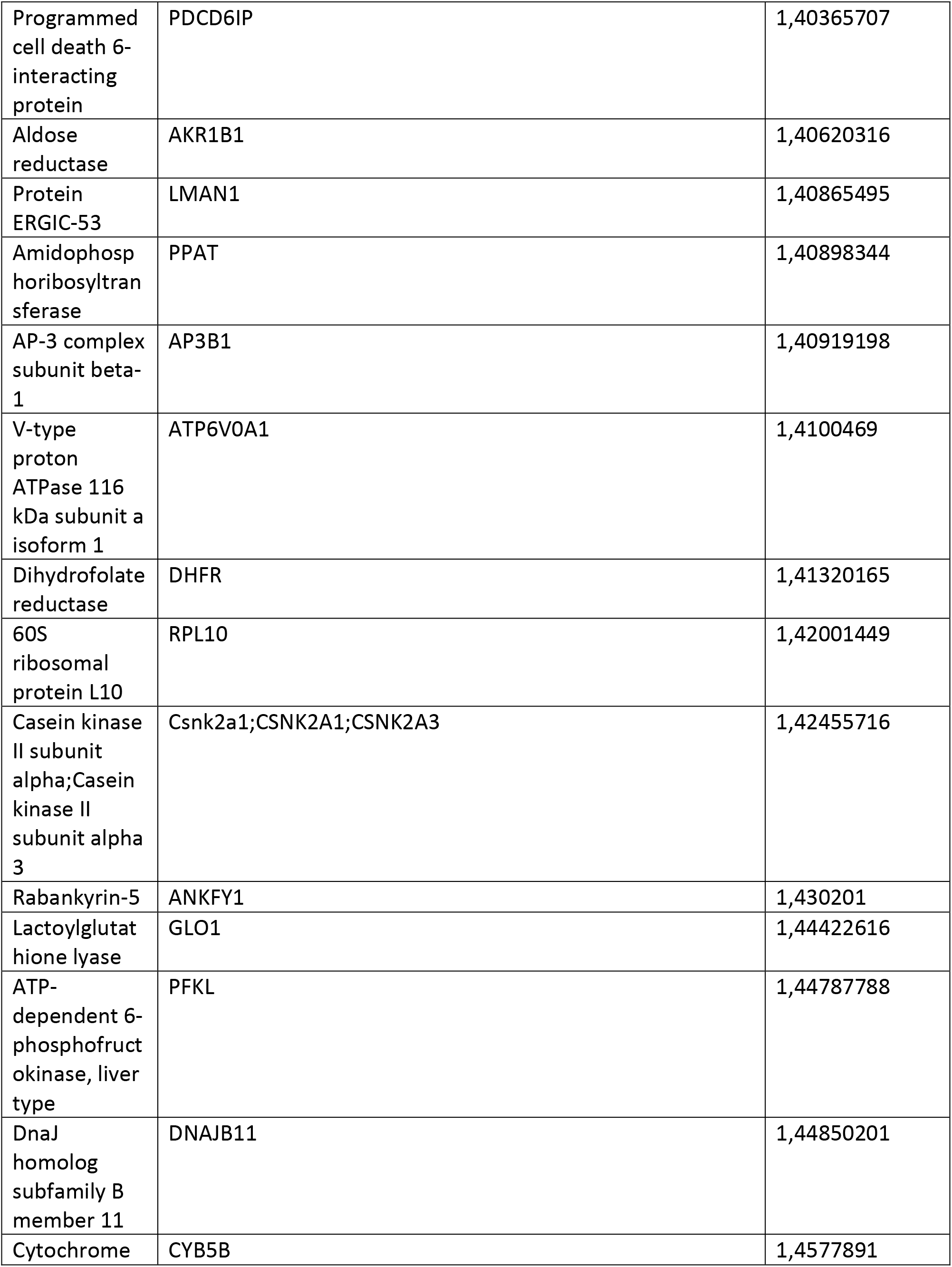
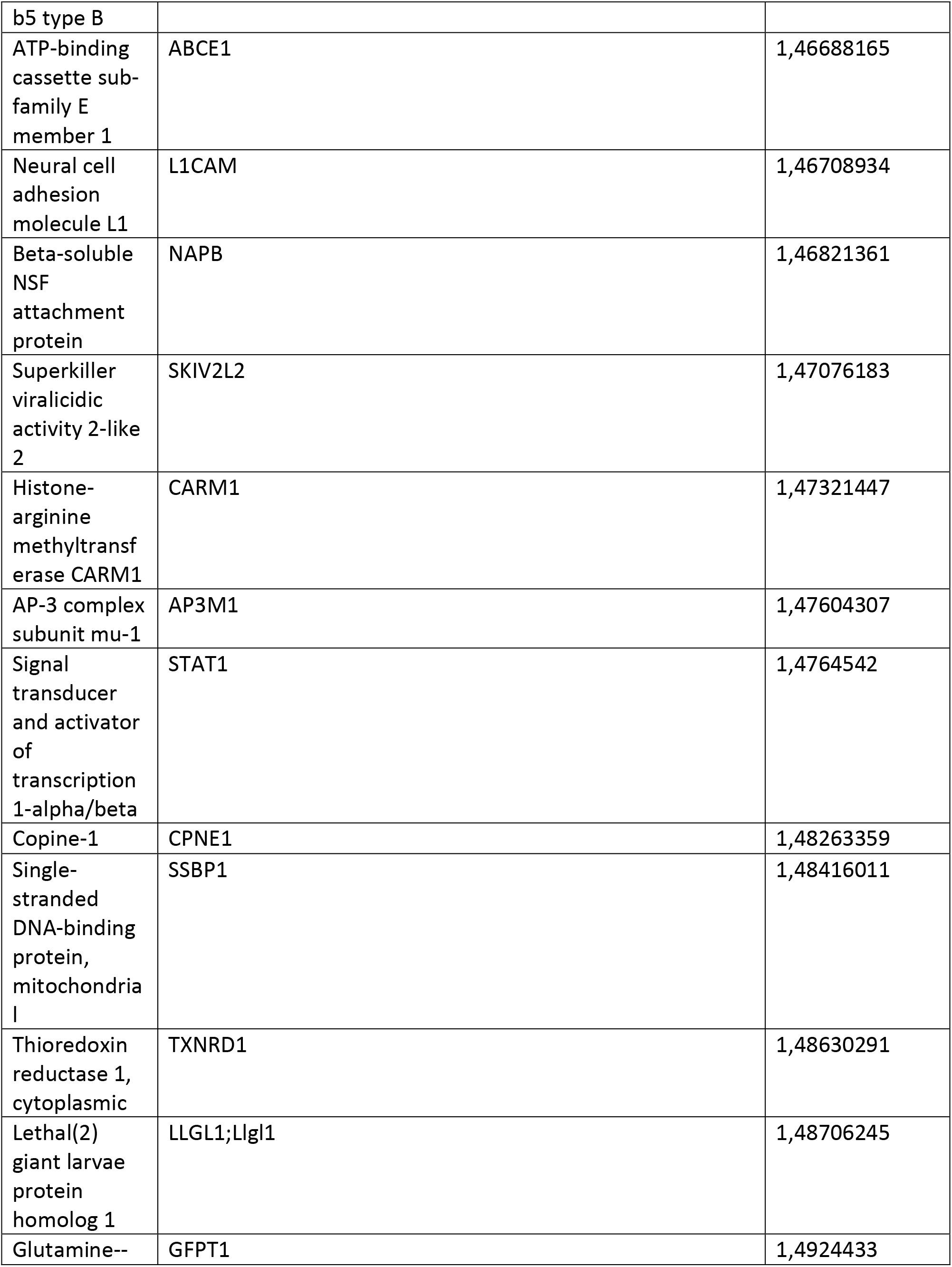
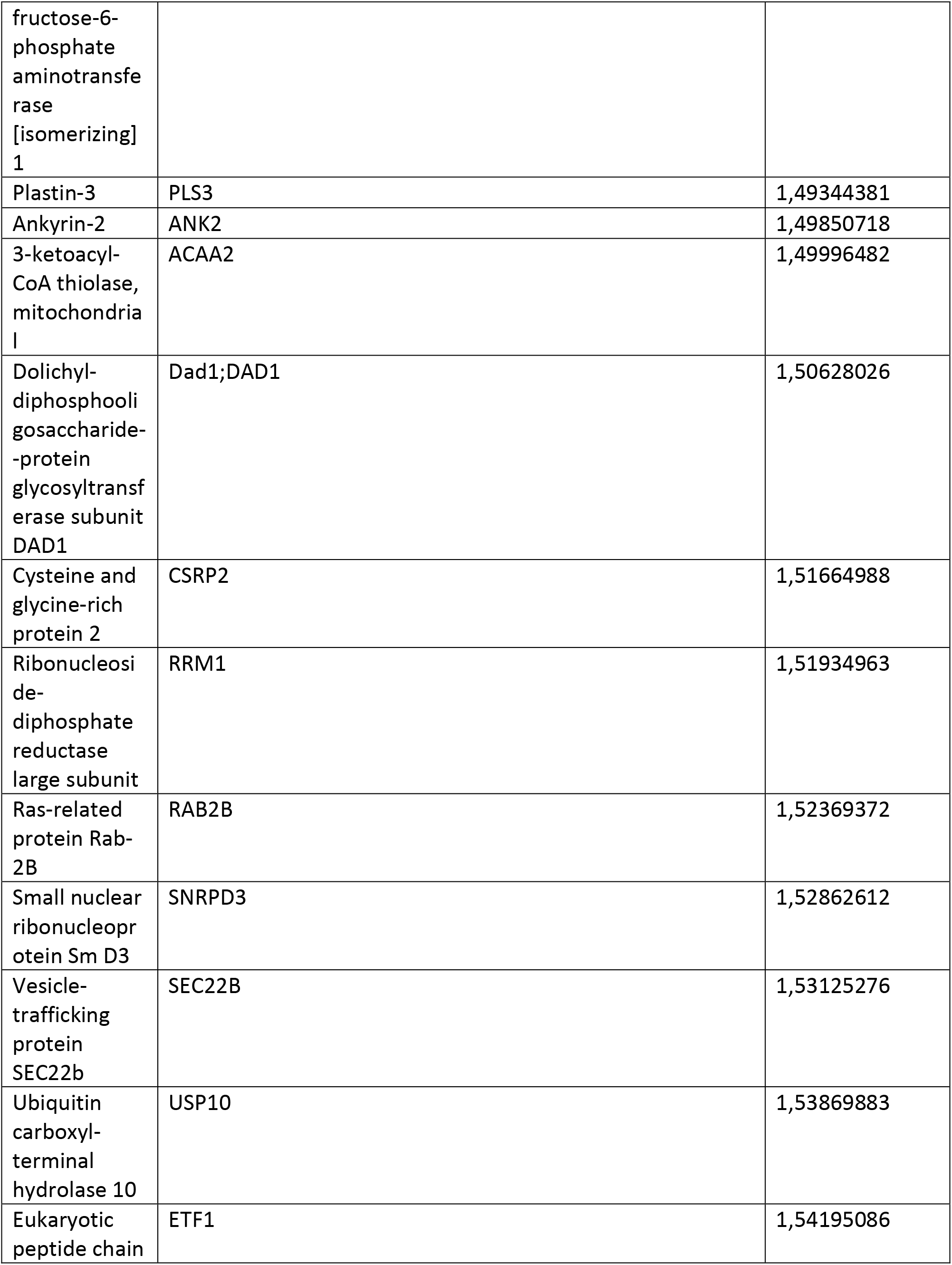
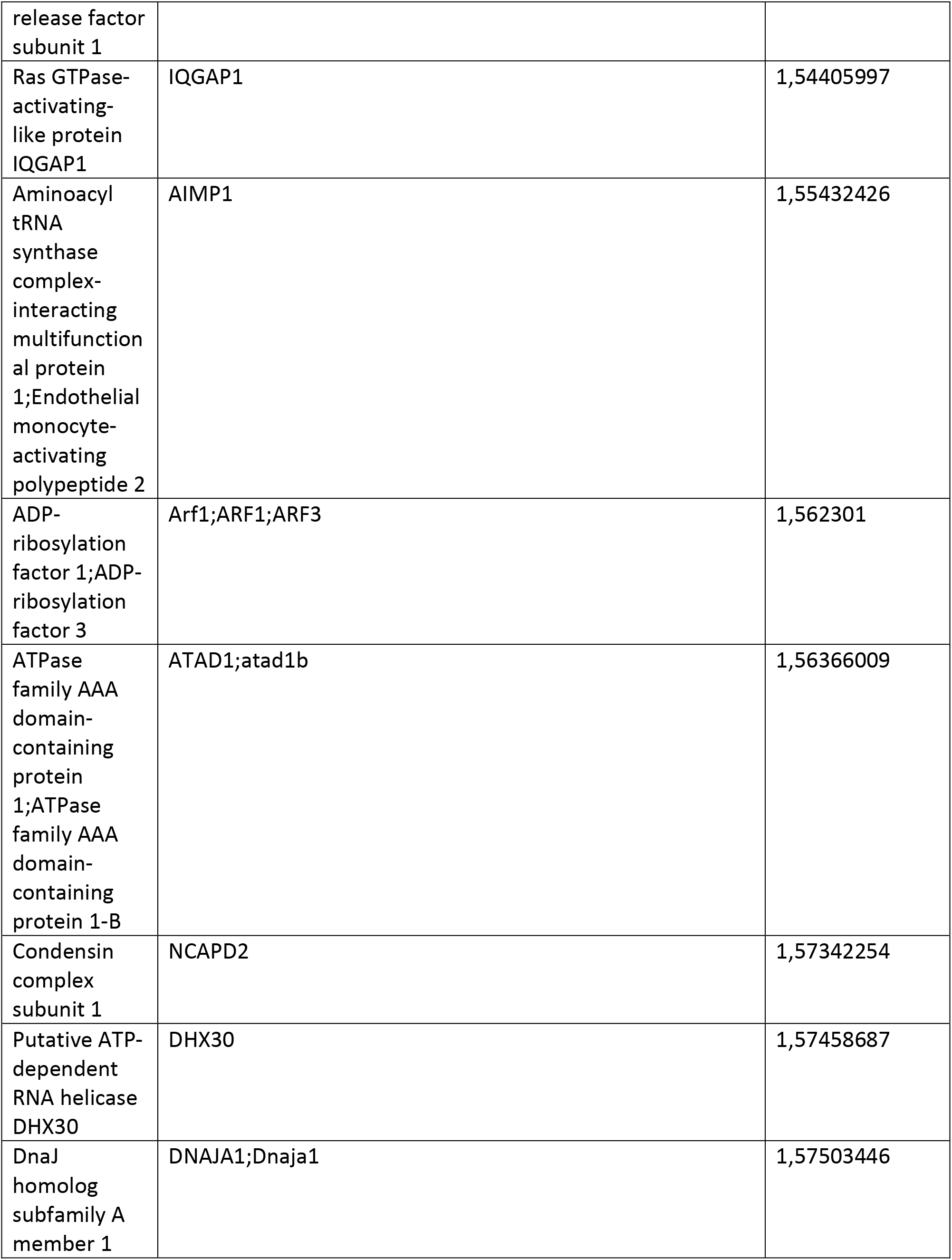
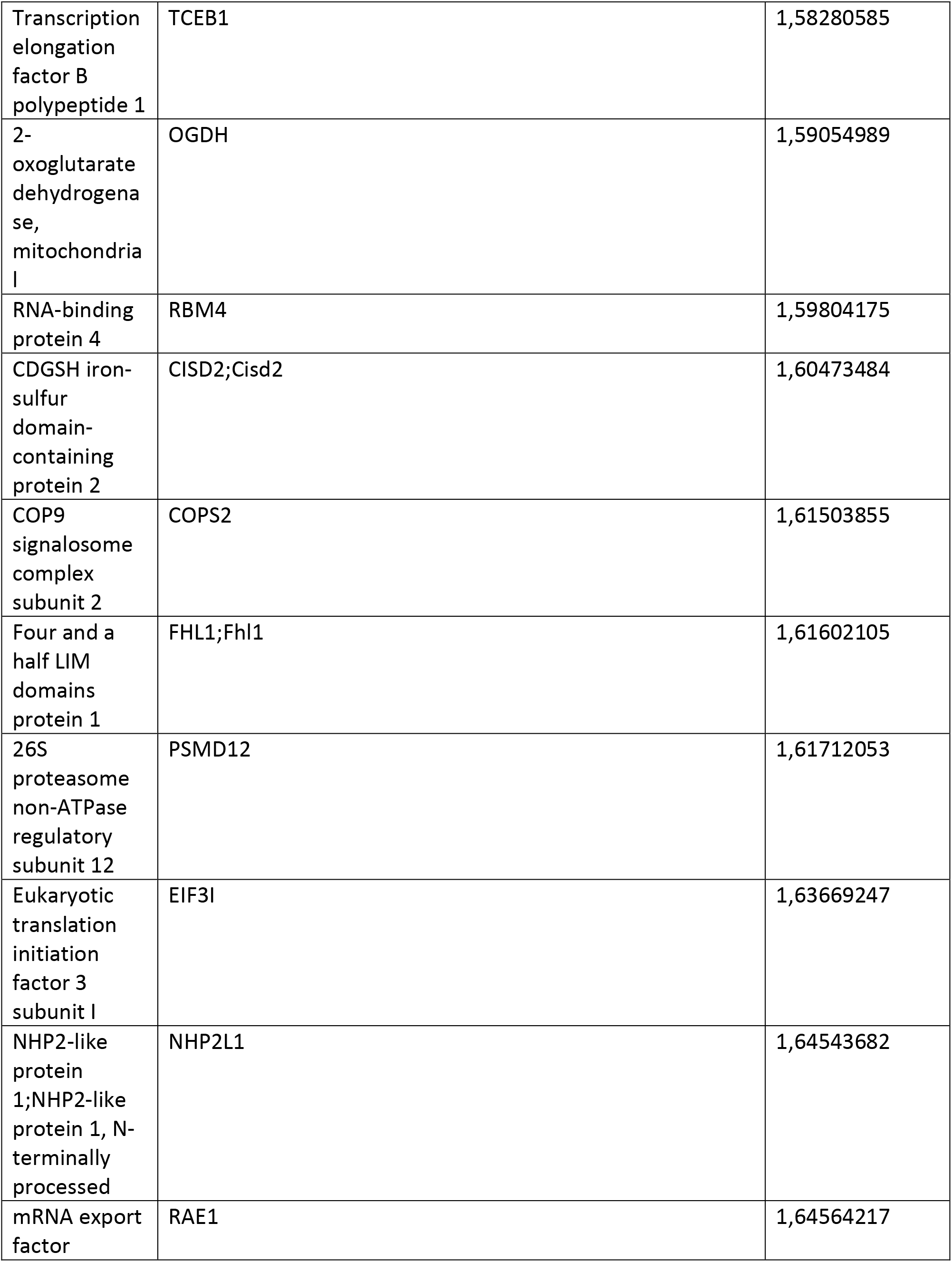
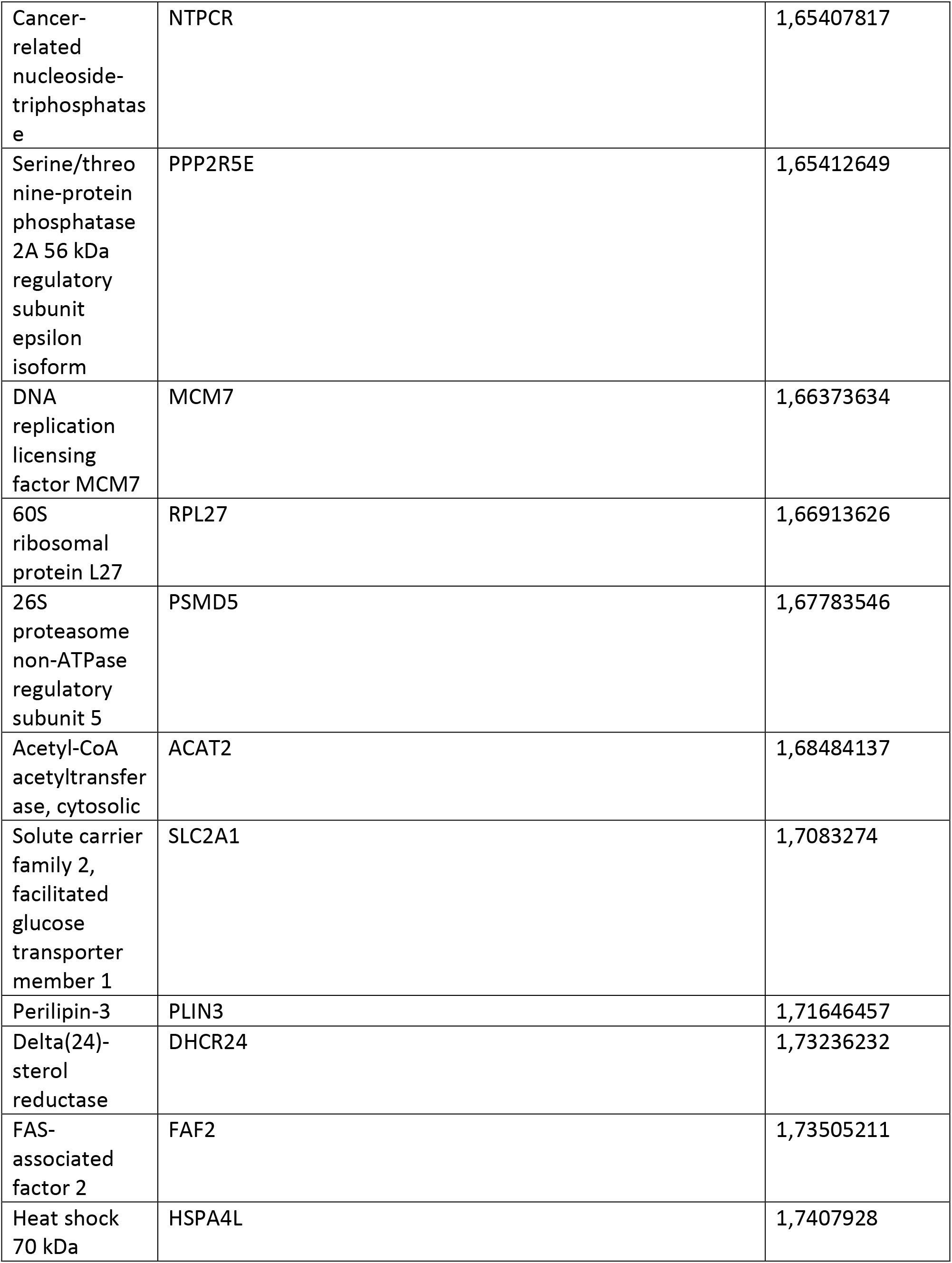
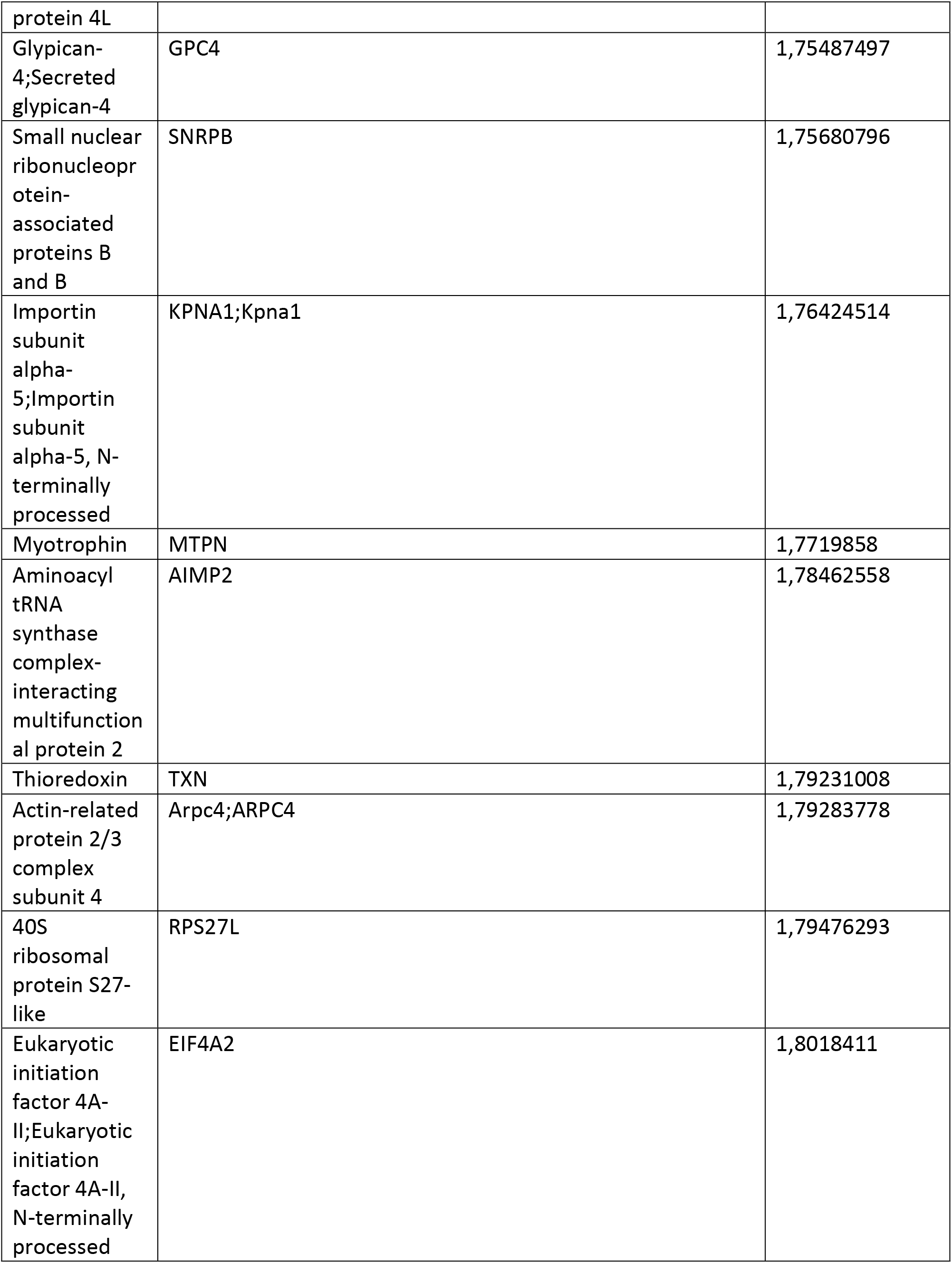
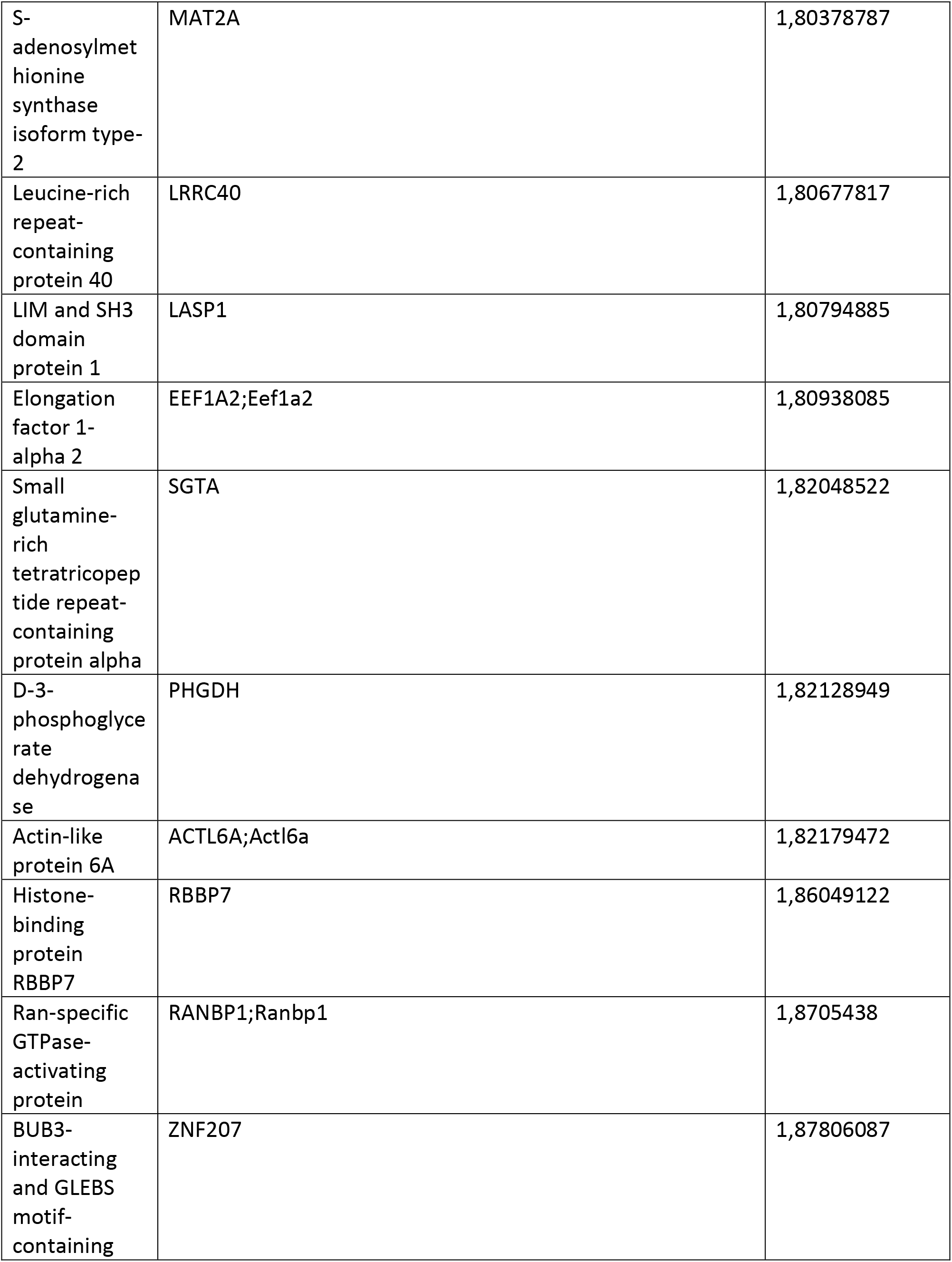
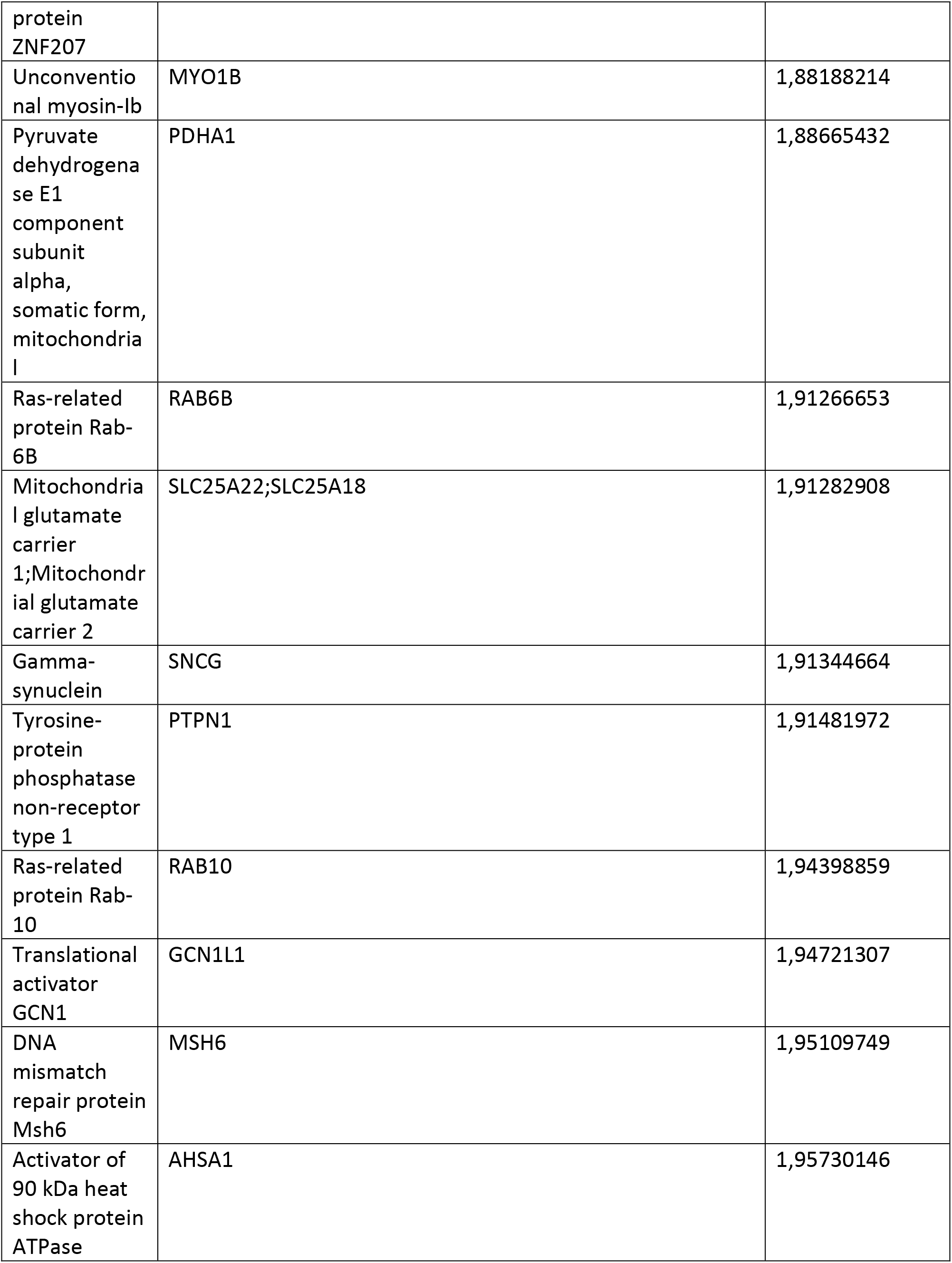
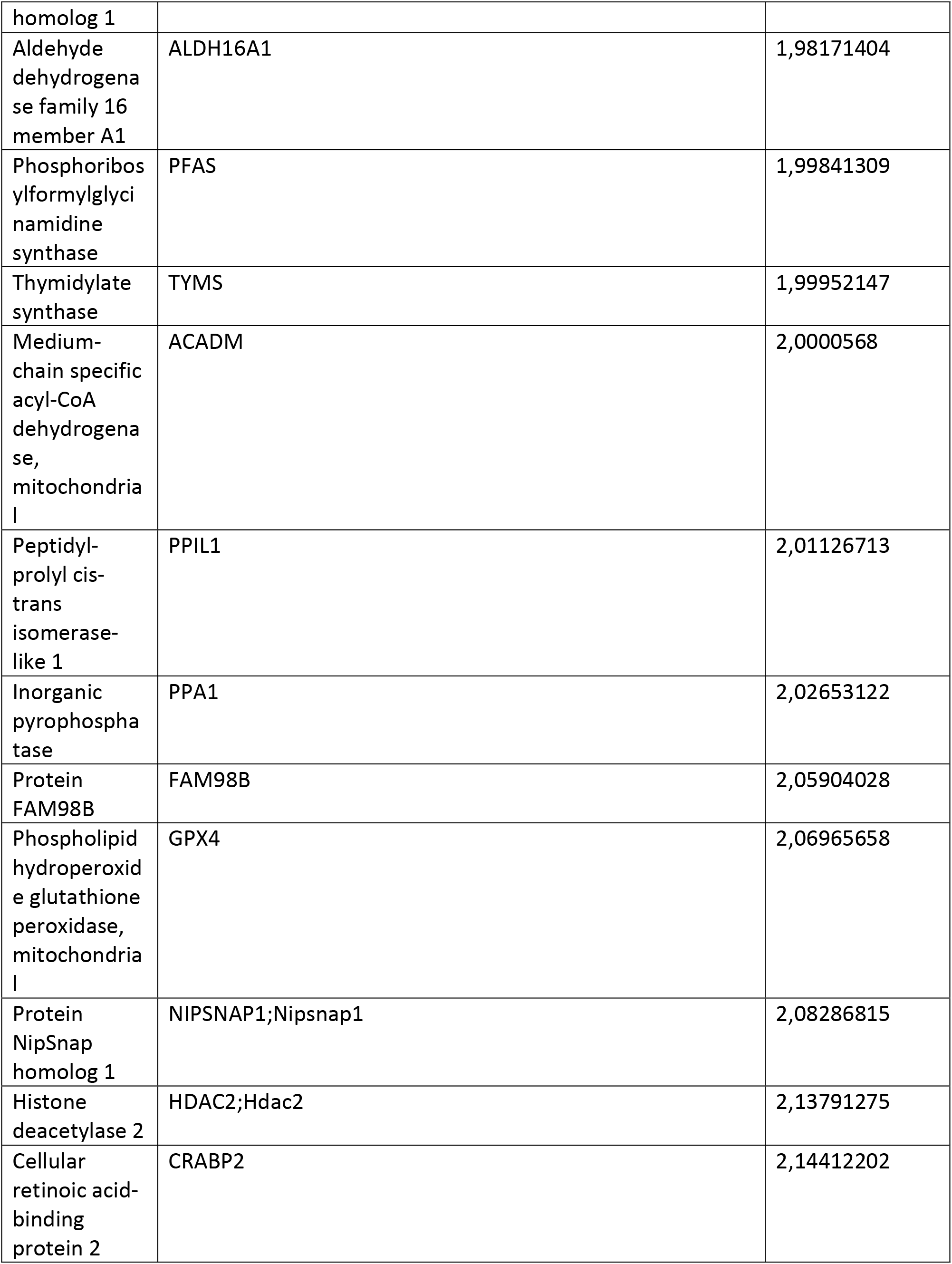
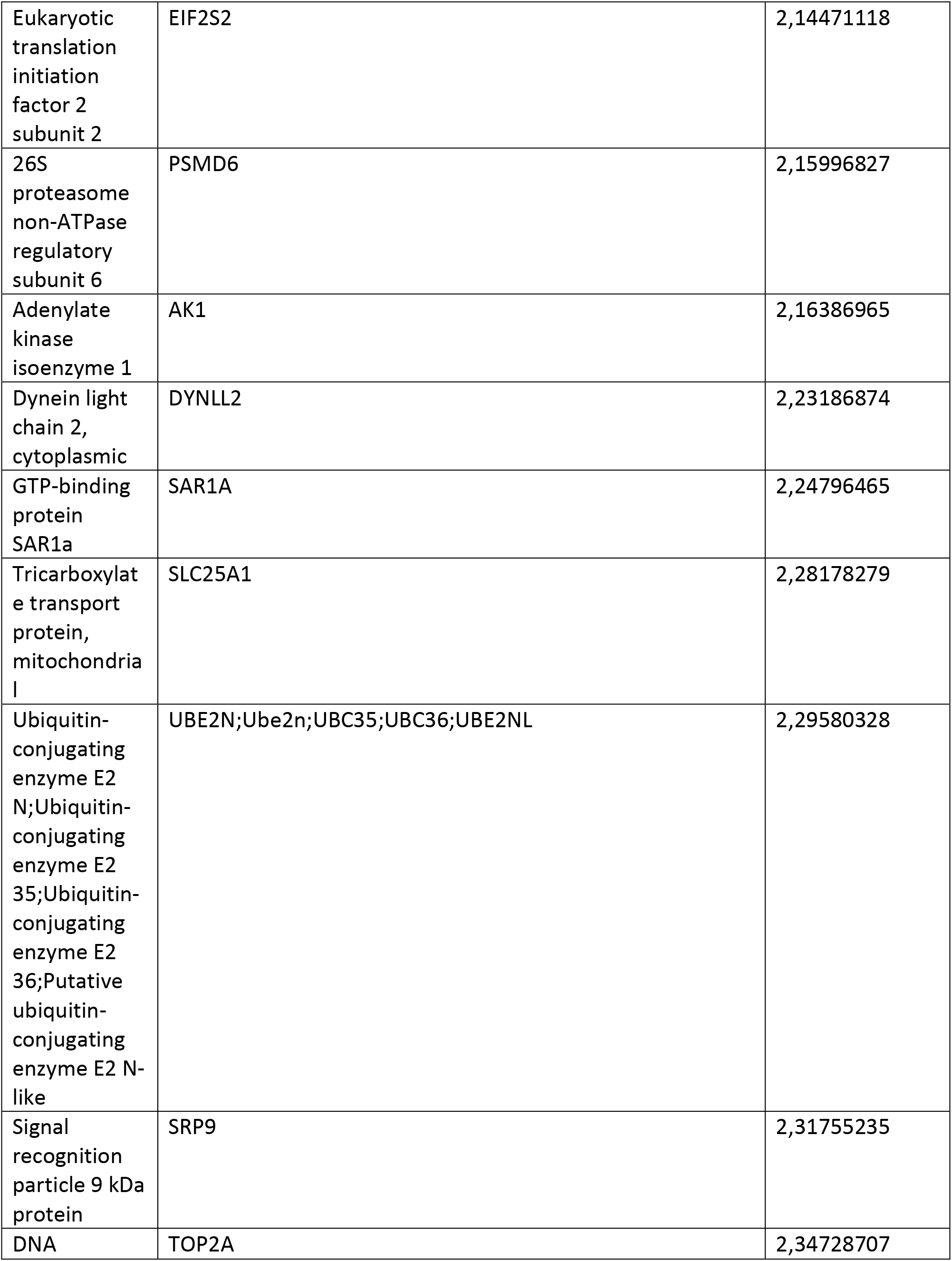
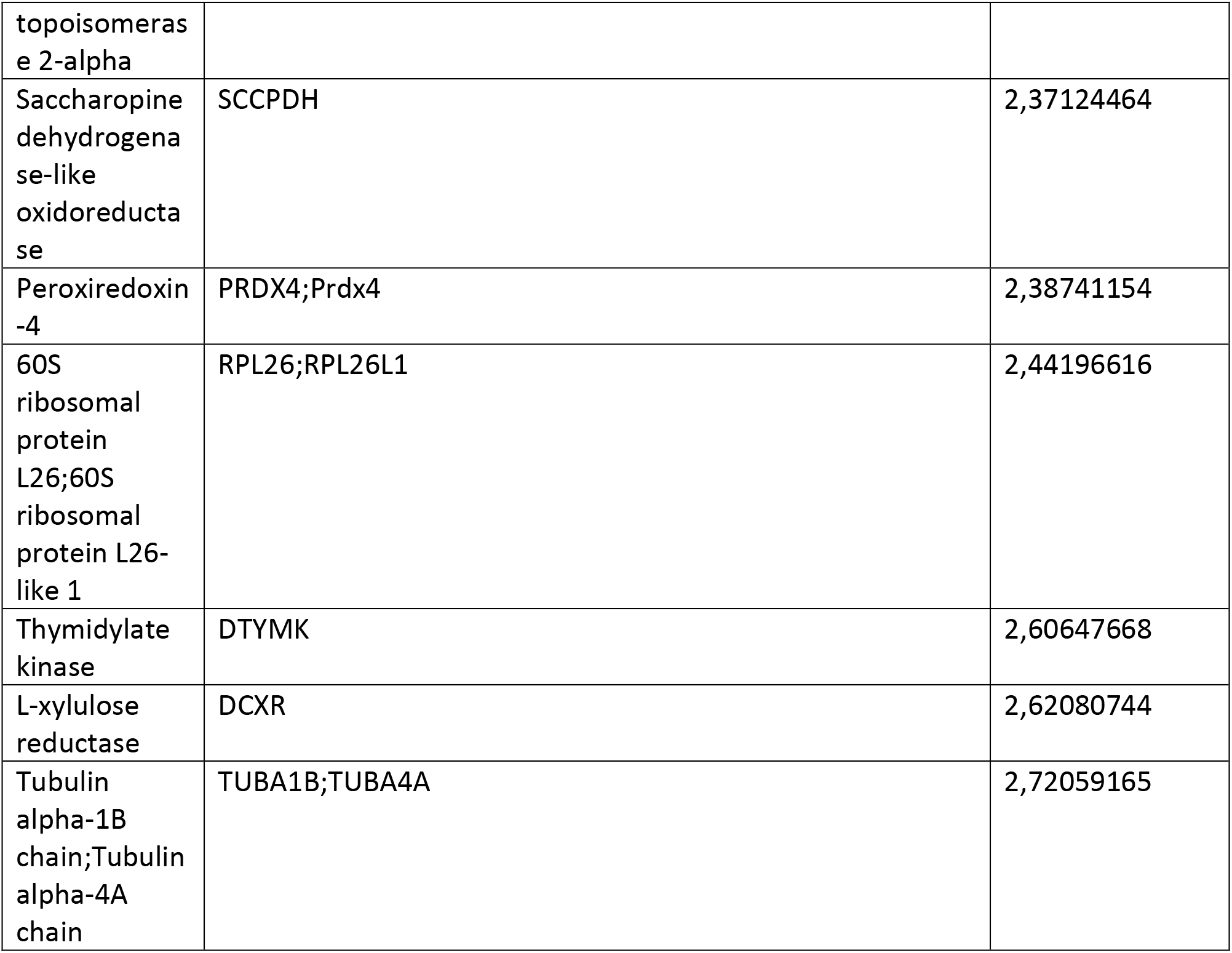
Differentially expressed proteins between pA53T and control neurons.

**Table S3:**
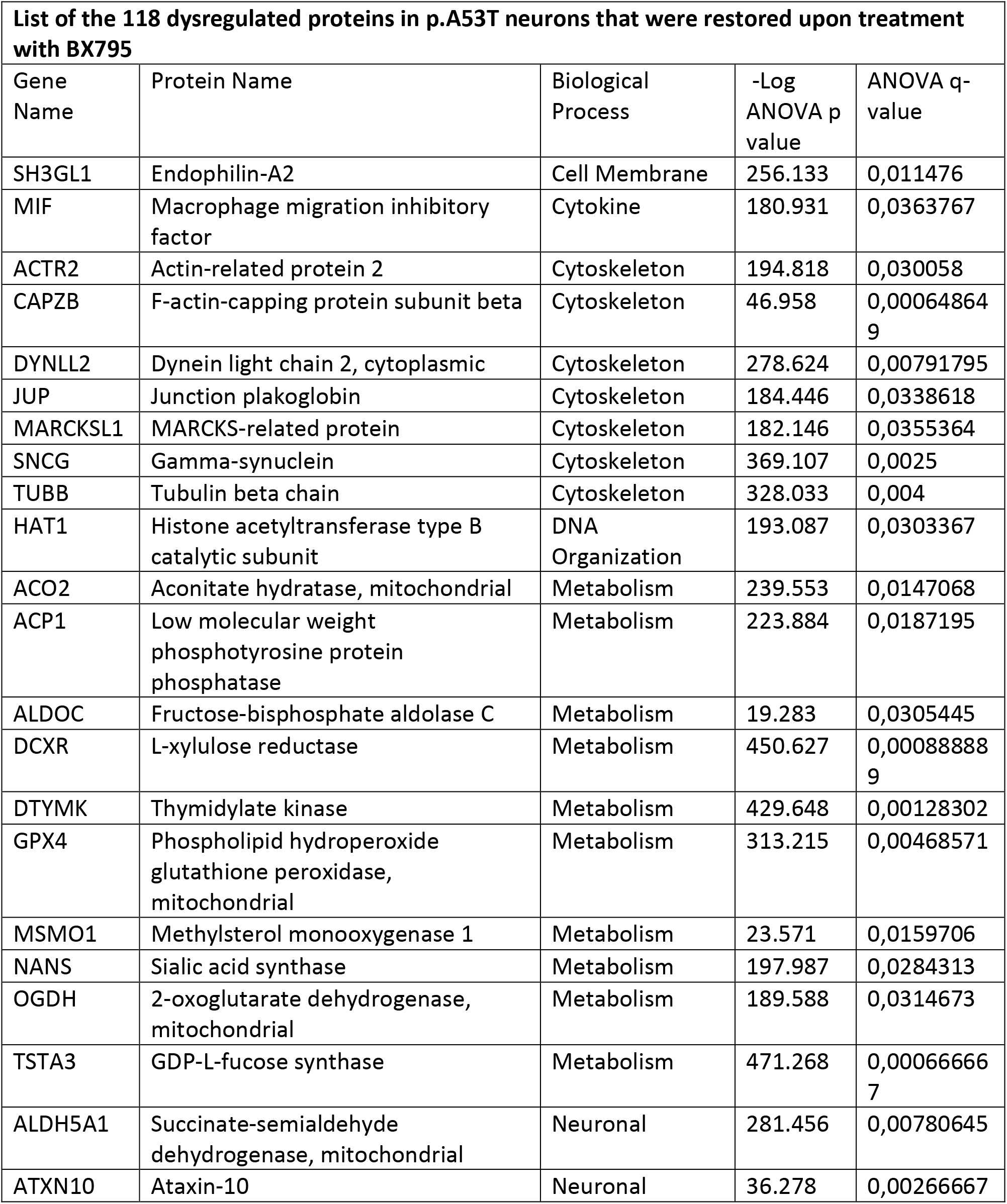
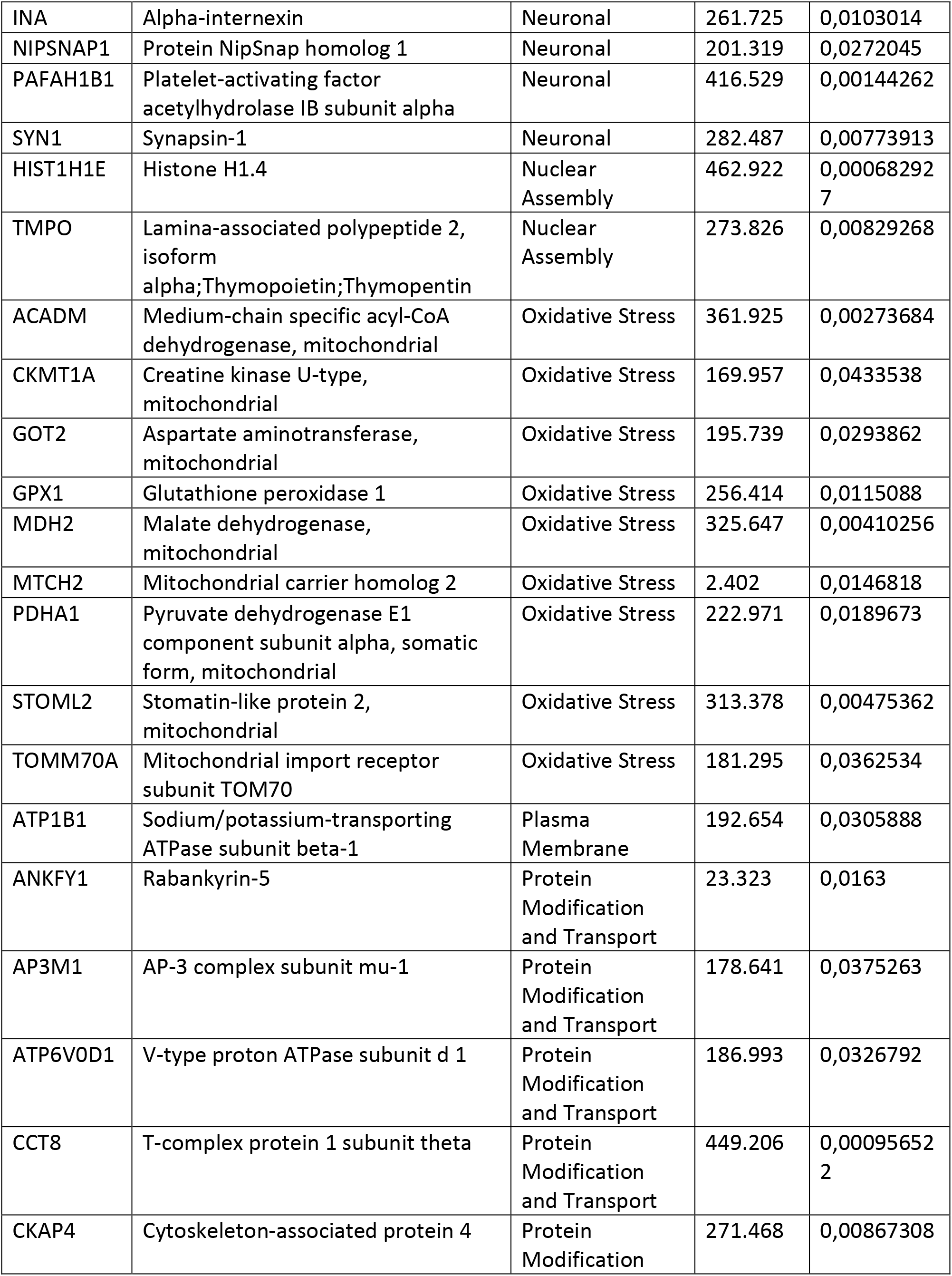
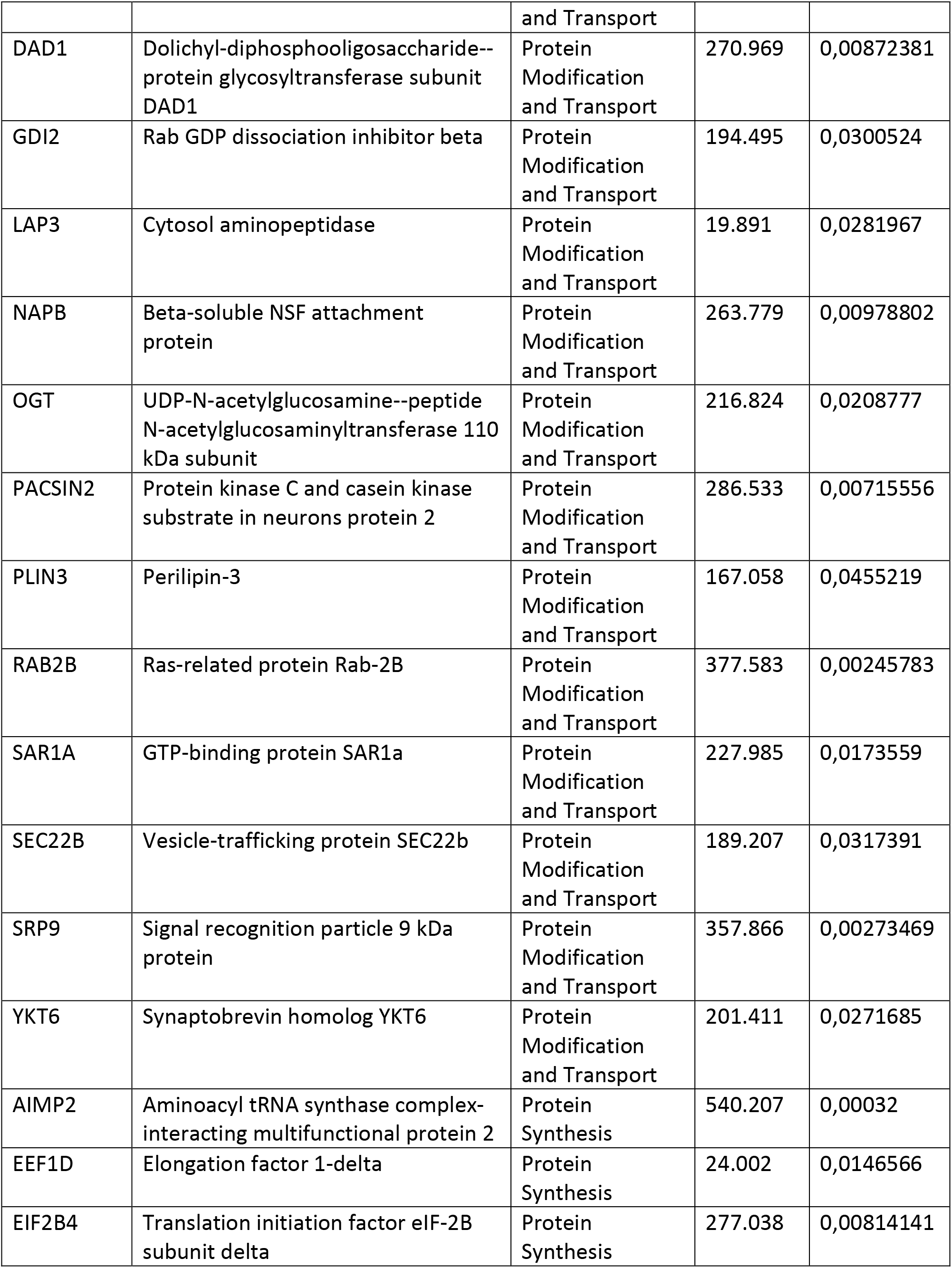
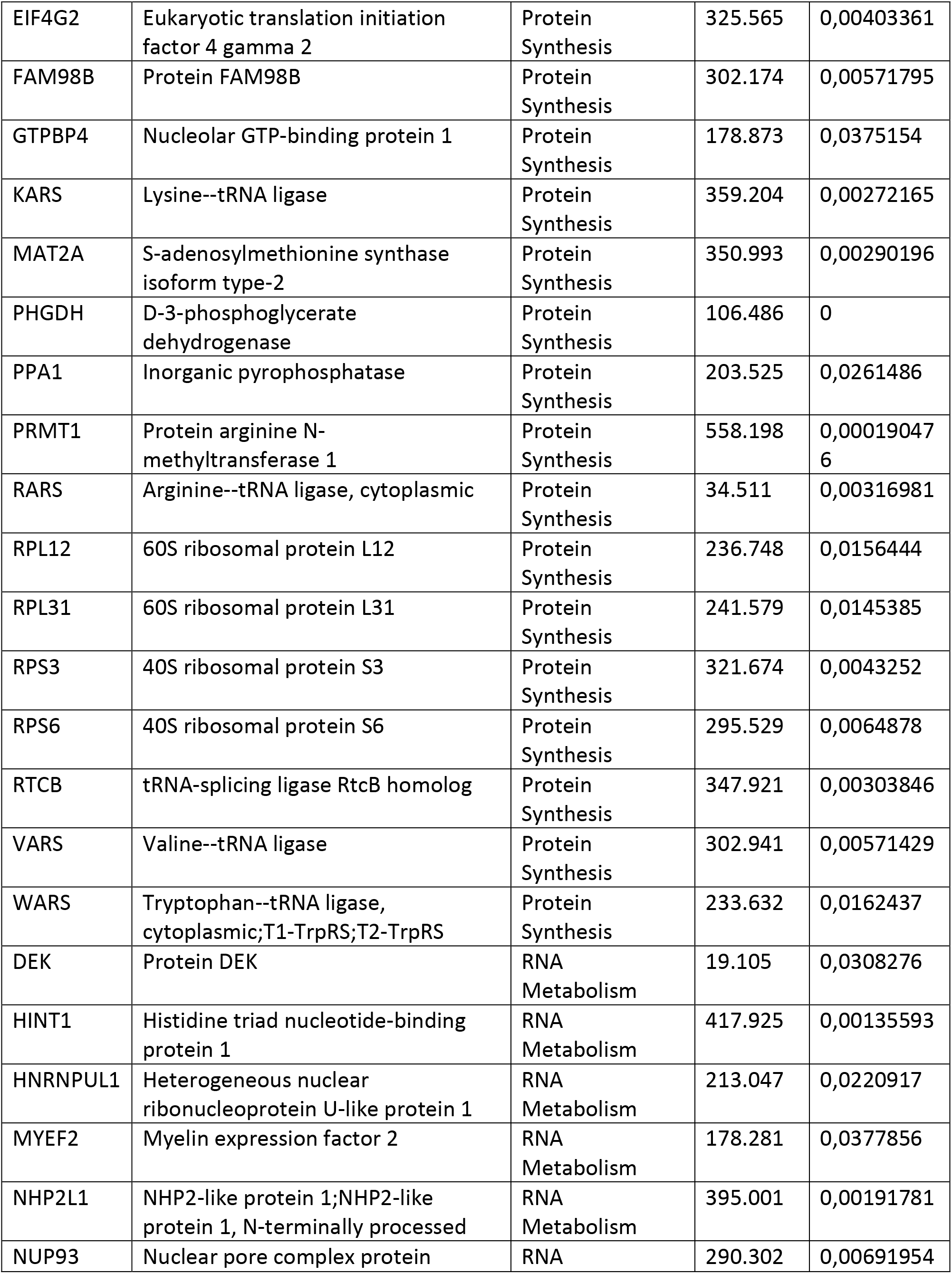
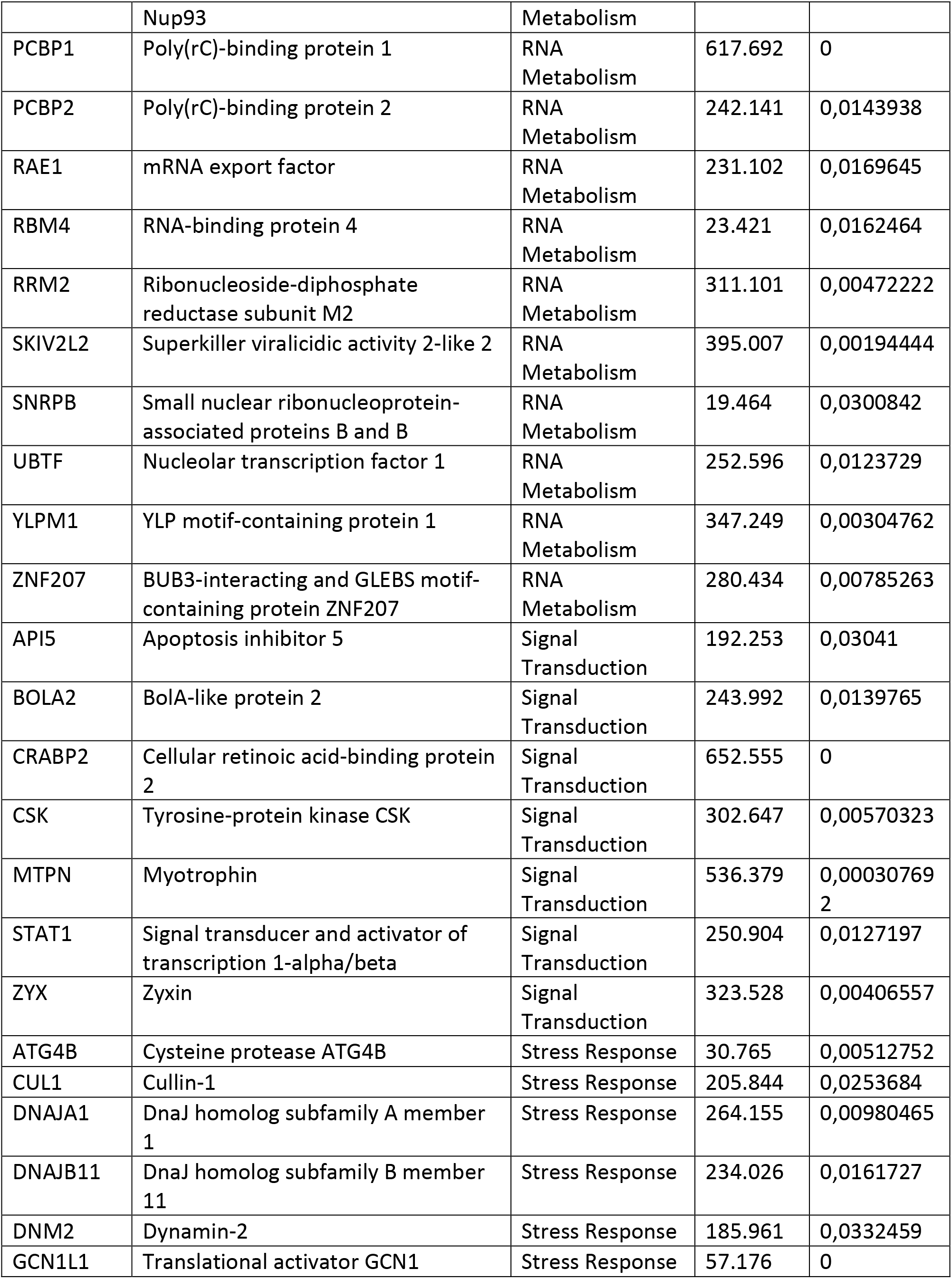
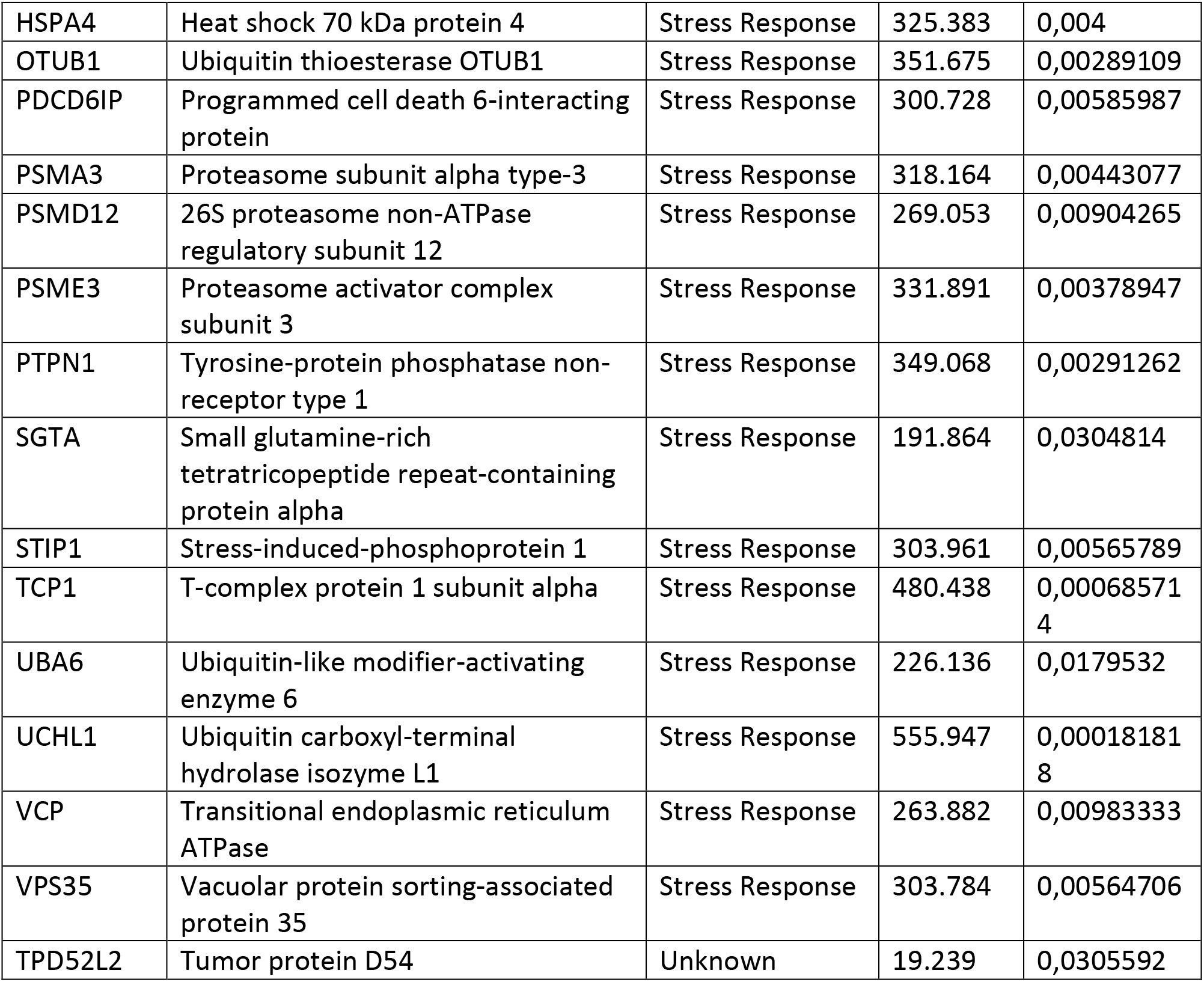
List of the 118 dysregulated proteins in p.A53T neurons that were restored upon treatment with BX795.

**Table S4:** GO analysis for cellular compartment between pA53T and control neurons.

|  |  |  |  |
| --- | --- | --- | --- |
| Cellular Compartment | No of genes | P-Value | Bonferroni |
| extracellular exosome | 299 | 1,6E-75 | 1,0E-72 |
| membrane | 204 | 2,8E-38 | 1,7E-35 |
| nucleoplasm | 230 | 7,9E-36 | 4,8E-33 |
| cytoplasm | 334 | 2,2E-32 | 1,3E-29 |
| mitochondrion | 118 | 1,6E-19 | 9,5E-17 |
| nucleus | 292 | 6,3E-15 | 3,9E-12 |
| nuclear pore | 22 | 6,7E-14 | 4,1E-11 |
| intracellular ribonucleoprotein complex | 29 | 7,5E-14 | 4,6E-11 |
| nucleosome | 23 | 2,4E-12 | 1,4E-9 |
| nuclear chromosome, telomeric region | 26 | 8,1E-12 | 4,9E-9 |
| nuclear nucleosome | 16 | 2,1E-11 | 1,3E-8 |
| nuclear envelope | 28 | 2,5E-11 | 1,5E-8 |
| proteasome complex | 18 | 2,8E-11 | 1,7E-8 |
| focal adhesion | 43 | 4,4E-10 | 2,7E-7 |
| eukaryotic translation initiation factor 3 complex | 10 | 2,1E-9 | 1,3E-6 |
| proteasome accessory complex | 9 | 5,6E-8 | 3,4E-5 |
| chaperonin-containing T-complex | 7 | 1,8E-7 | 1,1E-4 |
| nuclear membrane | 27 | 3,3E-7 | 2,0E-4 |
| cell body | 14 | 3,6E-7 | 2,2E-4 |
| proteasome regulatory particle | 7 | 9,3E-7 | 5,7E-4 |
| eukaryotic translation initiation factor 3 complex, eIF3m | 6 | 1,3E-6 | 7,8E-4 |
| axon cytoplasm | 10 | 3,40E-06 | 7,3E-5 |

**Table S5:**
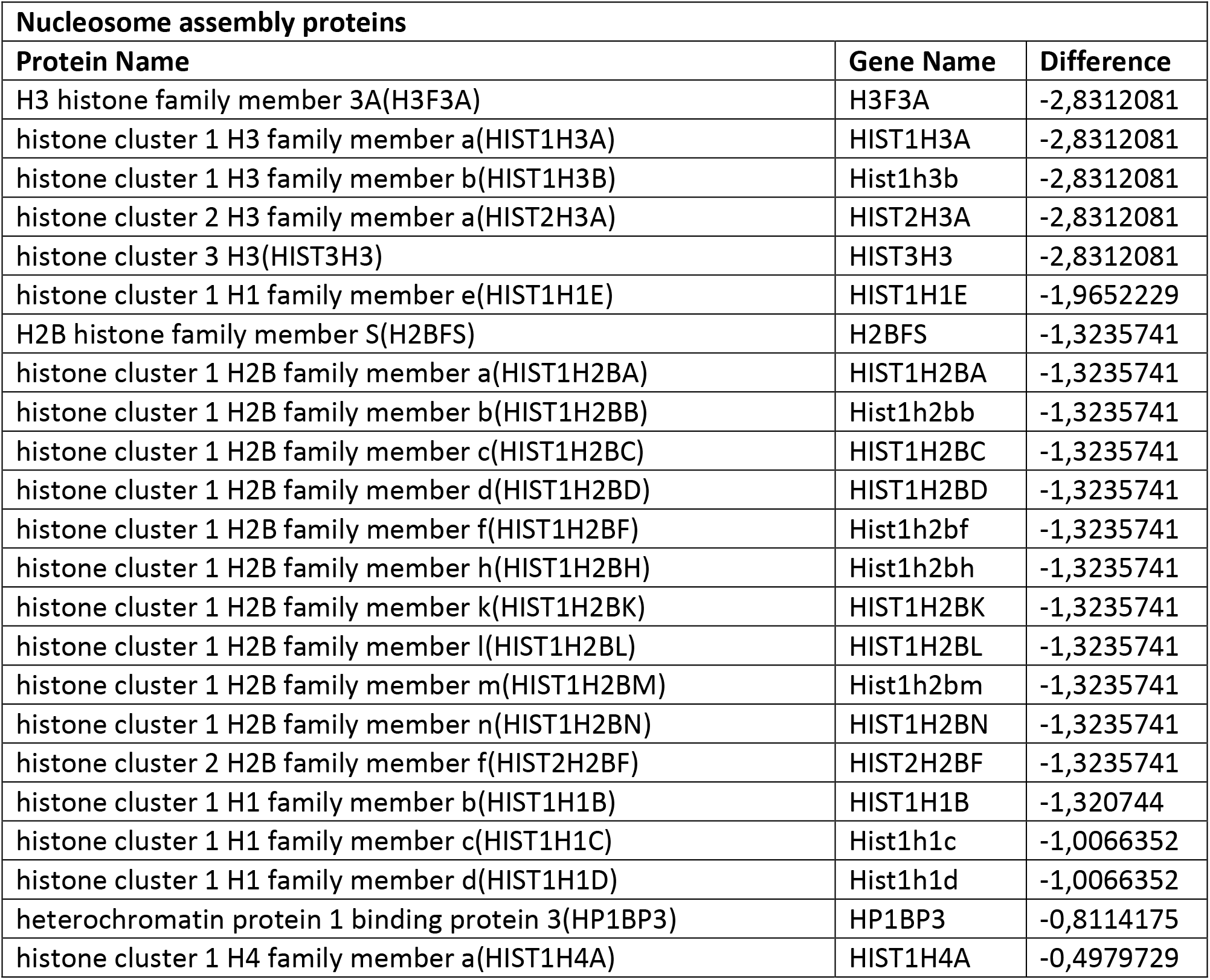
Nucleosome assembly proteins.

**Table S6:**
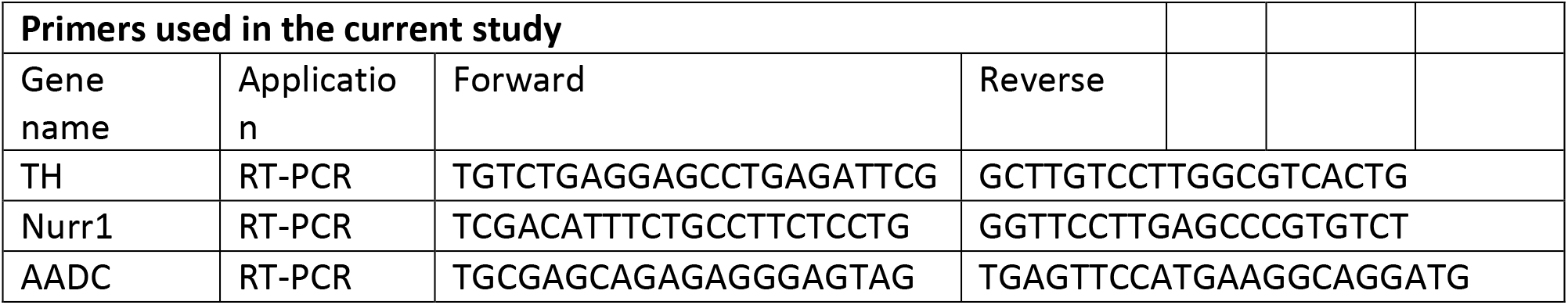
Primers used in the current study.

**Table S7:** Primary antibodies used in the current study.

| Primary antibodies used in the current study |  |  |  |  |
| --- | --- | --- | --- | --- |
| Name | Host | Dilution | Vendor | Catalog# |
| Anti-GAPDH | Mouse | 1/1000 | Santa Cruz Biotechnology | sc-365062 |
| Anti-beta actin | Mouse | 1/5000 | Abcam | ab8227 |
| Anti-MAP2 | Mouse | 1/200 | Merck-Millipore | MAB3418 |
| Anti-NESTIN | Rabbit | 1/200 | Merck-Millipore | ABD69 |
| Anti-α-Synuclein (αSyn) | Mouse | 1/500 | BD Biosciences | 610787 |
| Anti-phosphorylated α-Synuclein (Ser129) | Mouse | 1/10000 | WAKO | 015-25191 |
| Anti-TH | Rabbit | 1/500 | Merck-Millipore | AB152 |
| Anti-VGLUT1 | Mouse | 1/1000 | Merck-Millipore | MAB5502 |
| Anti-TUJ1 | Mouse | 1/1000 | Biolegend | 801202 |
| Anti-PAX6 | Mouse | 1/100 | DSHB | AB528427 |
| Anti-ki67 | Rabbit | 1/400 | Abcam | ab15580 |
| Anti-Phospho-S6 Ribosomal Protein (Ser235/236) | Rabbit | 1/1000 | Cell Signalling | 4858 |
| Anti-S6 Ribosomal Protein (5G10) | Rabbit | 1/1000 | Cell Signalling | 2217 |
| Anti-Phospho-mTOR (Ser2448) (D9C2) | Rabbit | 1/1000 | Cell Signalling | 5536 |
| Anti- mTOR (7C10) | Rabbit | 1/1000 | Cell Signalling | 2983 |
| Anti-Phospho-PRAS40 (Thr246) (C77D7) | Rabbit | 1/1000 | Cell Signalling | 2997 |
| Anti-PRAS40 (D23C7) | Rabbit | 1/1000 | Cell Signalling | 2691 |
| Anti-TBK1/NAK | Rabbit | 1/1000 | Cell Signalling | 3013 |
| Anti-Phospho-TBK1/NAK (Ser172) (D52C2) | Rabbit | 1/1000 | Cell Signalling | 5483 |
| Anti-Phospho-PDK1 (Ser241) | Rabbit | 1/1000 | Cell Signalling | 3061 |
| Anti-PDK1 (D37A7) | Rabbit | 1/1000 | Cell Signalling | 5662 |

